# Targeting tumour-intrinsic neural vulnerabilities of glioblastoma

**DOI:** 10.1101/2022.10.07.511321

**Authors:** Sohyon Lee, Tobias Weiss, Marcel Bühler, Julien Mena, Zuzanna Lottenbach, Rebekka Wegmann, Miaomiao Sun, Michel Bihl, Bartłomiej Augustynek, Sven Baumann, Sandra Goetze, Audrey van Drogen, Patrick Pedrioli, Daniel Kirschenbaum, Flavio Vasella, Elisabeth J. Rushing, Bernd Wollscheid, Matthias A. Hediger, Weller Michael, Berend Snijder

**Author notes:** These authors contributed equally (Sohyon Lee, Tobias Weiss). These authors contributed equally (Weller Michael, Berend Snijder).

## Abstract

Glioblastoma is the most common yet deadliest primary brain cancer^1^. The neural behavior of glioblastoma, including the formation of synaptic circuitry and tumour microtubes, is increasingly understood to be pivotal for disease manifestation^2–9^. Nonetheless, the few approved treatments for glioblastoma target its oncological nature, while its neural vulnerabilities remain incompletely mapped and clinically unexploited. Here, we systematically survey the neural molecular dependencies and cellular heterogeneity across glioblastoma patients and diverse model systems. In 27 surgical patient samples, we identify cancer cell morphologies indicative of poor prognosis, and discover repurposable neuroactive drugs with anti-glioblastoma efficacy by image-based drug screening. Glioblastoma cells exhibit functional dependencies on highly expressed neuroactive drug targets, while interpretable molecular machine learning (COSTAR) reveals their downstream convergence on AP-1-driven tumour suppression. This drug-target connectivity signature is confirmed by accurate *in silico* drug screening on >1 million compounds, as well as by multi-omic profiling of glioblastoma drug responses. Thus, Ca^2+^-driven AP-1 pathway induction represents a tumour-intrinsic vulnerability at the intersection of oncogenesis and neural activity-dependent signaling. Opportunities for clinical translation of this neural vulnerability are epitomized by the antidepressant Vortioxetine synergizing with current standard of care treatments *in vivo*. Together, the results presented here provide a mechanistic foundation and conceptual framework for the treatment of glioblastoma based on its neural origins.

## Introduction

Glioblastoma is a deadly brain cancer with limited treatment options, shaped by heterogeneous developmental programs, genetic drivers, and tumour microenvironments ^10–14^. Despite an increasing understanding of this heterogeneity, the alkylating agent Temozolomide (TMZ), which prolongs median survival from 12 to 15 months, remains the only first-line drug approved for glioblastoma ^15–17^. Targeted therapies have been largely unsuccessful, in part due to the blood-brain barrier (BBB) limiting tumour accessibility, the presence of treatment-resistant glioblastoma stem cells (GSCs), and the lack of clinically predictive models ^18–23^. Systemically addressing these therapeutic roadblocks is an urgent clinical need.

An emerging paradigm is to consider glioblastoma in the context of the nervous system. Single-cell RNA sequencing (scRNA-Seq) and lineage tracing studies of glioblastoma have identified stemness signatures resembling neural development ^7,12,13,24–29^. At the brain-tumour interface, synaptic integration of cancer cells into neural circuits regulates tumour growth ^3,5,6,9^. Within the tumour, the extension of microtubes akin to neuronal protrusions promotes the formation of treatment-resistant invasive networks ^2,4,8^. Furthermore, modulating specific neurotransmitter or other secretory pathways in the tumour microenvironment impairs glioblastoma metabolism and survival ^3,30–32^. Such neural aspects of glioblastoma offer new clinically-targetable vulnerabilities that could be targeted by repurposing approved “neuroactive” drugs (NADs). Neuroactive drugs can cross the BBB and are routinely prescribed for indications such as psychiatric or neurodegenerative diseases. Yet, as neuroactive drugs are originally developed to modulate the nervous system, their anti-cancer activity in glioblastoma patients is largely unknown.

Several key questions arise. First, how does neural intratumour heterogeneity across glioblastoma patients relate to disease course and response to therapy? Second, are there tumour-intrinsic neural vulnerabilities that are therapeutically targetable? Third, if so, which molecular dependencies and associated pathways are involved?

Here, we find morphological and neural stemness features across glioblastomas that relate to disease prognosis and drug response. Using pharmacoscopy (PCY), an *ex vivo* imaging platform ^33–35^ that captures patient and tumour complexity, we screen repurposable neuroactive drugs and identify a set with potent anti-glioblastoma activity. Top neuroactive drugs work consistently across patient samples and particularly target GSCs with neural morphologies associated with invasion and poor prognosis. These top drugs are validated across multiple glioblastoma model systems including patient-derived cultures and orthotopic xenograft mouse models. Integration of anti-glioblastoma response with multiplexed RNA-Seq, reverse genetic screening, and machine learning of drug-target networks reveals convergence of neuroactive drugs with anti-glioblastoma activity on AP-1 and BTG gene families. In a neural context, AP-1 transcription factors, including JUN and FOS, are immediate early genes (IEGs) induced in response to neural activity or insult, while BTG1/2 are known tumour suppressors ^36–39^. Using this convergent AP-1/BTG connectivity signature, we predict and validate new candidate drugs across >1 million compounds *in silico*. The antidepressant Vortioxetine is the top PCY-hit and inducer of the AP-1/BTG signature across diverse experimental model systems, synergizing with both first- and second-line glioblastoma therapies *in vivo*. Our study identifies clinically-actionable neuroactive drugs for the treatment of glioblastoma converging on a gene regulatory network involved in cell proliferation and neural activity.

## Main

### Capturing the phenotypic single-cell heterogeneity of glioblastoma

Glioblastoma cells adopt unique cellular morphologies and stemness properties to integrate and survive in the brain ^2,4,8,13,40^. To comprehensively profile the morphological and molecular heterogeneity between and within glioblastoma patients, we investigated surgically-resected material from a clinically-annotated cohort of 27 glioblastoma patients (prospective cohort; Fig. 1a, Extended Data Fig. 1a and Supplementary Table 1). We quantified cell type composition and morphology by high-content confocal imaging of freshly dissociated surgical samples cultured for two days *ex vivo* (n=27 patients), as well as in patient-matched tissue sections *in situ* (n=10 patients). In parallel, we measured somatic genetic alterations by targeted next-generation sequencing (NGS; n=27 patients; Supplementary Table 2) and single-cell transcriptomes by scRNA-Seq (n=4 patients).

**Fig. 1:**
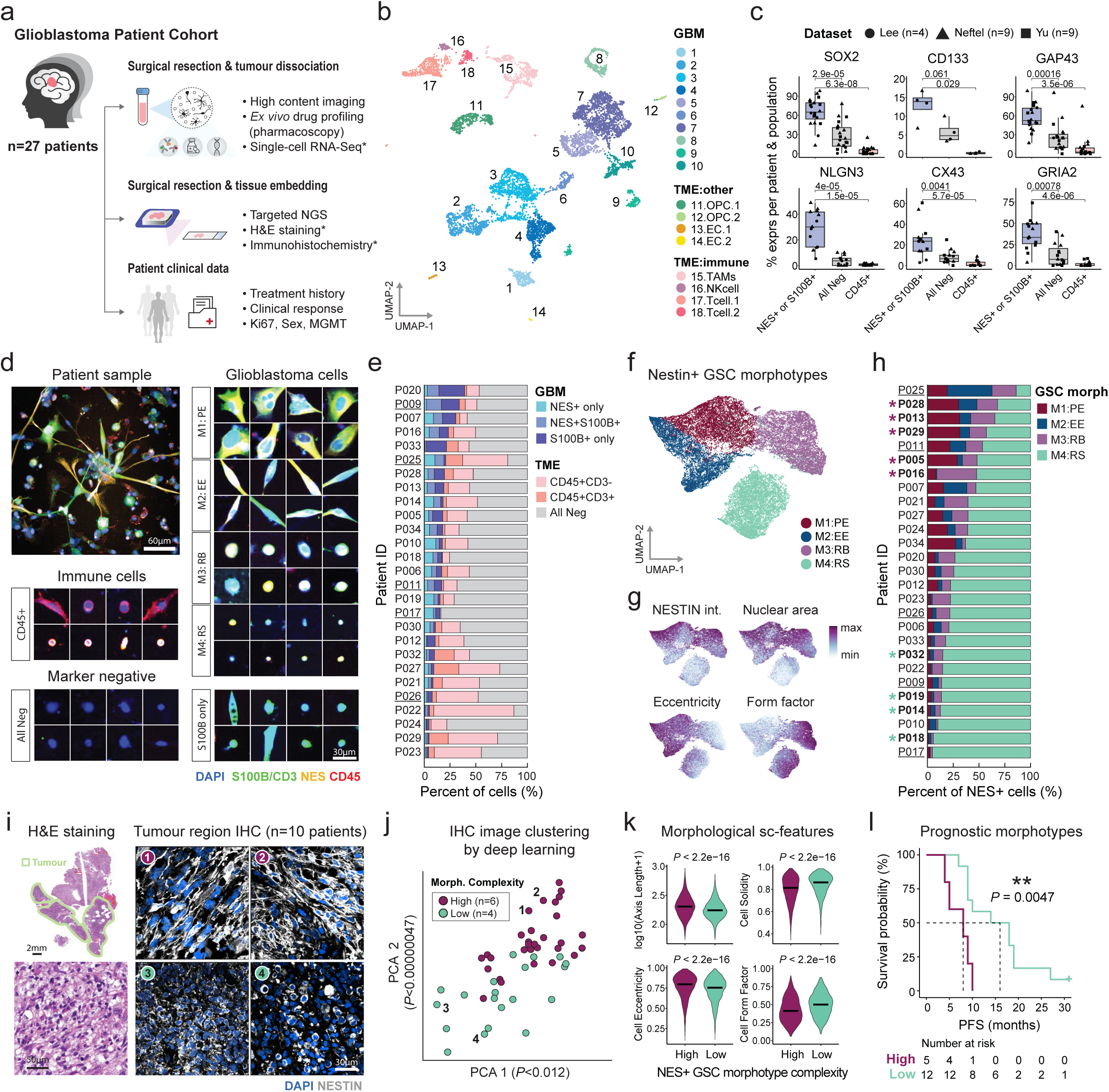
Neural intratumour heterogeneity across glioblastoma patients relates to disease prognosis. **a**, Prospective glioblastoma patient cohort (n=27 patients) and associated experiments. Asterisks (*) indicate assays performed with a subset of patients. **b,** UMAP projection of 7684 single-cell transcriptomes from four glioblastoma patient samples (P007, P011, P012, P013) colored by cluster-id (see methods and *Supplementary Fig.S1*). TME, tumour microenvironment; OPC, oligodendrocyte precursor cells; EC, endothelial cell; TAM, tumour-associated macrophage; NK, natural killer cell. **c,** Percent of cells expressing key marker genes (y-axis) per subpopulation (x-axis) across 22 glioblastoma patient samples (data points). Data from 3 scRNA-Seq datasets (data point shape; Lee *et al.*, this study; n=4 patients; Neftel *et al.*, n=9 patients; Yu *et al.*, n=9 patients) (see also *Extended Data* Fig. 1g). *P*-values calculated from a two-sided Wilcoxon rank sum test. Boxplots show 25th–75th percentiles with a line at the median; whiskers extend to 1.5 times the interquartile range. **d,** Compositional and morphological diversity of cells from dissociated glioblastoma patient samples captured by high-content *ex vivo* imaging. Glioblastoma ([NES+ or S100B+] and CD45-), immune (CD45+ and NES- and S100B-), and marker negative (NES- and S100B- and CD45-) cells are shown, as well as Nestin+ glioblastoma stem cell (GSC) morphotypes (M1:PE, polygonal cell with extensions; M2:EE, elongated cell with extensions; M3:RB, round big cells; M4:RS, round small cells). **e,** Cellular composition across the prospective glioblastoma cohort (n=27 patients). **f-g,** UMAP projection of the morphological CNN feature space of 84,180 Nestin+ GSCs (up to n=1000 cells per morphotype and patient; n=27 patients). Colored by **f,** assigned Nestin+ GSC morphotype (M1-M4); **g,** local median of selected single-cell features. Nestin Int.; Nestin expression measured by immunofluorescence. **h,** Nestin+ GSC morphotype composition across the prospective glioblastoma cohort (n=27 patients). Asterisks (*) indicate samples that were also profiled by immunohistochemistry (IHC) of patient-matched tissue sections. Red and green * indicate patients with high or low GSC morphotype complexity, respectively. **e,h,** Underlines indicate recurrent glioblastoma patient samples. **i,** Example images of glioblastoma patient tissue sections stained by H&E and IHC (DAPI, Nestin). H&E stained tissue section of patient P016 with tumour regions marked in green (top left) and zoom in (bottom left); Example IHC staining of tumour regions from patients with high *ex vivo* GSC morphotype complexity (top middle and right; 1, P016; 2, P040) and low *ex vivo* GSC morphotype complexity (bottom middle and right; 3, P014; 4, P019). **j,** Principal component analysis (PCA) of unsupervised deep learning-derived features for 50 multicellular IHC images (n=5 images/patient; n=10 patients). P-values indicate the significance of the differences in the corresponding PCs between images from patients with high (n=30 images; red dots) and low (n=20 images; green dots) *ex vivo* GSC morphotype complexity by Wilcoxon rank-sum test. Labeled numbers correspond to patient images in Fig. 1i. **k,** Violin plots comparing *in situ* morphological single-cell features (sc-features) of Nestin+ cells (n=400 randomly sampled cells/patient) between high (n=6 patients) and low (n=4 patients) morphotype complexity groups. Line denotes median and P-values based on Wilcoxon rank test. **l,** Kaplan-Meier curves of progression-free survival (PFS) in newly diagnosed glioblastoma patients (n=17 patients) stratified by M1-M3 morphotype abundance (high, low) within Nestin+ GSCs. Survival curves are compared using the log-rank (Mantel-Cox) test. Tick mark indicates ongoing response.

We mapped the single-cell landscape of glioblastoma patient samples captured by both scRNA-seq and high-content *ex vivo* imaging. Across technologies, glioblastoma cells were placed along a neural stemness gradient against the neural progenitor marker Nestin and the mature astrocytic marker S100B (Fig. 1b-e, Extended Data Fig. 1b-g, and Supplementary Fig. S1). Concordant with previous literature using Nestin/S100B as glioblastoma markers ^2,5,7,20,41^, analysis of 25,510 single cell transcriptomes across three independent scRNA-Seq datasets revealed the highest co-expression of markers associated with malignancy (e.g. SOX2, CD133, EGFR, Ki67) and neural properties of glioblastoma (e.g. GAP43, NLGN3, CX43, GRIA2; Fig. 1c and Extended Data Fig. 1e-g) in the [Nestin+ or S100B+ and CD45-] glioblastoma cell definition (n=22 glioblastoma patients; Lee *et al.*, this study; n=4 patients; Neftel *et al.^13^*, n=9 patients; Yu *et al.^29^*, n=9 patients). Expression profiles of glioblastoma cells displayed high inter-patient heterogeneity within the Nestin/S100B spectrum, and were distinct from CD45+ immune cells present in the tumour microenvironment (TME; Fig. 1b,c, Extended Data Fig. 1c,d, and Supplementary Fig.S1). Cell-type specific enrichment analysis of gene modules enriched in Nestin/S100B/CD45-negative cells (‘All Neg’) confirmed the presence of additional TME cell types, including CD45-low tumour-associated macrophages/microglia, fibroblasts, and stromal cells (Extended Data Fig. 1k,l). By immunofluorescence (IF), we quantified cell type composition and morphology for over 100 million imaged patient cells. This confirmed on average >90% *ex vivo* viability of glioblastoma cells across the prospective cohort (Extended Data Fig. 1h-j and *Methods*) and revealed a high degree of inter- and intra-tumour heterogeneity: Across patients, glioblastoma cells ranged from 4-39%, immune cells from 1-82%, and all marker negative TME cells 13-84% (Fig. 1d,e). Imaging glioblastoma sample composition underscored the molecular tumour heterogeneity present within the Nestin/S100B spectrum and revealed a diversity of glioblastoma cell morphologies (Fig. 1d,e).

### Glioblastoma stem cell morphologies prognostic of poor outcome

At the apex of the neural stemness gradient, Nestin+ cells represent a treatment-resistant glioblastoma stem cell (GSC) subpopulation shown to sustain long-term tumour growth ^18,20,42–44^. Visual inspection of Nestin+ GSCs disclosed recurring cellular morphologies (“morphotypes”) distinguishable by the presence of tumour extensions as well as cell size and shape (Fig. 1d,f-h and Supplementary Fig. S2). Using deep learning on 51,028 manually curated single-cell image crops across all patient samples, we trained a convolutional neural network (CNN) to classify Nestin+ cells into four main *ex vivo* morphotypes (M1-M4; Fig. 1f,h and Extended Data Fig. 2a). Single-cell feature maps extracted from the CNN and nuclei segmentation revealed a continuum of M1-M3 morphotypes and a distinct cluster of small M4 cells (Fig. 1f,g and Extended Data Fig. 2b). M1 (polygonal with extensions; PE) and M2 (elongated with extensions; EE) GSC morphotypes had varying distributions of extensions per cell, largely overlapping with previously reported dimensions of tumour microtubes (TMs; Extended Data Fig. 2c) associated with glioma grade ^2^. M3 (round big; RB) and M4 (round small; RS) morphotypes without extensions were characterized by their roundness yet differed in cell size (Fig. 1g and Extended Data Fig. 2b). Gap-junction protein CX43 and Nestin expression were higher in M1-M3 than M4, while similar EGFR expression and only modest differences in cell viability were observed among the four morphotypes (Fig. 1g, Extended Data Fig. 2d-g). Across patients, GSC morphotype composition varied dramatically, with complex M1-M3 morphotypes ranging from 5% to 86% among Nestin+ GSCs (Fig. 1h).

To evaluate if this distinct contrast of patients with high or low *ex vivo* GSC morphotype complexity reflected *in situ* tumour organization, we imaged tumour regions in cohort-matched tissues from patients across the morphotype spectrum (M1-M3 high, ‘high complexity’, n=6; M1-M3 low, ‘low complexity’, n=4; Fig. 1h,i and Supplementary Fig.S3) by 100x confocal microscopy. Striking higher-level tumour organizational differences between the two patient groups were visually evident (Fig. 1i and Extended Data Fig. 2i), which coincided with significant stratification of multicellular immunohistochemistry (IHC) images by unsupervised deep learning (Fig. 1j). Subsequent single-cell image analysis of 12,799 Nestin+ cells and manual single-cell tracing demonstrated the presence of all four morphotypes *in situ* (Fig. 1k and Extended Data Fig. 2j). Comparison of quantitative morphological features across 4,000 Nestin+ cells (n=400 cells/patient) further confirmed significant differences between the two patient morphotype groups: Larger and more extended cell morphologies were more abundant in the high complexity patient group (Fig. 1k).

The morphological makeup of an individual patient’s Nestin+ GSCs was the strongest prognostic signal measured in our cohort. Higher baseline abundance of complex morphotypes (M1-M3% of Nestin+ cells) was associated with worse patient outcome (*P*=0.0047; n=17 patients with annotated PFS; Fig. 1l). The abundance of complex GSC morphotypes further correlated with Ki67 levels measured by pathology (Extended Data Fig. 2k). However, Ki67 levels alone did not stratify patient survival, nor did stratification based on IF marker-defined cell type composition (Extended Data Fig. 2l-n). While *MGMT* promoter methylation status, a prognostic factor associated with response to alkylating agents, also stratified patient survival in our cohort (*P*=0.038), complex GSC morphotype abundance was independent of *MGMT* status (Fisher’s test, *P*=0.19, Extended Data Fig. 2m). These results provide compelling evidence that complex M1-M3 GSC morphotypes and corresponding *in situ* tumour organization characterizes particularly aggressive disease with poor clinical outcome among glioblastomas.

### Therapeutically targeting neural tumour heterogeneity

The question arises whether it is possible to pharmacologically target the heterogeneous spectrum of glioblastoma cells in surgical patient samples, both in terms of neural stemness and morphological complexity. Image-based *ex vivo* drug screening (pharmacoscopy; PCY) enables measurements of drug-induced relative reduction of any marker- or morphology-defined cancer cell population. Positive PCY scores indicate a drug-induced reduction of cancer cells relative to non-malignant TME cells (Fig. 2a). In previous clinical trials, PCY identified effective therapies for aggressive haematological malignancies^33–35,45,46^. To evaluate the use of pharmacoscopy in glioblastoma, we measured *ex vivo* drug responses to first- and second-line glioblastoma chemotherapies (n=3 drugs) in patient samples from two independent patient cohorts: our main prospective cohort (n=27 patients) and a retrospective cohort (n=18 patients, Fig. 2b,c, Extended Data Fig. 3a-c, and Supplementary Table 1). Patient samples were either dissociated on the same day of surgery (prospective cohort), or dissociated after cell recovery from biobanking (retrospective cohort), and directly incubated with drugs for 48 hours. Limiting our analysis to newly diagnosed glioblastoma patients that received 1st-line Temozolomide (TMZ) treatment in the clinic, we found that *ex vivo* TMZ sensitivity of glioblastoma cells significantly stratified patient survival in both cohorts (Fig. 2b,c), recapitulating higher TMZ sensitivities in patients with tumours with *MGMT* promoter methylation (Extended Data Fig. 3c). Despite the limited success of targeted therapies for glioblastoma, we additionally tested an oncology drug library (ONCDs; n=65 drugs; Supplementary Table 3) across a subset of our prospective cohort (n=13 patients; Fig. 2d and Extended Data Fig. 3d-f). This also retrieved *ex vivo* drug sensitivities concordant with prior clinical trials in glioblastoma: for example, the RTK inhibitor Regorafenib was among the top ONCD hits and has shown preliminary signs of activity in a randomized phase II clinical trial for recurrent glioblastoma, while the mTORs inhibitors Temsirolimus and Everolimus showed no efficacy *ex vivo* and in clinical trials ^47–49^ (Extended Data Fig. 3d). When we explored associations between patient *ex vivo* ONCD responses and genetic alterations measured by targeted NGS, the strongest pharmacogenetic association was increased *ex vivo* sensitivity of patients with tumours carrying p53 mutations to the CDK4/6 inhibitor Abemaciclib (Extended Data Fig. 3e). Taken together, our evaluation of standard-of-care chemotherapies and oncology drugs by pharmacoscopy demonstrates the clinical concordance of image-based *ex vivo* drug profiling for glioblastoma.

**Fig. 2:**
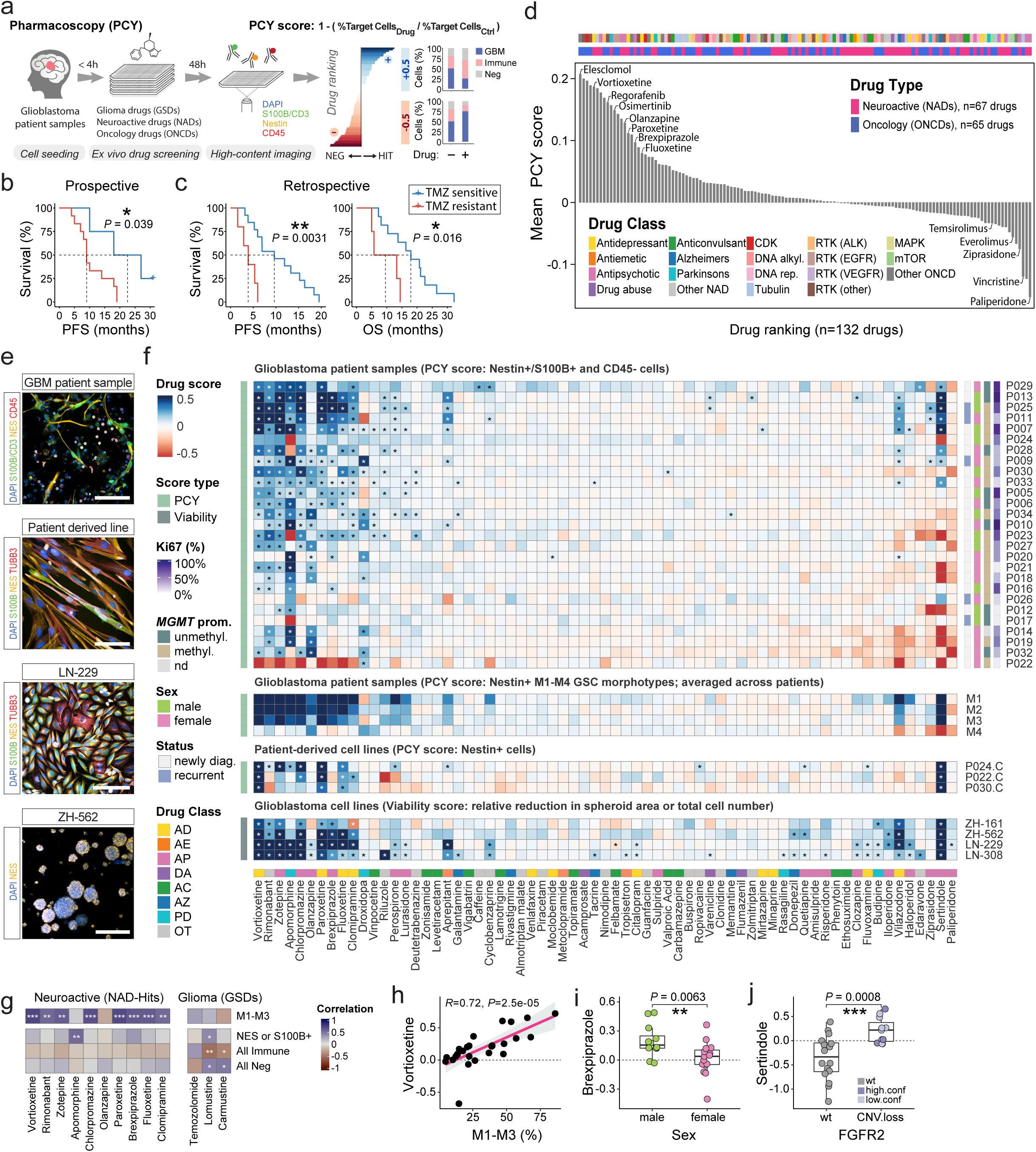
Image-based single-cell drug profiling across glioblastoma patient samples and model systems identifies repurposable neuroactive drugs. **a**, Workflow schematic showing image-based *ex vivo* drug screening (pharmacoscopy; PCY) of dissociated glioblastoma patient samples. The PCY score quantifies drug-induced “on-target” killing by measuring the change in fraction of a defined target population (e.g. Nestin+/S100B+ and CD45-glioblastoma cells) compared to the (-) vehicle control. Positive PCY scores (blue) indicate a drug-induced relative reduction of cancer cells compared to control, as illustrated in the stacked bar graphs on the right. **b-c,** Stratification of newly diagnosed glioblastoma patient survival based on *ex vivo* Temozolomide sensitivity (TMZ PCY score) of (Nestin+/S100B+ and CD45-) cells (blue, sensitive; red, resistant). Kaplan-Meier survival curves are compared using the log-rank (Mantel-Cox) test. **b**, Progression-free survival (PFS) of the prospective glioblastoma cohort (n=16 patients; *P*=0.039) stratified by 100µM TMZ PCY score. Tick mark indicates ongoing response. **c,** Progression-free survival (PFS; *P*=0.0031; left) and overall survival (OS; *P*=0.016; right) of the retrospective validation cohort (n=18 patients) stratified by mean TMZ PCY score. **d,** Drug ranking (n=132 drugs) by their mean (Nestin+/S100B+ and CD45-) PCY scores across glioblastoma patients (NADs, n=27 patients; ONCDs, n=12 patients). Drug annotations indicate drug type (NADs; n=67 drugs, ONCDs; n=65 drugs) and drug class. RTK, receptor tyrosine kinase; alkyl, alkylation; rep, replication. **e,** Representative immunofluorescence images of a glioblastoma patient sample (P040; scale bar, 100µm), a patient-derived cell line (P040.PDC; 100µm), an long-term glioblastoma cell line (LN-229; scale bar, 150µm), and a glioblastoma-initiating cell line (ZH-562, scale bar, 250µm). Cells are labeled with the nuclear stain DAPI (blue), astrocyte lineage marker S100B (green), and neural progenitor marker Nestin (yellow). Other markers are indicated in their respective colors. **f,** Drug response heatmaps of neuroactive drugs (NADs, n=67 drugs; columns) across glioblastoma patient samples (n=27 patients; rows), Nestin+ GSC M1-M4 morphotypes (n=4 classes; averaged response across n=27 patients), patient-derived lines (PDCs; n=3 lines, patient id followed by ‘.C’), and glioblastoma cell lines (n=4 lines). Drug score (heatmap color scale) indicates the PCY score for glioblastoma patient samples and patient-derived cell lines while for long-term glioblastoma cell lines the drug score is a viability score. Outliers beyond color scale limits set to minimum and maximum values. Clinical annotations per patient sample (rows) indicate the Ki67 labeling index, *MGMT* promoter methylation status (unmethyl, unmethylated; methyl, methylated; nd, not determined), sex, and newly diagnosed or recurrent tumour status. Annotation per drug indicates neuroactive drug class. Asterisks (*) denote FDR-adjusted *P* < 0.05 calculated by one-sided t-tests. **g,** Pearson correlations of marker and morphology-based sample composition at baseline (rows) with the (S100B+/Nestin+ and CD45-) PCY scores of top neuroactive drug hits (NAD-Hits) and glioma drugs (GSDs) across patients (n=27). **h,** Correlation of morphotype M1-M3 abundance of Nestin+ GSCs at baseline (x-axis) with Vortioxetine efficacy (y-axis; PCY score) across patients (n=27), as in **g**. Linear regression line with a 95% confidence interval. Pearson correlation coefficient with *P*-value annotated. **i,** Association of patient sex with response to Brexpiprazole (PCY score; *P* = 0.0063). **j,** Association of *FGFR2* copy number loss with response to Sertindole (PCY score; *P* = 0.0008). CNV, copy number variation; High.conf, high confidence; Low.conf, low confidence. **i,j,** *P*-values calculated from a two-sided Wilcoxon rank sum test. **g,i,j,** *P*-values: not significant (ns), P > 0.05, *P < 0.05, **P < 0.01, ***P < 0.001, ****P < 0.0001. Boxplots as in Fig. 1c.

The majority of oncological drugs, however, have limited access to the brain and are designed to target the transformed nature of cancer. Neuroactive drugs (NADs), in contrast, are developed to cross the blood-brain barrier and act upon the nervous system. To find repurposable neuroactive drugs that target the neural stemness and morphological features of glioblastoma, we tested a panel of NADs (n=67 drugs; Supplementary Table 3) across the prospective cohort (n=27 patients) by pharmacoscopy. The NAD library consisted of drugs approved for neurological diseases such as depression, schizophrenia, epilepsy, and Alzheimer’s disease. This identified 15 of 67 drugs (22%) with consistent anti-glioblastoma activity across patients (top NADs; PCY-hits; mean PCY score > 0.03; Fig. 2e,f and Extended Data Fig. 3e-g). Remarkably, top NADs effectively reduced fractions of aggressive M1-M3 GSC morphologies in patient samples *ex vivo*, reduced Nestin+ cells in patient-derived cultures (PDCs, n=3 lines), and reduced the viability of established glioblastoma cell lines also in the absence of the TME (n=4 lines, Fig. 2e,f). We additionally confirmed dose-response relationships in glioblastoma cell lines (n=9 drugs; Extended Data Fig. 4a,b and Supplementary Fig.S4) and robustness of the PCY score to potential technical factors (e.g. the presence of apoptotic cells after sample dissociation) in two glioblastoma patient samples (n=67 drugs; Extended Data Fig. 4c-e).

Among the NADs, the top mean ranking PCY-hit was Vortioxetine, a safe and novel class of antidepressant without known anti-glioblastoma activity (Fig. 2f). Strikingly, Vortioxetine and other top NADs were more potent in patient samples with higher baseline abundance of aggressive M1-M3 GSC morphologies, while standard-of-care chemotherapies did not show this association (Fig. 2g,h and Extended Data Fig. 4f). Other clinically attractive NADs included Paroxetine and Fluoxetine, both antidepressants of the selective serotonin reuptake inhibitor (SSRI) class, and Brexpiprazole, an atypical antipsychotic used for the treatment of schizophrenia. Brexpiprazole *ex vivo* response was related to biological sex, with increased drug sensitivities in male patient samples (Fig. 2i and Extended Data Fig. 3g). Sertindole response was highly variable among patient samples, despite its potency in the other evaluated glioblastoma models. This patient variability in *ex vivo* Sertindole response related to FGFR2 copy number loss, representing the most significant pharmacogenetic NAD association (Fig. 2j and Extended Data Fig. 3f). Not all top NADs were clinically attractive, considering the historically reported side-effects of the cannabinoid receptor blocker Rimonabant and the antipsychotic Zotepine, yet these could still provide mechanistic insights.

Thus, by comprehensively screening across heterogeneous patients and model systems, we identify a set of repurposable neuroactive drugs that effectively target the neural heterogeneity of glioblastoma cells. The consistency of the anti-glioblastoma efficacy of these neuroactive drugs across diverse model systems, even in the absence of the TME and a functioning *in vivo* nervous system, indicates that they target tumour-intrinsic vulnerabilities.

### Divergent functional dependencies on neuroactive drug targets

The multitude of neuroactive drugs with anti-glioblastoma activity was unexpected, prompting the question as to whether there could be shared underlying mechanisms. A previous screen of neurochemical compounds in patient-derived stem cell lines found various neurochemical classes represented among their hits^30^, and the antidepressant Fluoxetine has been reported to target glioblastoma metabolism ^32^. In our *ex vivo* patient drug screens, top NADs represented diverse drug classes without significant enrichment, indicating that canonical mode-of-action did not explain drug efficacy (Fig. 3a). Among our tested serotonin and dopamine pathway modulators, for example, only 4 out of 11 antidepressants (36%) and 6 out of 16 antipsychotics (38%) exhibited anti-glioblastoma activity in patient samples (Extended Data Fig. 4g). Such drug classifications, however, simplify the polypharmacological drug-target profiles of neuroactive drugs. The majority of NADs act on multiple primary target genes (PTGs). These include ion channels, GPCRs, and enzymes that modulate neurotransmission in the central nervous system, whose expression remains a largely unexplored dimension of glioblastoma heterogeneity. Dependency on neuroactive PTGs with high lineage specificity and consistent expression across patients could explain the activity of top NADs.

**Fig. 3:**
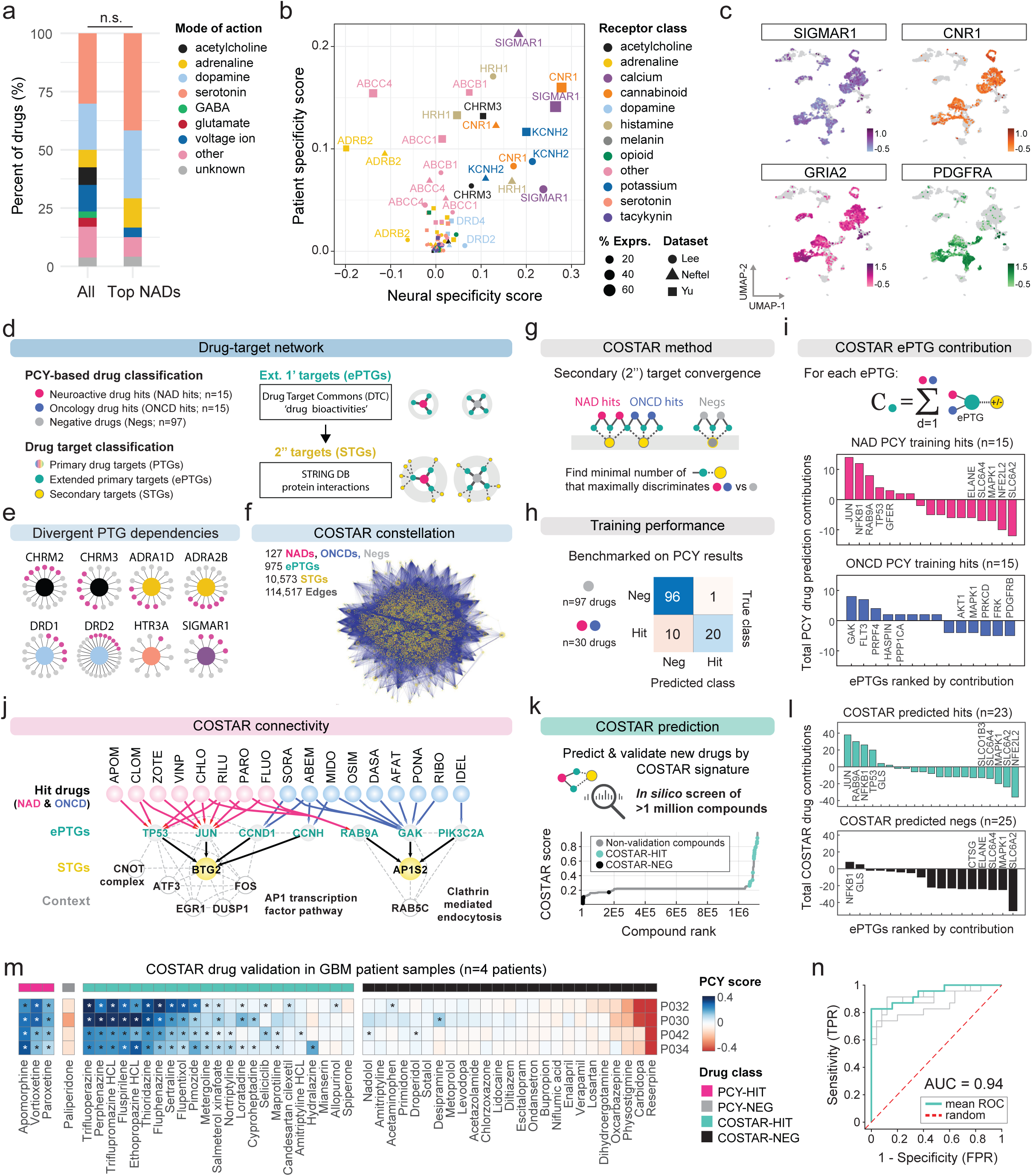
Neuroactive drugs with anti-glioblastoma efficacy converge upon an AP-1 and cell cycle connectivity signature through divergent primary targets. **a**, Drug mode-of-action for all neuroactive drugs (n=67 drugs; left) and top neuroactive drug hits (n=15 drugs with a mean patient PCY score > 0.03; right) represented as stacked bar plots. n.s., not significant by hypergeometric enrichment test. **b,** Primary target gene (PTG) expression of neuroactive drugs in 22 glioblastoma patient samples across three scRNA-Seq datasets (shape) plotted as the neural specificity score (x-axis) versus patient specificity score (y-axis) for each PTG (dot, gene; size, percent expression; color, receptor class). **c,** scRNA-Seq log10(expression) of selected neuroactive PTGs (*SIGMAR1*, *CNR1*, *GRIA2*) and oncogenic RTK (*PDGFRA*) visualized on the UMAP projection, as in Fig. 1b. **d,** Workflow for the collection of extended primary target genes (ePTGs) and associated secondary target genes (STGs) of drugs tested across patient samples by PCY to search for “convergence of secondary drug targets analyzed by regularized regression” (COSTAR). **e,** Example PTGs with genetic dependencies (core nodes; colored as in **b**; see also *Extended Data* Fig. 5c) linking to both PCY-HIT (pink) and PCY-NEG (grey) drugs. **f,** The full COSTAR network constellation of 127 PCY-tested drugs, 965 ePTGs, and 10573 STGs, connected by a total of 114517 edges **g,** COSTAR method: Logistic LASSO regression is performed on the COSTAR constellation to learn a linear model that discriminates PCY-HIT drugs (n=30, equally split across NADs and ONCDs) from PCY-NEG drugs (n=97) based on a small set of STGs. **h,** COSTAR training model performance as a confusion matrix, where the ‘true’ class denotes PCY-based experimental ground truth, and the ‘predicted’ class denotes the COSTAR-prediction. **i,** ePTGs (x-axis) ranked by their integrated contribution to predict a hit (+1) or non-hit (-1) (y-axis) in the COSTAR model, separated for PCY-hit NADs (top) and ONCD (bottom). **j,** COSTAR connectivity (solid lines) reveals convergence of NAD (pink) and ONCD (blue) hits to key ePTGs (grey) and STGs (yellow) included in the final model. See *Extended Data* Fig. 6b for the full model. Additional proteins (white nodes) with high confidence interactions to STGs (dashed lines) are shown. **i,** *In silico* COSTAR predictions based on drug-target connectivity across 1,120,823 compounds annotated in DTC. Compounds are ranked (x-axis) by their predicted PCY-hit probability (COSTAR score; y-axis). Predicted drug hits (COSTAR-HIT; mint green) and predicted non-hits (COSTAR-NEG; black) selected for experimental validation are indicated. **k,** As in **g**, but for COSTAR-HITs (top) and COSTAR-NEGs (bottom). **k,** Experimental validation by pharmacoscopy of COSTAR-HIT (n=23; mint green) and COSTAR-NEG (n=25; black) drugs (columns) across four glioblastoma patient samples (rows) including positive (PCY-HITs; pink; n=3) and negative (PCY-NEG; dark grey; n=1) control drugs. Heatmap color scale indicates the PCY score of (Nestin+/S100B+ and CD45-) cells. Outliers beyond color scale limits set to minimum and maximum values. Asterisks (*) denote FDR-adjusted P < 0.05. **l,** Receiver Operating Characteristic (ROC) curves (grey, n=4 patients; mint green, mean across patients; red dashed, random classifier) describing the COSTAR validation accuracy in glioblastoma patient samples of the COSTAR-predicted drugs (n=48 drugs; corresponding to Fig. 3m). AUC; area under the curve.

We therefore determined the expression of NAD PTGs by scRNA-Seq across the three independent datasets (Fig. 3b,c and Extended Data Fig. 5a,b) ^13,29^. Among PTGs with biochemical NAD-interactions reported in the Drug Targets Commons database (DTC; Fig. 3d)^50^, certain classes of ion channels and GPCRs were enriched in neural lineage cells (e.g. potassium channels, glutamate receptors, and cannabinoid receptors), while other classes showed broader expression patterns (e.g. calcium channels, adrenergic receptors; Extended Data Fig. 5a). To characterize PTG expression across cell types and patients, we defined neural- and patient-specificity scores (NS and PS; Fig. 3b, Extended Data Fig. 5b and Methods). For detected genes, a higher NS indicates relative enrichment in neural lineage cells (range -1 to 1) and a higher PS (range 0 to 1) indicates more patient-specific expression, while both scores will be close to zero for low-abundance genes (Extended Data Fig. 5b and Supplementary Table 4). Ion channels and receptors with high neural-specificity included the calcium signaling modulator SIGMAR1, glutamatergic AMPA receptor subunit GRIA2, and cannabinoid receptor CNR1 (Fig. 3c). Patient-specificity for neurological receptors SIGMAR1 and CNR1 were on average 1.7 to 3-fold lower than for oncogenic RTKs EGFR and PDGFRA, despite similar detection levels. Thus, we find abundant and consistent pan-patient expression of neuroactive drug targets on glioblastoma cells.

We next tested dependencies on these NAD PTGs by performing a reverse genetic screen in LN-229 glioblastoma cells (n=59 genes; Extended Data Fig. 5c,d and Supplementary Table 5), confirmed to have comparable PTG expression and NAD sensitivities to patient samples (Fig. 2f and Extended Data Fig. 5c). Knockdown of 9 PTGs significantly decreased cell viability (Extended Data Fig. 5c,d), of which lower expression levels of DRD1, DRD2, HTR3A, and TACR1 were also associated with better patient survival in The Cancer Genome Atlas (TCGA) glioblastoma cohort (4 out of 9; Extended Data Fig. 5e). However, these PTGs showing genetic dependencies were predominantly targeted by NADs that showed no anti-glioblastoma activity by PCY. For example, only 5 of the 16 NADs interacting with DRD1 were PCY-hits, and only 1 out of 11 NADs interacting with HTR3A was a PCY-hit (Fig. 3e). Therefore, while presenting possible neural vulnerabilities, these genetic PTG dependencies did not explain the anti-glioblastoma activity of our top neuroactive drug hits.

### Anti-glioblastoma activity explained by drug-target convergence

Despite their chemical and primary target diversity, our top NADs may still converge upon common downstream signaling pathways. To test this, we developed an interpretable machine learning approach that searches for “convergence of secondary drug targets analyzed by regularized regression” (COSTAR). COSTAR is designed to identify the minimal drug-target connectivity signature predictive of efficacy.

We expanded the drug-target search space to include PTGs with any bioactivity annotated by DTC, termed extended PTGs (ePTGs). Secondary target genes (STGs) downstream of ePTGs were subsequently mapped by high-confidence protein-protein interactions annotated in the STRING database (Fig. 3d). This resulted in a drug-target connectivity map, or “COSTAR constellation”, of all DTC-annotated drugs in our NAD and ONCD libraries (n=127 of 132 tested drugs) with 975 extended primary targets, 10,573 secondary targets, and 114,517 network edges (Fig. 3f). Using logistic LASSO regression, we trained a multi-linear model that identifies the minimal set of STGs that maximally discriminates PCY-hit drugs (n=30; top-15 drugs from both NADs and ONCDs) from PCY-negative drugs (n=97; all other tested drugs) in a cross-validation setting (Fig. 3g,h Extended Data Fig. 6a, and Methods). Thereby, COSTAR converged upon the minimal connectivity signature that was predictive of anti-glioblastoma drug efficacy (Fig. 3g and Extended Data Fig. 6a,b). Encouragingly, COSTAR identified a signature that classified the 127 drugs in our training data with 92.1% accuracy, correctly predicting 20/30 PCY-hits and 96/97 negative drugs (Fig. 3h).

The COSTAR connectivity signature linked PCY-hit NADs to the secondary target BTG2, predominantly through JUN and TP53 ePTGs (Fig. 3i,j and Extended Data Fig. 6b). BTG2 and TP53 are both tumour suppressors that control cell cycle and differentiation, while JUN is a member of the AP-1 transcription factor (TF) family that, in a neural context, regulates gene expression and apoptosis in response to stimuli such as neural activity or insult ^36,38^. Conversely, the majority of PCY-hit ONCDs were connected to the secondary target AP1S2, a protein involved in clathrin coat assembly, through the cyclin G-associated kinase GAK (Fig. 3i,j and Extended Data Fig. 6b). A subset of PCY-hit ONCDs were also linked to BTG2 through cyclins CCND1 and CCNH, while a subset of PCY-hit NADs were linked to AP1S2 through RAB9A, a member of the RAS oncogene family (Fig. 3j). Taken together, this reveals pathway convergence on AP-1 transcription factors and cell cycle regulation as a unique signature predictive of anti-glioblastoma activity of neuroactive drugs.

COSTAR can compute the hit probability (COSTAR score) of any annotated compound, by matching its drug target profile to the learned connectivity signature. To evaluate the predictive power of the COSTAR signature and find additional neuroactive drug candidates with anti-glioblastoma activity, we performed a large-scale *in silico* drug screen of 1,120,823 DTC-annotated compounds, and experimentally validated 48 previously untested drugs among the top (COSTAR-hits, n=23) and bottom (COSTAR-negs, n=25) scoring compounds (Fig. 3k-n). All COSTAR-hits were linked to the secondary target BTG2 primarily through JUN (Fig. 3l), while none of the COSTAR-negs had annotated connections to BTG2 (Extended Data Fig. 6c). We experimentally tested all 48 drugs across four GBM patient samples *ex vivo* (P030, P032, P034, P042), and observed excellent agreement between COSTAR predictions and *ex vivo* results (mean AUC=0.94, Fig. 3m,n). The COSTAR-hits again represented diverse drug classes, including the antipsychotic Trifluoperazine, antiparkinsonian Ethopropazine, antidepressant Sertraline, and bronchodilator Salmeterol (Fig. 3m). These results substantiate AP-1 transcription factor and cell cycle signaling pathway convergence as an actionable signature of neuroactive drugs with *ex vivo* anti-glioblastoma activity.

### From neural activity-dependent signaling to tumour suppression

The convergent COSTAR signature suggested a common gene regulatory network (GRN) underlying the anti-glioblastoma activity of top NADs. We determined the transcriptional response of LN-229 cells at 6 and 22 hours to PCY-hit NADs (n=11) spanning diverse drug classes, PCY-hit ONCDs (n=7), and negative controls (NEG; n=2 PCY-neg NADs and DMSO; Fig. 4a-d, Extended Data Fig. 7a,b, and Supplementary Table 3). In remarkable alignment with COSTAR, differential gene expression analysis upon PCY-hit NAD treatment (PCY-hit NADs vs NEGs) revealed a common transcriptional response involving AP-1 and BTG family members (Fig. 4b,d and Extended Data Fig. 7e). This AP-1/BTG upregulation was observed even for Vortioxetine and Brexpiprazole, both lacking DTC-annotations at the time of analysis and thus not contributing to the COSTAR training (Fig. 4d).

**Fig. 4:**
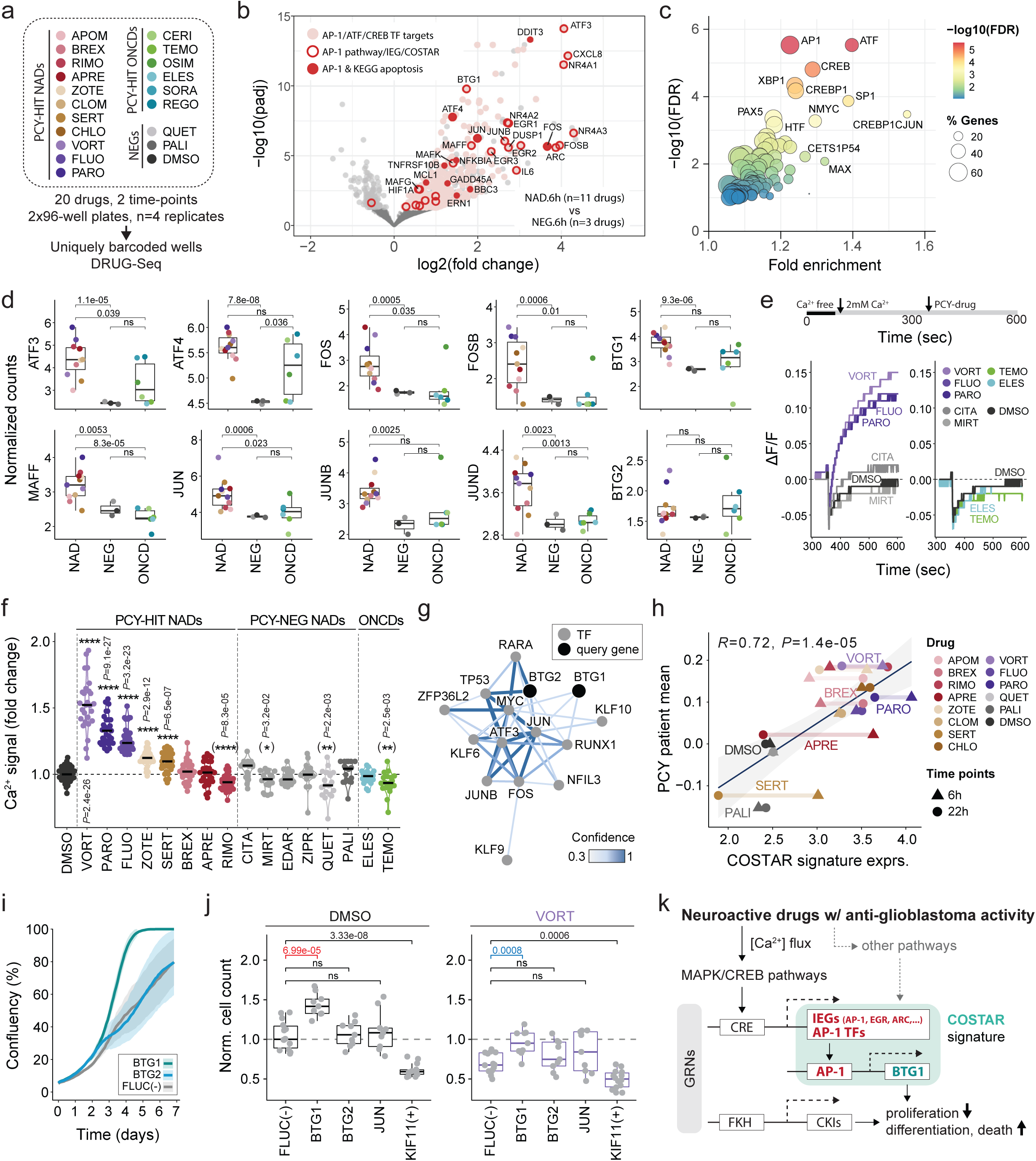
Glioblastoma suppression is driven by a tumour-intrinsic AP-1 gene regulatory network. **a**, Multiplexed RNA-Seq (DRUG-Seq^79,80^) of LN-229 cells after pharmacoscopy-hit neuroactive drug (PCY-hit NADs, n=11 drugs), pharmacoscopy-hit oncology drug (PCY-hit ONCDs, n=6 drugs), and negative control drug (NEGs, n=2 PCY-neg NADs and DMSO vehicle control) treatment (n=4 replicate wells per drug/time-point; n=2 time-points, 6 and 22 hours). **b**, Transcriptional response of PCY-hit NAD-treated cells compared to NEG-treated cells (6 hours). X-axis: log2(fold change), positive value indicating PCY-hit NAD upregulated genes; y-axis: –log10(adjusted *P*-value). Genes above a –log10(0.05 adjusted P-value) threshold (light grey or colored), and non-significant genes (dark grey). Target genes of AP-1, ATF, or CREB TFs (pink), as well as AP-1 pathway, IEG, and COSTAR model member genes (red outlines), and genes present in both the AP-1 & KEGG apoptosis pathway(solid red) are indicated. Gene names of indicated genes with an adjusted *P*-value < 0.01 are shown. **c**, Transcription factor binding site enrichment analysis of significantly upregulated genes upon PCY-hit NAD treatment in Fig. 4b. Circles correspond to TF annotations, sizes scale with the percent of genes present in the annotation, and colors indicate –log10(false discovery rate). **d**, Expression of AP-1 TF and BTG family genes (y-axis, normalized RNA-Seq counts) that are significantly upregulated upon PCY-hit NAD treatment (6 hours). Box plot groups (x-axis) correspond to drug categories and dots represent the average expression per drug. ‘PCY-hit NAD’ and ‘PCY-hit ONCD’ abbreviated to NAD and ONCD, respectively. **e**, Calcium response (ΔF/F; y-axis) over time (x-axis) of LN-229 cells upon drug treatment measured by high-throughput FLIPR assay. Timeline depicts assay setup (*Methods*). Representative traces from 8 drug conditions (out of 17 tested) including 5 NADs (left), and 2 ONCDs (right). DMSO vehicle control traces shown in both. ΔF/F, change in fluorescence intensity relative to the baseline. **f**, Fold-change in extracellular calcium influx upon drug treatment relative to DMSO vehicle control measured by FLIPR assays in LN-229 cells (n=8 assay plates; n=17 conditions; n=18-30 wells/drug; DMSO, n=47 wells). Drug categories including PCY-hit NADs, n=8 drugs; PCY-neg NADs, n=6 drugs; PCY-hit ONCDs, n=2 drugs were compared. Asterisks (*) in parentheses denote conditions where the median [Ca2+ fold change] < 0. Black line indicates the median value. **g**, Transcriptional regulation of BTG1/2 based on PathwayNet ^60^. Query genes (*BTG1/2*, black nodes) and the top-13 inferred transcription factor interactions (grey nodes) are shown. Edge colors indicate relationship confidence. **h**, Correlation of average COSTAR signature expression (x-axis) with *ex vivo* patient neuroactive drug response (y-axis) plotted per drug (color) and time-point (shape). Mean glioblastoma PCY score across patients (n=27 patients) of neuroactive drugs (n=11 PCY-H NADs, n=3 NEGs) plotted against their corresponding geometric mean expression of AP-1 TFs and *BTG1/2* genes as shown in Fig. 4d. Linear regression line (black) with a 95% confidence interval (light grey). Pearson correlation coefficient R=0.72, *P*-value 1.4e-05. **i,** Confluency of LN-229 cells measured by IncuCyte live-cell imaging (y-axis) across 7 days (x-axis) in two siRNA knockdown conditions (*BTG1*, *BTG2*) and a negative firefly luciferase control (*FLUC*). Mean of n=4 replicate wells shown +/-one standard deviation. **j**, Effect of target gene siRNA knockdown (columns) on normalized LN-229 cell counts (y-axis) at baseline (DMSO; left panel) and upon Vortioxetine treatment (VORT; 10µM; right panel). *KIF11*; positive (+) control. Cell counts are normalized to the *FLUC* negative (-) control siRNA within the DMSO condition per experiment (n=9-14 replicate wells/condition, n=2 experiments). **k**, Diagram summarizing mechanistic pathways by which neuroactive drugs target glioblastoma. GRN; gene regulatory network. IEG; immediate early gene. CKI: cyclin-dependent kinase inhibitor. CRE; cAMP response element. FKH; forkhead binding motif. **a,d-f,h,** Colors correspond to drugs and drug name abbreviations annotated in *Supplementary Table 3*. **d,f,j** Two-sided t-test. **f,j** *P*-values adjusted for multiple comparisons by Holm correction. *P*-values: not significant (ns), P > 0.05, *P < 0.05, **P < 0.01, ***P < 0.001, ****P < 0.0001. Boxplots as in Fig. 1c.

In response to PCY-hit NAD treatment, we observed rapid and sustained upregulation of several AP-1 TFs, such as the canonical immediate-early genes (IEGs) JUN and FOS, with well-known roles in mediating neural activity and apoptosis, and stress-induced AP-1 family members ATF3 and ATF4, where ATF3 represented the most significantly upregulated gene across both time-points. The presence of other upregulated IEGs including NR4A1, EGR1, and ARC, and pathway enrichment in MAPK signaling, strengthened this surprising involvement of neural-activity dependent signaling in glioblastoma (Fig. 4b and Extended Data Fig. 7d). BTG1, a close homologue of BTG2 identified by COSTAR, was also among the top 20 most significant genes (Fig. 4b,d and Extended Data Fig. 7c) while BTG2 was strongly induced in response to select drugs, including Vortioxetine and Paroxetine (Fig. 4d). Induction of AP-1 factors was primarily NAD-specific, where ONCD treatment did not elicit a similar global transcriptional response (Fig. 4d and Extended Data Fig. 7c). Additionally, over half of AP-1 factors showed no transcriptional upregulation (Extended Data Fig. 7e). For example, ATF2 expression remained unchanged, despite it being one of the key dimerization partners of JUN ^51^, as did FOSL1, previously implicated in response to irradiation in glioblastoma ^52^.

To find the transcriptional regulators mediating the response to PCY-hit NADs, we performed transcription factor binding-site (TFBS) enrichment analysis of the upregulated genes (Fig. 4c and Extended Data Fig. 7f). The most significantly enriched TF motifs at 6 hours were AP-1, ATF, and CREB a calcium-activated regulator of AP-1 transcription ^36,53,54^. Over 60% of upregulated genes were annotated targets of AP-1/ATF/CREB TFs (n=434 out of 719 genes; Fig. 4b,c). Though NAD-induced AP-1 expression was sustained across both time-points, TFBS enrichment analysis of upregulated genes at the later 22 hour time-point identified forkhead TF family members (e.g. FOXO1, FOXO3, FOXD3, HFH1) known to regulate long-term cell differentiation as a gene regulatory module succeeding AP-1^55–57^ (Extended Data Fig. 7f).

A rapid Ca^2+^ influx and calcium-dependent signaling typically precede IEG expression and AP-1 activation in neural lineage cells ^36,54,58,59^. We therefore measured both NAD-mediated extracellular calcium influx as well as endoplasmic reticulum (ER) calcium store release in LN-229 cells by high-throughput FLIPR calcium assays (n=17-18 drugs; Supplementary Table 3). We observed an immediate and strong extracellular Ca^2+^ influx in response to 5 out of 8 of our PCY-hit NADs, while none led to ER Ca^2+^ store release (Fig. 4e,f and Extended Data Fig. 8a). The strongest Ca^2+^ influxes were triggered by antidepressants Vortioxetine, Paroxetine and Fluoxetine (Fig. 4e,f). In contrast, the PCY-neg NADs (n=6) including antidepressants Citalopram and Mirtazapine, and PCY-hit ONCDs Elesclomol and TMZ did not trigger calcium influxes (Fig. 4e,f). These results demonstrate that for the majority of our top NADs a rapid drug-induced Ca^2+^ influx precedes IEG upregulation and subsequent anti-glioblastoma activity.

Downstream of AP-1, we evaluated whether BTG tumour suppressors could be direct effectors of the AP-1 gene regulatory network. To delineate regulators of BTG family genes, we leveraged genome-wide mapping of transcriptional regulatory networks by PathwayNet, a tissue-aware data integration approach that utilizes 690 ChIP-Seq datasets from the ENCODE project ^60^. The most enriched transcriptional regulators of BTG1/2 were members of the AP-1 TF network (e.g. JUN, ATF3, FOS), implying BTG tumour suppressor gene expression is directly mediated by AP-1 factors (Fig. 4g). Congruence between NAD-induced AP-1/BTG activation and its anti-glioblastoma activity would strengthen a causal role for this gene regulatory network. Indeed, drug-induced expression of this COSTAR signature was strongly correlated with a drug’s *ex vivo* anti-glioblastoma efficacy in patient samples (R=0.72, *P*=1.4e-05; Fig. 4h). We additionally performed BTG1/2 and JUN loss-of-function experiments by siRNA-mediated knockdown in LN-229 cells. Quantitative RT-PCR after 72 hours of gene silencing confirmed reduced expression of BTG1/2 and JUN and revealed interdependent regulatory interactions governing their expression (Extended Data Fig. 8b). Particularly BTG1 inhibition accelerated cell growth measured by live-cell imaging across 7 days (Fig. 4i, Supplementary Video 1), and increased the total number of cells measured by IF after 3 days (Fig. 4j). Furthermore, after two days of siRNA-mediated gene silencing and one subsequent day of top-NAD Vortioxetine treatment, BTG1 inhibition attenuated Vortioxetine’s anti-glioblastoma efficacy (Fig. 4j).

Together, these results propose a model in which neuroactive drugs that mediate anti-glioblastoma activity trigger a rapid calcium influx, IEG and AP-1 transcription factor activation, and engagement of an antiproliferative program that includes BTG-driven tumour suppression (Fig. 4k).

### AP-1 orchestrated anti-glioblastoma activity of neuro-active drugs

To further delineate the molecular dynamics of this discovered anti-glioblastoma program, we performed deep transcriptomic, proteomic, and phosphoproteomic profiling at 3-6 time-points in Vortioxetine treated LN-229 cells (Fig. 5a and Extended Data Fig. 8c-h). NH-2 terminal JUN phosphorylation occurring within 30 minutes to 3 hours after Vortioxetine treatment was central to several differentially phosphorylated pathways, including the stress response pathway (e.g. HSPB1, HSP90B1, RIPK2), mRNA processing (HRNPA2B1, NONO), and clathrin mediated endocytosis (DNM2, M6PR) (Extended Data Fig. 8h). Consistent with this observation, a number of AP-1 TFs and associated pathway annotations were upregulated at both the RNA and protein level across all timepoints. This included induction of FOS, JUNB, ATF4 already at 3 hours, as well as the ER stress response, DNA damage, and MAPK signaling pathways (Fig. 5a and Extended Data Fig. 8e). We also observed upregulation of BTG1/2 and negative cell cycle regulators CDKN1B and PPM1B (Fig. 5a and Extended Data Fig. 8g). Conversely, cytoskeletal components and oncogenic RTKs associated with the malignant phenotype of glioblastoma, including EGFR, NTRK2, and PDGFRA, were downregulated upon Vortioxetine treatment (Fig. 5a).

**Fig. 5:**
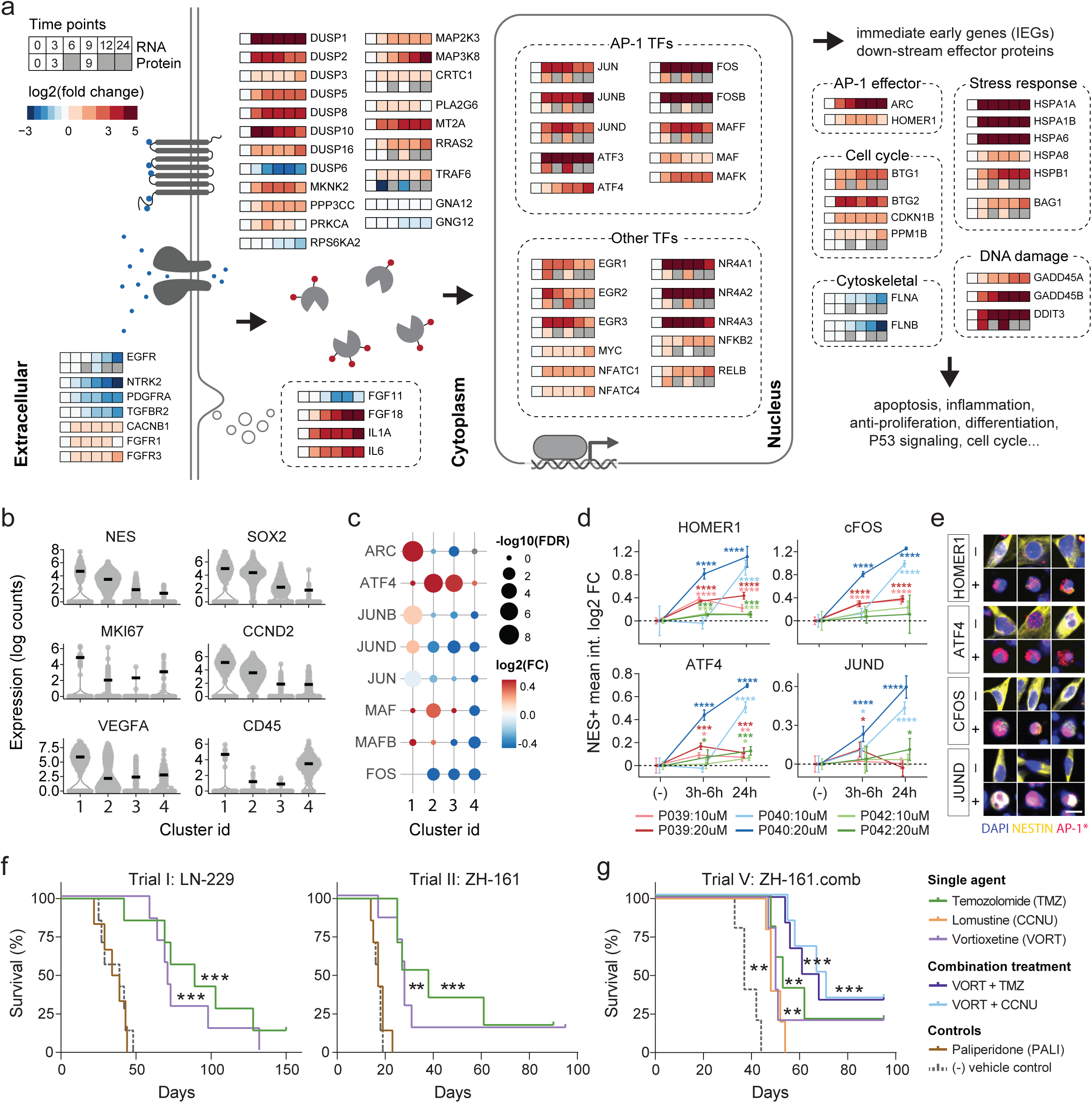
The antidepressant Vortioxetine induces a potent AP-1 response that synergizes *in vivo* with current standard of care drugs for glioblastoma. **a**, Time-course visualization of AP-1 (PID) and MAPK (KEGG) pathway induction following Vortioxetine treatment (20µM) in LN-229 cells measured by RNA-Seq (n=6 time-points) and by proteomics (n=3 time-points). n=3 replicates/time-point. Genes selected for visualization are significantly differentially expressed by RNA-Seq at all time-points compared to the first time-point (0h). Heatmap color scale represents log2(fold change) compared to the 0h time-point. **b,** scRNA-Seq expression log2(UMI) of selected glioblastoma and top cluster marker genes from glioblastoma patient sample P024. Cluster ids are based upon UMAP clustering of both DMSO and Vortioxetine (VORT, 20µM) treated cells (3h) shown in *Extended Data* Fig. 10a. Black lines indicate the median value. **c,** Differentially expressed AP-1 transcription factors and downstream effector gene ARC per scRNA-Seq cluster upon Vortioxetine treatment compared to DMSO in P024. Circle sizes scale with the –log10 (adjusted *P*-value) and heatmap color scale represents VORT-induced log2(fold change) compared to DMSO treated cells per cluster. **d,** Induction of AP-1 transcription factors and downstream effector *HOMER1* in glioblastoma patient samples (n=3 additional patients; P039, P040, P042) upon *ex vivo* Vortioxetine treatment (time-points: 0, 3-6, and 24 hours; concentrations: 10 and 20µM) in Nestin+ cells measured by immunofluorescence. Two-sided t-test compared to negative control. **e,** Representative pseudo-colored single-cell image crops from glioblastoma patient P040 of Nestin+ (yellow) cells after Vortioxetine treatment (+; 20µM) and DMSO vehicle control (-) at 24 hours stained with different AP-1 transcription factors/ HOMER1 (red) and DAPI (blue). Scale bar, 15µm. **f,** Survival analysis of Trial I: LN-229 (left) or Trial II: ZH-161 (right) tumour-bearing mice (n=6-7 mice/group). Mice were treated intraperitoneally (*i.p.*) between days 5-21 after tumour implantation with a PCY-HIT NAD, Vortioxetine (VORT; 10mg/kg; Trial I, *P*=0.0001; Trial II, *P*=0.0016), a positive control, Temozolomide (TMZ; 50mg/kg; Trial I, *P*=0.0009; Trial II, *P*=0.0002), a PCY-NEG NAD, Paliperidone (PALI; 5mg/kg), and a negative vehicle control. *See also Extended Data* Fig. 9 *for Trial III-IV and full results of Trials I and II including other PCY-hit NADs.* **g,** Trial V: *in vivo* treatment of Vortioxetine (VORT; 10mg/kg) in combination with 1st- and 2nd-line glioblastoma chemotherapies; Temozolomide (TMZ; 50mg/kg) and Lomustine (CCNU; 20mg/kg) compared to single-agents and vehicle control in ZH-161 tumour-bearing mice (n=5-6 mice/group). Combination treatments, TMZ+VORT/CCNU+VORT, both *P*=0.0007; Single-agents, TMZ/CCNU/VORT, all *P*=0.0018. **f-g,** Survival plotted as Kaplan–Meier curves and P values calculated using log-rank (Mantel-Cox) test. Censored mice denoted as tick marks. *P*-values: not significant (ns) P > 0.05, *P < 0.05, **P < 0.01, ***P < 0.001, ****P < 0.0001.

Next, we determined the cell type specificity of AP-1 induction at the single-cell gene and protein level in compositionally heterogeneous glioblastoma patient samples. We performed scRNA-Seq on dissociated cells from patient P024 following 3 hours of *ex vivo* Vortioxetine treatment (Fig. 5b,c and Supplementary Fig. S5). Analysis of 1736 single-cell transcriptomes revealed 4 main clusters intermixed with Vortioxetine-treated and DMSO-control cells (Supplementary Fig. S5a). Clusters 1 to 3 represented glioblastoma cells expressing Nestin, Ki67, CCND2, and VEGFA, with Cluster 1 showing the most aggressive signature, while Cluster 4 represented immune cells (Fig. 5b and Supplementary Fig. S5b). Analyzing the transcriptional response to Vortioxetine treatment confirmed glioblastoma-specific induction of AP-1 factors (Fig. 5c). For example, JUNB, JUND, and AP-1 effector gene ARC were upregulated in Cluster 1, while ATF4 and MAF were induced in all three glioblastoma clusters, with a more pronounced induction of ATF4 in Clusters 2 and 3 (Fig. 5c). Immunofluorescence against AP-1 pathway members in three additional glioblastoma patient samples (P039, P040, P042) following Vortioxetine treatment demonstrated the patient-, time-point, and concentration-dependent AP-1 induction in Nestin+ glioblastoma cells (Fig. 5d,e). The strongest induction was seen in patient sample P040 that had high abundance of complex GSC morphotypes (M1-M3), which were reduced upon Vortioxetine treatment (Fig. 5d,e). HOMER1 and ATF4 were induced in all three patient samples, while FOS and JUND exhibited more patient variability (Fig. 5d). Together, this single-cell analysis highlights the added dimension of cellular and patient complexity, yet supports AP-1 induction to be a key neural vulnerability targeted by PCY-hit NADs.

### Preclinical translation of neuroactive drugs

To evaluate the *in vivo* anti-glioblastoma efficacy of our top neuroactive drugs, we tested PCY-hit NADs spanning different drug classes in orthotopic human-xenograft glioblastoma mouse models (n=4 or 5 drugs; Fig. 5f and Extended Data Fig. 9a-c). We accounted for the variability observed in different orthotopic models by evaluating two different models (LN-229, ZH-161) across three independent trials (Trials I-III) of *in vivo* drug-testing (Fig. 5f and Extended Data Fig. 9a). We included Temozolomide (TMZ) as a positive control, and as negative controls we tested PCY-neg NAD Paliperidone and a vehicle control. Since all tested NADs have confirmed BBB-penetrance and are approved for other neurological disorders, doses were determined *a priori* based on literature and clinical evidence ^61–66^. Vortioxetine was consistently the most effective drug *in vivo* (in 3/3 trials), showing significant survival benefit comparable to the chemotherapeutic TMZ despite being tested at considerably lower dosage (Fig. 5f). Brexpiprazole was the 2nd-best PCY-hit NAD (2/3 trials), while other NADs showed a significant survival benefit in 1 out of 3 trials (Extended Data Fig. 9a). Consistent with PCY, the negative control Paliperidone did not show a significant survival benefit (2/2 trials) (Fig. 5f). In the most aggressive orthotopic model with the shortest median survival of the vehicle control, Vortioxetine and TMZ were the only effective drugs (Trial II: ZH-161; Fig. 5f, right), whereas for the least aggressive model, all 5 tested PCY-hit NADs significantly prolonged survival (Trial III: ZH-161; Extended Data Fig. 9a). MRI images of ZH-161 transplanted mice (Trial II) after 15 days of Vortioxetine, Apomorphine, and Temozolomide treatment showed marked reduction of tumour size (Extended Data Fig. 9b,c). Finally, we confirmed that the potent efficacy of Vortioxetine is not common to serotonin modulating drugs by directly comparing Vortioxetine to the PCY-neg antidepressant Citalopram at the same dose (10mg/kg) in an additional *in vivo* trial (Trial IV; Extended Data Fig. 9d-h). Unlike Citalopram, Vortioxetine again provided a robust survival benefit (Extended Data Fig. 9d), and reduced tumour burden (Extended Data Fig. 9e,f) and Ki67 levels (Extended Data Fig. 9g,h).

The striking consistency of our patient *ex vivo* and mouse *in vivo* results demonstrates strong translatability of PCY-based NAD discovery and confirms Vortioxetine as the most promising clinical candidate. Vortioxetine furthermore displayed multifaceted anti-tumour effects *in vitro*, reducing glioblastoma invasiveness (Extended Data Fig. 10a,b), long-term survival (Extended Data Fig. 10c), and growth (Extended Data Fig. 10d) across 2D and 3D glioblastoma cell lines (2D cultures: LN-229 and, LN-308; 3D spheroids: ZH-161 and ZH-562). Lastly, we tested the combination of Vortioxetine with either first- or second-line standard of care drugs for glioblastoma, TMZ and Lomustine (CCNU) *in vivo* (Trial V: ZH-161; Fig. 5g). All three single agents significantly prolonged survival, with Vortioxetine results now confirmed in 5 out of 5 *in vivo* trials (Fig. 5f,g and Extended Data Fig. 9a,d). Remarkably, compared to TMZ or CCNU single agents, the combination of Vortioxetine with either drug provided a further median survival increase of 20-30% compared to the single agents (Fig. 5g).

This strong preclinical evidence of the anti-glioblastoma efficacy of the safe antidepressant Vortioxetine urges for clinical investigation of Vortioxetine in patients. Given the complementary mechanisms of neuroactive drugs and approved chemotherapies, their successful combination could facilitate the rapid adoption of NADs into clinical routine for this dire disease.

## Discussion

Here we present the first therapeutic single-cell map across glioblastoma patient samples that reveals the morphological and neural molecular heterogeneity of glioblastoma. Glioblastoma stem cells adopt distinct cell morphological states that reflect *in situ* tumor organization and encodes clinical prognostic value. While the presence of tumour microtubes has been associated with tumour grade ^2^, we now show that, even within glioblastoma, complex GSC morphologies are prognostic of shorter progression-free survival. Critical to this discovery is the image-based evaluation of minimally-cultured surgical patient samples, which empowers scalable drug screening (pharmacoscopy; PCY) across a genetically and clinically heterogeneous patient cohort.

PCY-based *ex vivo* drug sensitivities predicted clinical response to chemotherapy and enabled the discovery of repurposable neuroactive drugs that target the spectrum of glioblastoma cells across 27 patients and various model systems, greatly expanding upon prior literature ^67–69^. Response to the antidepressant Vortioxetine, the most promising preclinical neuroactive candidate, was particularly aligned with aggressive GSC morphotypes. These efforts expand the nascent community of glioblastoma research focusing on the investigation of patient-derived tumour explants that facilitate translational investigation of complex tumour behavior, including the development of genetically characterized patient cultures, organoid biobanks, and regionally annotated samples ^30,41,70–74^. As pharmacoscopy allows patient-tailored evaluation of tumour-extrinsic responses to immuno- and cell-based therapies ^46,75–77^, further development may enable investigation of additional complex tumour physiology, including the neuron-glioma interface.

Our systematic analysis of the neuroactive drug mechanisms, drug target expression, and functional genetic dependencies indicated a diverse set of possible neural vulnerabilities of glioblastoma. Despite this diversity, our interpretable machine learning approach COSTAR identified a simple underlying drug-target connectivity signature predictive of anti-glioblastoma efficacy. COSTAR effectively applies Occam’s razor to the collective biochemical drug-protein-protein interaction network, offering a novel conceptual framework applicable to all fields of drug discovery. Through COSTAR, we uncovered a convergence of AP-1 transcription factor activity and cell cycle regulation on BTG-mediated tumour suppression. AP-1 and BTG upregulation was a defining feature of the response to neuroactive drugs with anti-glioblastoma activity, where a growth-suppressing role for BTG1 was confirmed by functional genetics. While the key pharmacological properties leading to AP-1 upregulation remain to be identified, and additional mechanisms may still contribute to the integrated effect of each individual drug, our results reveal diverse neuro-active drugs converging on this novel and potent glioblastoma-suppressing pathway.

Previous studies have demonstrated the role of neuronal input in regulating glioblastoma growth at the brain-tumour interface, highlighting the influence of the tumour microenvironment in modulating the neural behavior of the tumour ^3,5,6,9^. Here, we uncover a *cell*-*intrinsic* AP-1 mediated neural vulnerability in glioblastoma, offering a therapeutic window that enables direct targeting of the tumour. In cancers, AP-1 factors were originally discovered as oncogenes, though an increasing number of studies report context-dependent anti-oncogenic functions of AP-1 factors ^78^. In contrast, for neural lineage cells such as neurons, immediate early gene expression of AP-1 factors is typically a hallmark of neural activity or insult ^36,38^. We now find that neuroactive drugs can target this activity-dependent neural signaling, triggering a strong transcriptional response that, in the context of glioblastoma cells, leads to rapid cell death. Treating glioblastoma tailored to the cellular history and lineage of the cancer rather than its unstably transformed state may represent new hope for this devastating disease.

## Data Availability

All transcriptomics data generated in this study including single-cell RNA-Seq, bulk RNA-Seq, and DRUG-Seq datasets have been deposited in the public repository NCBI Gene Expression Omnibus (GEO; https://www.ncbi.nlm.nih.gov/geo/) under the following accession numbers: <u>GSE214965</u> (DRUG-Seq; multiplexed RNA-Seq of 20 drugs, 2 time points; Reviewer token: **uxglimouvdszzwh**), <u>GSE214966</u> (scRNA-Seq; 4 glioblastoma patients at baseline; Reviewer token: **szezsuewrhcrpcl**), <u>GSE214967</u> (scRNA-Seq; glioblastoma patient sample after Vortioxetine vs DMSO treatment; Reviewer token: **kdaligmwptuvlin**), and <u>GSE214968</u> (RNA-Seq; Vortioxetine time course; Reviewer token: **yhadscswlxwnzkv**). Previously published single-cell RNA-Seq datasets analyzed in this study are publicly available at GEO under accession numbers <u>GSE117891</u> and <u>GSE131928</u>. Proteomics and phosphoproteomics data can be accessed via Panorama (https://panoramaweb.org/GlioB.url; Username, Password: **TUqPvoSy**). DIA and phosphopeptide enrichment datasets are available from MASSIVE (ftp://massive.ucsd.edu/MSV000090357/; Username: **MSV000090357**, Password: **wlab@2022**). Drug-target annotations and protein-protein interaction data were retrieved from the following publicly available databases: Drug Target Commons(DTC; https://drugtargetcommons.fimm.fi/) and STRING (https://string-db.org/). Other publicly available databases used in this study include DAVID (https://david.ncifcrf.gov/), KEGG (https://www.genome.jp/kegg/), Gene Ontology (http://geneontology.org/), and PathwayNet (http://pathwaynet.princeton.edu/). Data provided via Supplementary Tables include *ex vivo* drug response of glioblastoma cells (pharmacoscopy scores; Supplementary Table 3), transcriptome-wide neural- and patient-specificity scores derived from three scRNA-Seq datasets (Supplementary Table 4), and *in silico* COSTAR drug screening results across 1,120,823 compounds (Supplementary Table 6).

## Code Availability

Code for de-multiplexing of DRUG-Seq data can be <u>found on GitHub</u>. COSTAR code and example data is available at: https://www.snijderlab.org/COSTAR. Additional code is available on reasonable request.

## Acknowledgements

We thank the patients and their families for making this study possible. We thank core facilities at ETH Zurich and University of Zurich, including the Functional Genomics Center Zurich (FGCZ), the Flow Cytometry Core Facility (EPIC, FCCF), ScopeM, and central informatics services (ID). This project has received funding from the European Research Council (ERC) under the European Union’s Horizon 2020 research and innovation programme (Grant agreement No. 803063-SCIPER; BS), and from the Personalized Health and Related Technologies (PHRT) Strategic Focus Area of the ETH Domain (Project #2021-566; BS). We further acknowledge funding from the Swiss National Science Foundation (grant no. 310030_204972, MAH; 310030_185155, MW), and financial support from the Department of Biology as well as the Institute of Molecular Systems Biology of the ETH Zurich. We thank the research and administrative staff who helped us throughout the study, including Yasmin Festl and Silas Krämer, and members of the Snijder lab for discussions.

## Extended Data Figure Legends

**Extended Data Fig. 1:**
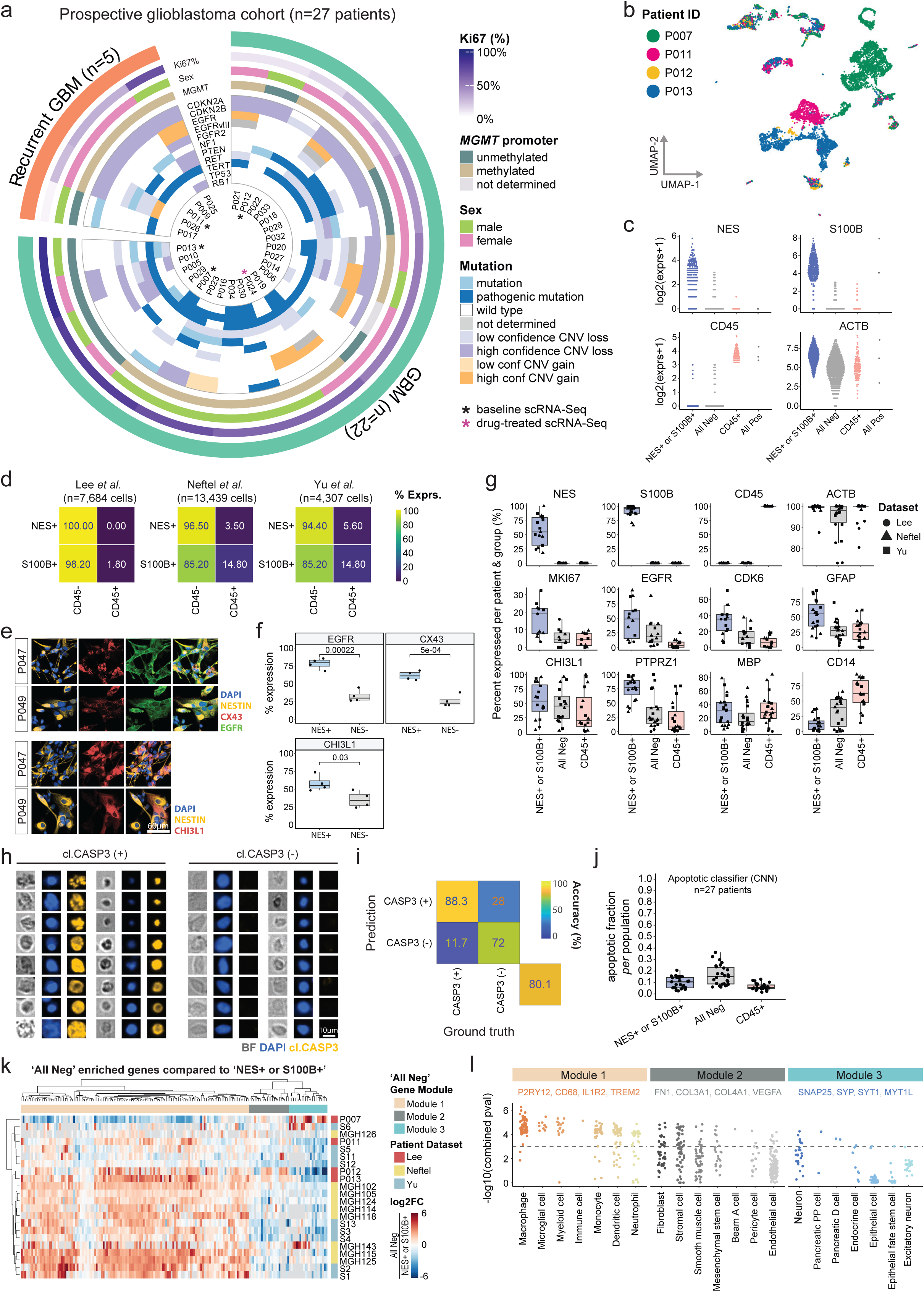
Glioblastoma prospective cohort overview and single-cell RNA-sequencing of four patient samples. **a**, Circos plot overview of the glioblastoma prospective patient cohort (n=27 patients) included in this study. Concentric circles from outermost to innermost show newly diagnosed versus recurrent tumour status, Ki67 labeling index, sex, *MGMT* promoter methylation status, and the most frequent genetic alterations (n=11) determined by targeted next-generation sequencing (NGS). Asterisks (*) denote scRNA-Seq patient samples (black, n=4 patients at baseline; pink, n=1 patient after drug treatment). CNV; copy number variation. *See Supplementary Table 1,2 for full cohort information*. **b**, UMAP projection of 7684 single-cell transcriptomes from four glioblastoma patient samples colored by patient (P007, 3475 cells; P011, 1490 cells; P012, 330 cells; P013, 2389 cells, referred to as ‘Lee *et al*., this study’). **c**, scRNA-Seq log2(expression+1) of glioblastoma markers (NES, S100B), pan-immune marker (CD45), and housekeeping gene (ACTB) of n=1,320 single cells as a subset of **b,** (n=330 randomly sampled cells per patient) **d**, Percent of Nestin or S100B positive cells (rows) either negative or positive for CD45 (columns) by scRNA-Seq across 22 glioblastoma patient samples and 3 scRNA-Seq datasets (Lee *et al.*, this study, n=4 patients, n=7,684 cells; Neftel *et al.*, n=9 patients, n=13,519 cells; Yu *et al.*, n=9 patients, n=4,307 cells). **e,** Representative immunofluorescence (IF) images of two glioblastoma patient samples labeled with different glioblastoma markers (Nestin, EGFR, CX43, and CHI3L1). **f,** Quantification of IF images in **e,** across n=4 glioblastoma patient samples. *P*-values from a two-sided t-test are shown. **g**, Percent of cells expressing key marker genes (y-axis) per patient (data points) and subpopulation (x-axis) across 22 glioblastoma patient samples (dots) and 3 scRNA-Seq datasets (shape) as in **d**. **h**, Example single-cell crops of cleaved CASP3+/-negative cells by IF in the image dataset used to train a convolutional neural network (CNN) based on nuclear (DAPI) and cell morphology (Brightfield) to detect apoptotic cells. **i**, Performance of the trained apoptotic classifier CNN in classifying the manually curated test image dataset consisting of CASP3+/-single-cell crops (n=1,214 images) into the corresponding classes. Cell classification accuracy shown as a confusion matrix. **j**, Fraction of cells classified as apoptotic by the CNN across the prospective patient cohort (n=27 patients) and marker defined populations. **k**, Genes (columns) enriched in (NES-, S100B-, and CD45-) triple-negative cells (‘All Neg’ cells) compared to ([NES+ or S100B+] and CD45-) cells across the 22 patients (rows) of the three scRNA-seq cohorts (row annotation color). Heatmap depicts log2(fold change) of genes enriched in the ‘All Neg’ cells. Expression of top-10 genes (columns) per patient (rows) clustered into 3 gene modules (Modules 1-3; column annotation color). **l**, Cell-type specific enrichment analysis (Web-CSEA, Dai et al. 2022) of the ‘All Neg’ enriched gene modules as in **k**. Dots represent individual Web-CSEA datasets, example genes that are members of their respective gene modules are annotated above. **g,f,l,** Boxplots as in *Fig. 1c*.

**Extended Data Fig. 2:**
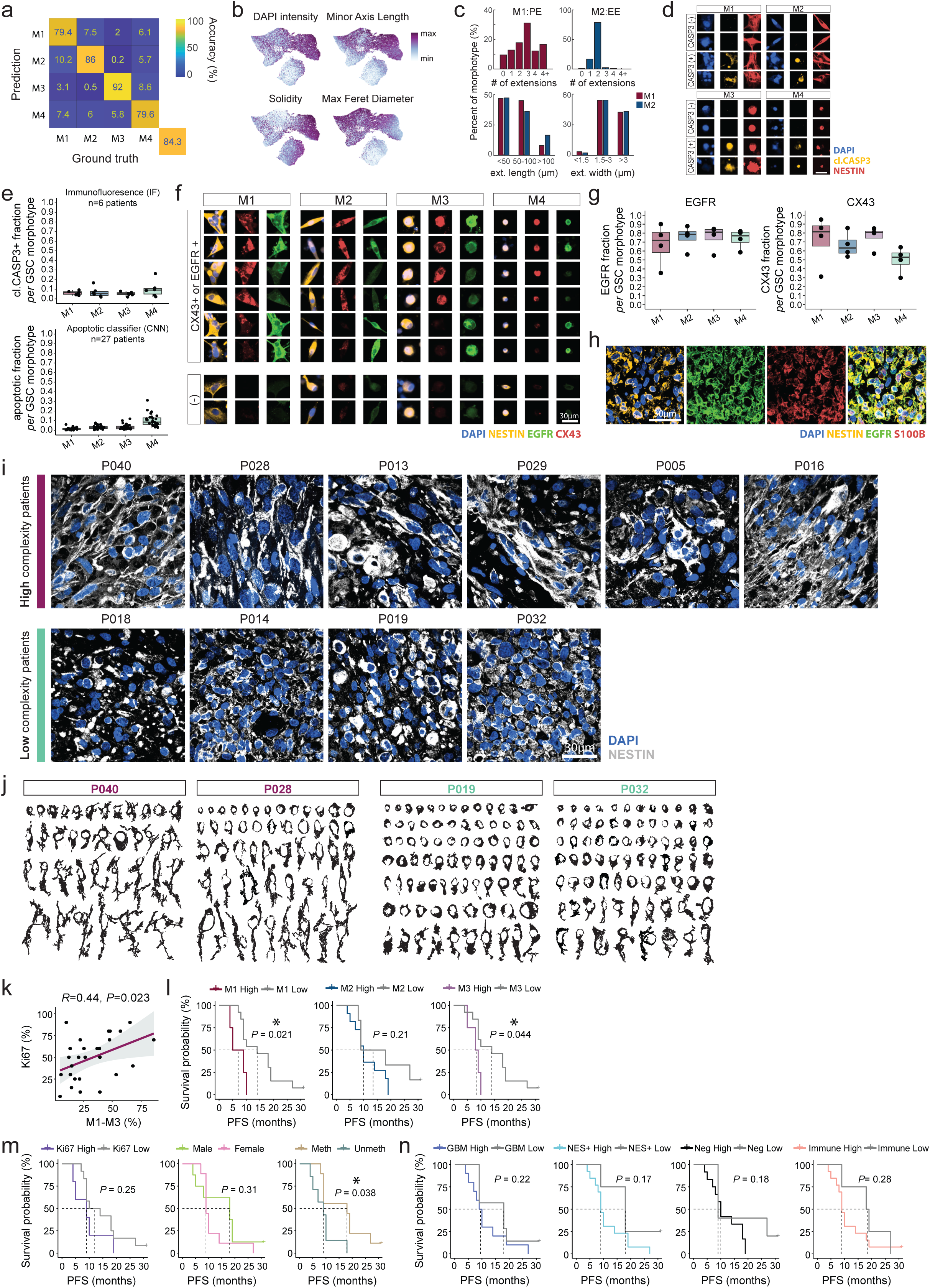
Deep learning of glioblastoma stem cell morphologies and clinical parameter-based stratification of patient survival. **a**, Performance of the trained GSC morphology CNN in classifying the manually curated test image dataset consisting of Nestin+ single-cell crops (n=10,204 images) into the corresponding four GSC morphotypes (M1-M4). Cell classification accuracy shown as a confusion matrix. **b,** UMAP projection of the morphological CNN feature space of 84,180 single cells (up to n=1,000 cells per morphotype and patient, n=27 patients). Cells are colored by the local median of selected single-cell image-based features as in *Fig. 1g*. **c,** Frequency of extensions per cell (top panels) in extension-containing morphotypes M1 (n=180 cells) and M2 (n=264 cells), and quantification of maximum extension length and extension width in M1 (n=111 cells) and M2 (n=127 cells) morphotypes (bottom panels). **d,** Example single-cell IF images of cleaved CASP3+/-negative cells across M1-M4 Nestin+ GSC morphotypes. Scale bar, 30µm. **e**, Fraction of apoptotic cells across M1-M4 Nestin+ GSC morphotypes quantified by either cleaved CASP3 IF (top; n=6 patients) or by the apoptotic CNN classifier (bottom; n=27 patients). **f,** Example single-cell IF images of NES+ and CX43+/EGFR+ (top) or NES+ and CX43-/EGFR-(bottom). **g,** Fraction of EGFR+ (left) or CX43+ (right) cells per Nestin+ GSC morphotype (x-axis) across four patients (dots). **h,** Example *in situ* immunohistochemistry (IHC) image of tumour region from patient P014 stained with nuclear stain (DAPI), glioblastoma stem cell marker (Nestin), epidermal growth factor (EGFR), and astrocyte lineage marker (S100B). **i,** Example *in situ* IHC images of tumour regions from patients with high *ex vivo* GSC morphotype complexity (top panels; n=6 patients) and low *ex vivo* GSC morphotype complexity (bottom panels; n=4 patients). **j,** Examples of manually segmented individual Nestin+ cells from binarized IHC images (Nestin channel) of individual patients confirming the presence and spectrum of M1-M4 morphotypes *in situ.* Two high morphotype complexity patients (left; red labels; P040, P028) and two low morphotype complexity patients (right; green labels; P019, P032) shown. **k,** Correlation of histopathological Ki67 labeling (y-axis) index with percent of GSCs with an M1-M3 morphotype (x-axis) per patient. Linear regression line (dark blue) with a 95% confidence interval (light grey). Pearson correlation coefficient and *P*-value shown. **l-n,** Kaplan-Meier survival curves of progression-free survival (PFS) in newly diagnosed glioblastoma patients (n=17 patients) stratified by **l,** M1-M3 morphotype abundance (high, low) among Nestin+ GSCs; **m,** histopathological Ki67% labeling index, sex, and *MGMT* promoter methylation status (n=1 patient with undetermined *MGMT* status omitted); **n,** IF marker-defined population abundances per sample as defined in *Fig. 1d.* **l-n,** Survival curves are compared using the log-rank (Mantel-Cox) test. Tick mark indicates ongoing response. **e,g,** Boxplots as in *Fig. 1c*.

**Extended Data Fig. 3:**
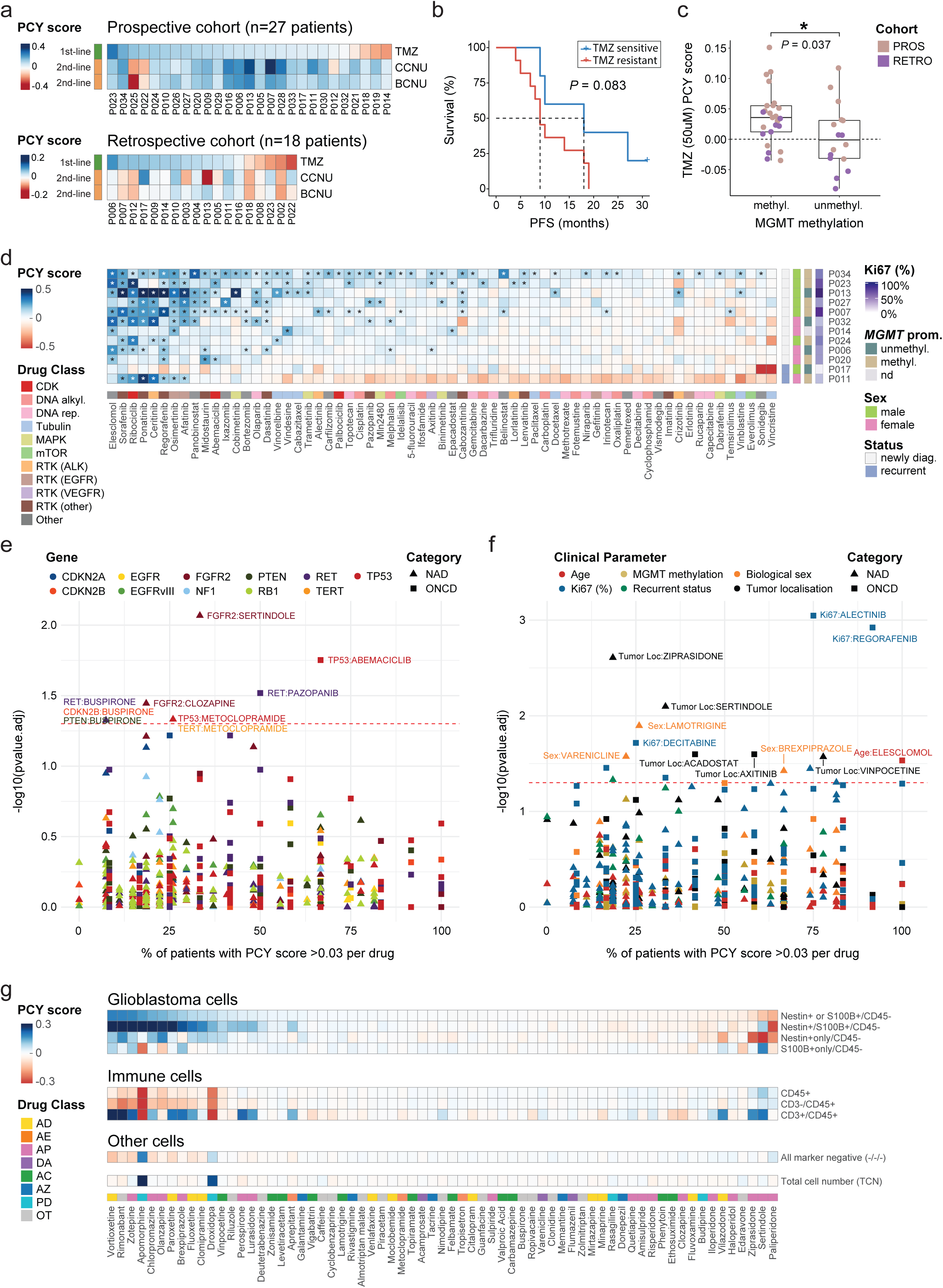
Patient *ex vivo* drug response relates to clinical parameters, tumour composition, and mutational profiles. **a**, Glioblastoma drug (GSDs; rows; n=3 drugs) response across patient samples (columns; prospective cohort, PROS, n=27 patients; retrospective cohort, RETRO, n=18 patients). GSD response is averaged across four concentrations for Temozolomide (TMZ; 1st-line chemotherapy; 50, 100, 250, 500µM) and Lomustine/Carmustine (CCNU and BCNU respectively; 2nd-line chemotherapies; 10, 50, 100, 250µM). **b,** Stratification of progression-free survival (PFS) of newly diagnosed glioblastoma patients (prospective cohort; n=16 patients) based on based on mean *ex vivo* Temozolomide sensitivity (TMZ PCY score) of (Nestin+/S100B+ and CD45-) cells (blue, sensitive; red, resistant). Kaplan-Meier survival curves are compared using the log-rank (Mantel-Cox) test and the optimal TMZ PCY score cut-point to stratify patients is determined by maximally selected rank statistics. Tick mark indicates ongoing response. **c,** Temozolomide (50µM) *ex vivo* response of glioblastoma patients (dots; n=41 patients across both cohorts) stratified by *MGMT* promoter methylation status. Unmethyl; unmethylated, Methyl; methylated. Wilcoxon rank sum test, *P*=0.037. Boxplots as in *Fig. 1c*. **d,** Drug response matrix of oncology drugs (ONCDs; columns; n=65 drugs) across glioblastoma patient samples (rows; n=12 patients). Clinical annotations per patient sample (rows) indicate the Ki67 labeling index, *MGMT* promoter methylation status (unmethyl; unmethylated, methyl; methylated, nd; not determined), Sex, and recurrent tumour status (Status). Column drug annotations indicate oncology drug class as in *Fig. 2d*. Asterisks (*) denote FDR-adjusted *P* < 0.05. **e,** Pharmacogenomic analysis of the most common genetic alterations (n=11) in glioblastoma patients and *ex vivo* drug response (PCY score). Each datapoint represents a [gene:drug] association, where x-axis denotes the percent of patients for which the respective drug’s PCY score >0.03 and the y-axis denotes FDR-adjusted *P*-values. **f,** As in **e**, but for associations between clinical parameters and *ex vivo* drug response (PCY score). **e,f** Colored by gene/clinical parameter and shape denote drug category. Red dashed line indicates the significance threshold. *P*-values were calculated using the Wilcoxon rank sum test for two groups, and for three or more groups, by the Kruskal-Wallis test. For ONCD associations, the following genetic mutations or clinical parameters had less than 3 patients in any category and were thus not analyzed: Genetic, *EGFRvIII*, *NF1*, *TERT*, *RB1*; Clinical, Recurrent status. **g,** Drug response matrix of neuroactive drugs (NADs, n=67 drugs; classes annotated as in *Fig. 2f*) averaged across glioblastoma patient samples (n=27 patients) for each cell population defined by immunofluorescence markers (Nestin, S100B, CD3, and CD45) and total cell number (TCN). **a,d,g,** Heatmap color scale indicates the PCY score of **a,d,** Nestin+ or S100B+ cells **g,** mean PCY score of each respective population averaged across patients. Outliers beyond color scale limits were correspondingly set to minimum and maximum values.

**Extended Data Fig. 4:**
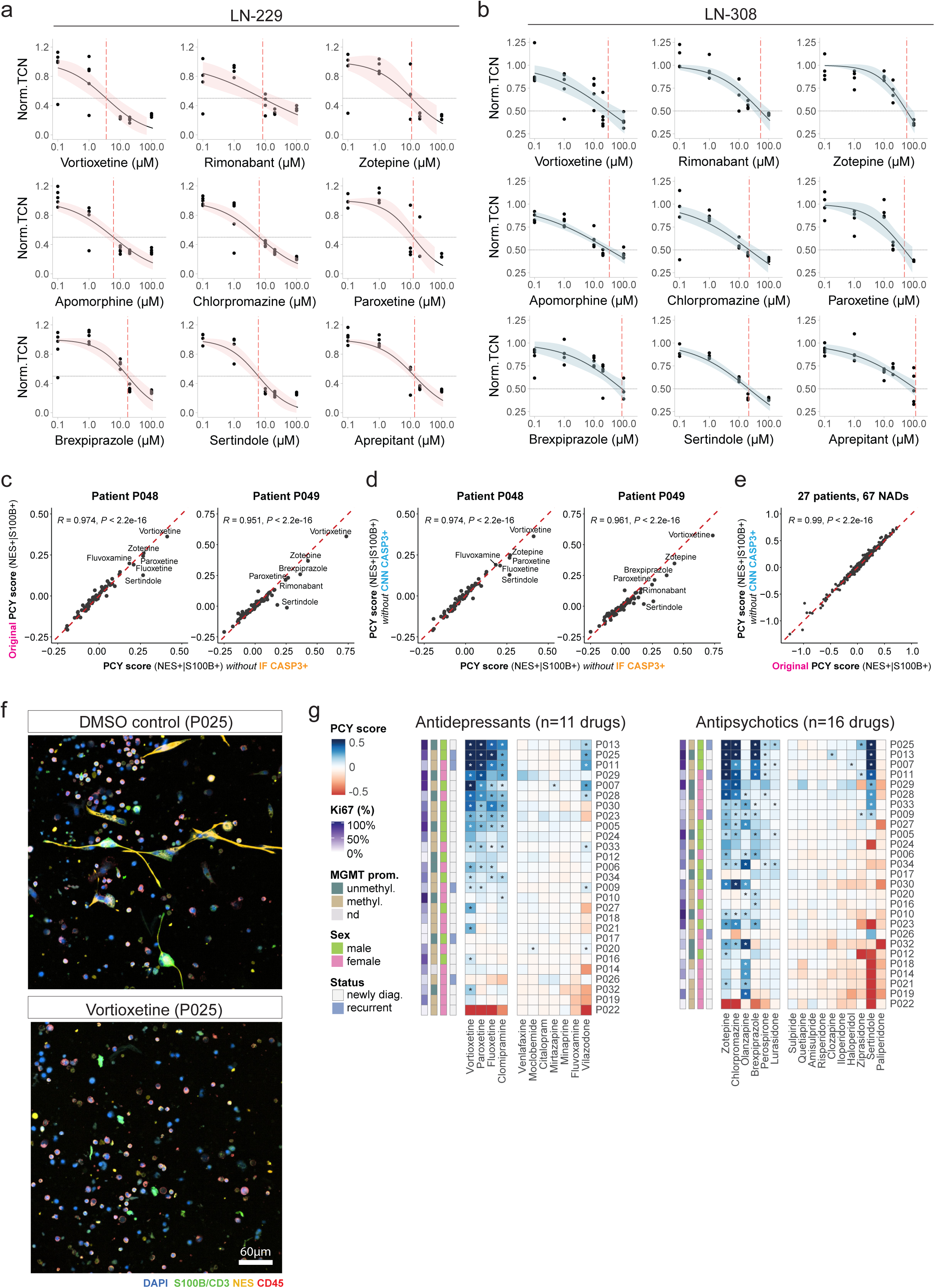
Robust and dose-dependent drug responses to neuroactive drugs across glioblastoma cell lines and patient samples. **a,b,** Dose-response curves of glioblastoma cell lines (LN-229/LN-308; *see also Supplementary Fig.S4 for spheroid lines ZH-161/ZH-562*) of a subset of top neuroactive drugs (n=9 drugs) across different concentrations (logarithmically spaced x-axis, n=5 concentrations). Y-axis denotes relative cell counts normalized to DMSO control. Dose-response curves (solid lines) are fitted with a two-parameter log-logistic distribution with 95% confidence intervals (colored per cell line) and ED50 (red dashed lines). n=3-5 replicate wells/drug, n=15 DMSO wells. **a,** Dose-response curves of glioblastoma cell line LN-229. **b,** Dose-response curves of glioblastoma cell line LN-308. **c-e**, Comparison of neuroactive drug pharmacoscopy scores of (Nestin+/S100B+ and CD45-) glioblastoma cells (n=67 NADs; original PCY score) to NAD PCY scores calculated by excluding cleaved CASP3+ apoptotic cells. Apoptotic cells are defined either by immunofluorescence (PCY score without IF CASP3+) or by the apoptotic CNN classifier (PCY score without CNN CASP3+; see also *Methods*). Pearson correlation coefficients with *P*-values annotated. **c,d**, NAD screens performed in two validation patient samples (P048, P049). **c**, Comparison of the original PCY score to the PCY score without IF CASP3+ **d**, Comparison of the PCY score without IF CASP3+ the PCY score without CNN CASP3+ **e**, Comparison of the original PCY score to the PCY score without CNN CASP3+ across the whole prospective cohort (n=27 patients) and neuroactive drugs (n=67 drugs). **f,** Representative immunofluorescence images of a patient sample (P025) at baseline (DMSO control; top) and treated with Vortioxetine (bottom). Glioblastoma cells are labeled with the nuclear stain DAPI, astrocyte lineage marker S100B, and neural progenitor marker Nestin while immune cells are labeled with the T-cell marker CD3 and pan-immune marker CD45. Scale bar, 60µM. **g,** Drug response matrix of antidepressants (left, n=11 drugs) and antipsychotics (right, n=16 drugs) across glioblastoma patient samples (n=27 patients) subsetted from the original matrix, as shown in *Fig. 2f*.

**Extended Data Fig. 5:**
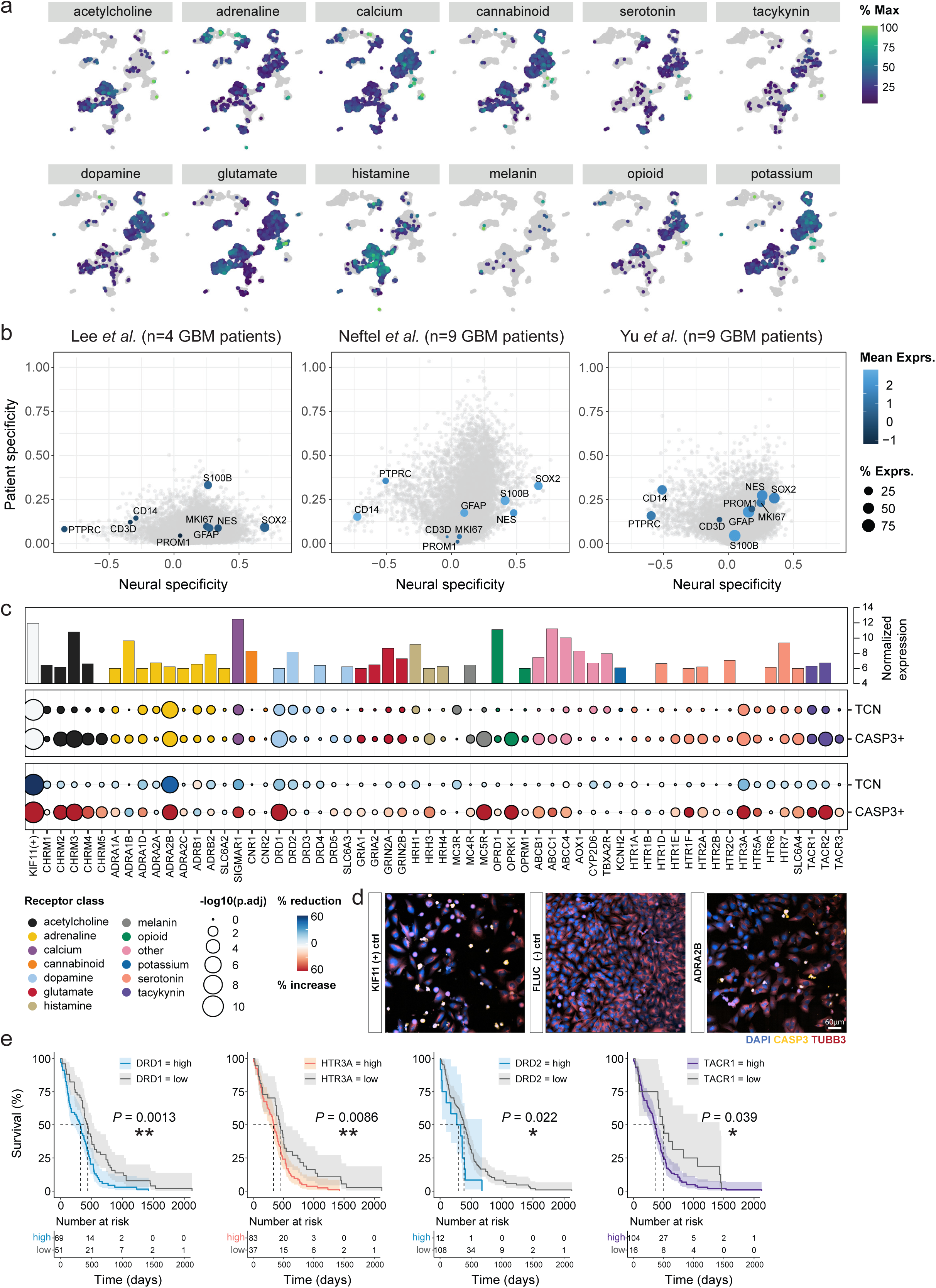
Single-cell heterogeneity and functional dependencies of primary neuroactive drug targets. **a**, UMAP projection of 7684 single-cell transcriptomes from four glioblastoma patient samples (P007, P011, P012, P013), colored by aggregate scRNA-Seq expression across primary target genes (PTG) per receptor class in Fig. 3b. Color scaled to percent of maximum expression per receptor class. **b**, Neural specificity score (x-axis) versus patient specificity score (y-axis) for three independent glioblastoma scRNA-Seq datasets. Each dot represents a gene, with key marker genes annotated with labels. Key marker genes colored by mean detected expression across cells and dot size scales with percent of expressed cells. All other detected genes are colored in grey. (Lee et al., this study; n=4 patients, n=7684 cells, n=15,668 genes; Neftel et al., n=9 patients, n=13,519 cells, n=22160 genes; Yu et al., n=9 patients, n=4307 cells, n=19,098 genes; see also Methods). **c**, Baseline RNA-Seq expression (top panel; y-axis) and siRNA-mediated gene silencing of PTGs in LN-229 cells (n=59 siRNA conditions; columns; middle and bottom panels). Total cell number reduction (TCN) and cleaved CASP3+ fraction increase (CASP3+) relative to the (-) control FLUC siRNA condition depicted as a circle per gene. Circle sizes scale with the –log10(FDR-adjusted P value). Color represents either the receptor class of each PTG (middle) or the total cell number (TCN; bottom) for each tested PTG. **d**, Representative immunofluorescence images of siRNA-mediated gene silencing of the positive control gene (KIF11 (+) ctrl; left), negative control gene (FLUC (-) ctrl; middle), and ADRA2B (right). Scale bar, 60µM. Cells are stained for DAPI (blue), cleaved CASP3 (yellow) and TUBB3 (red). **e**, Kaplan-Meier survival analysis and associated risk tables of the TCGA primary glioblastoma cohort (n=120 patients) based on RNA-Seq expression of 4 PTGs (panels) that significantly reduce cell viability in c, and stratify patient survival. Optimal cut-point for patient stratification (high, low) is determined by maximally selected rank statistics. Survival curves are compared using the log-rank (Mantel-Cox) test. 95% confidence intervals are indicated in shaded curves.

**Extended Data Fig. 6:**
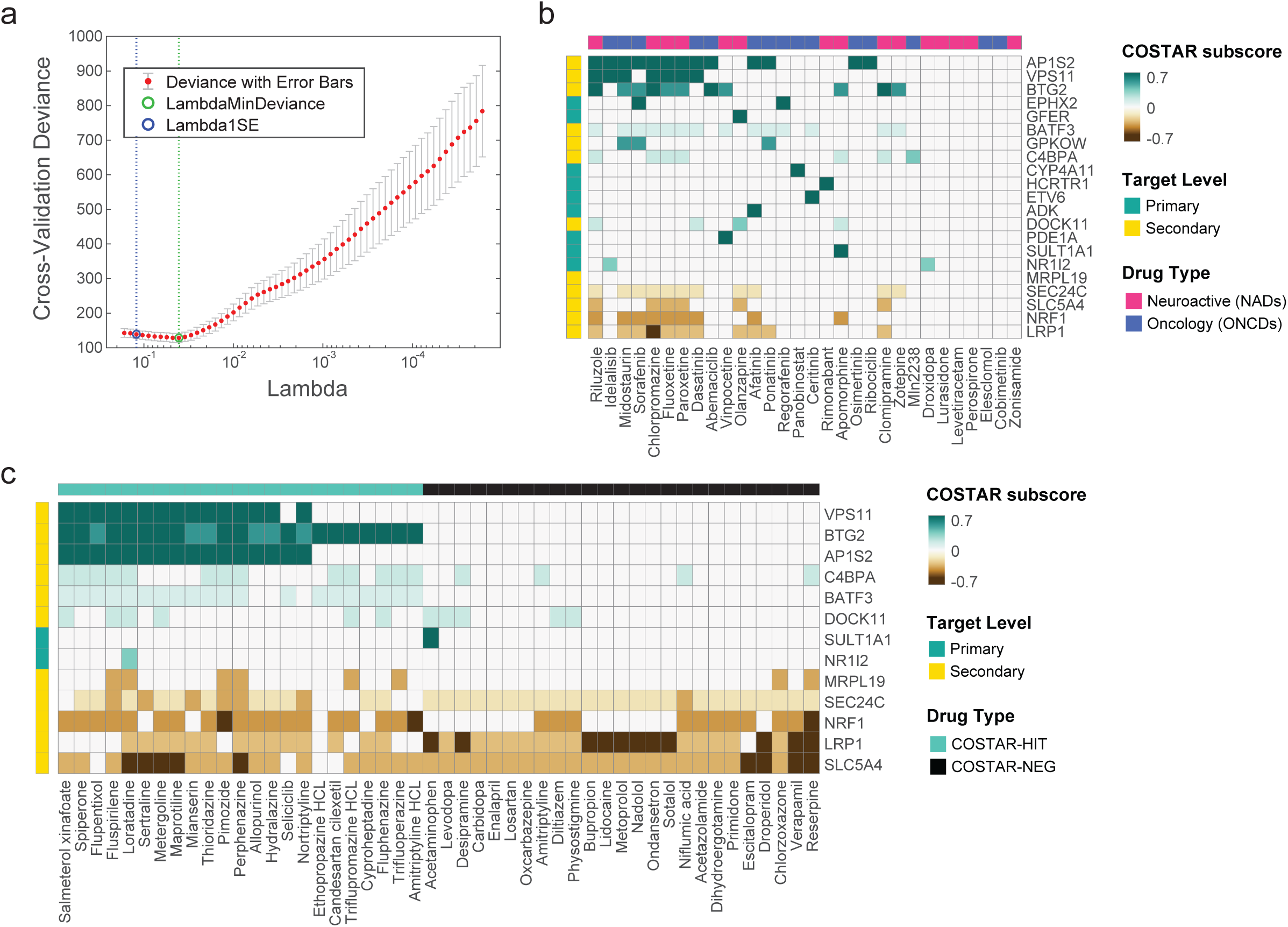
COSTAR identifies a drug-target connectivity signature predictive of anti-glioblastoma efficacy. **a**, Visualization of the local optimum in the cross-validated predictive power of COSTAR LASSO regression when fitting a binomial model to predict drug activity by PCY (hit vs neg) based on a drugs connectivity pattern (COSTAR constellation, shown in Fig. 3f). X-axis denotes the Lambda regularization parameter and the y-axis denotes the cross-validated error of the model (deviance). Red dots and light grey error bars indicate the average and standard deviation in deviance across 20 bootstrapped runs. Vertical dashed lines and colored circles indicate either the Lambda value with the minimal mean squared error (green, MSE) or the more conservative Lambda value with minimal MSE plus one standard deviation (blue, MSE+1STD). **b**, COSTAR subscores of PCY-hit drugs that were part of the COSTAR training data (columns; n=30 drugs) to primary and secondary drug targets (rows). **c**, COSTAR subscores of COSTAR-predicted drugs that were chosen for experimental validation in glioblastoma patient samples (columns; n=23 COSTAR-HIT drugs; n=25 COSTAR-NEG drugs) to primary and secondary drug targets (rows). **b-c,** Heatmap color scale indicates the COSTAR subscore which is the LASSO model coefficient multiplied by the integrated connectivity of drug to target mapping. Target genes with absolute COSTAR LASSO coefficients >0.1 are displayed. Target level (primary or secondary target) is annotated per gene on the left.

**Extended Data Fig. 7:**
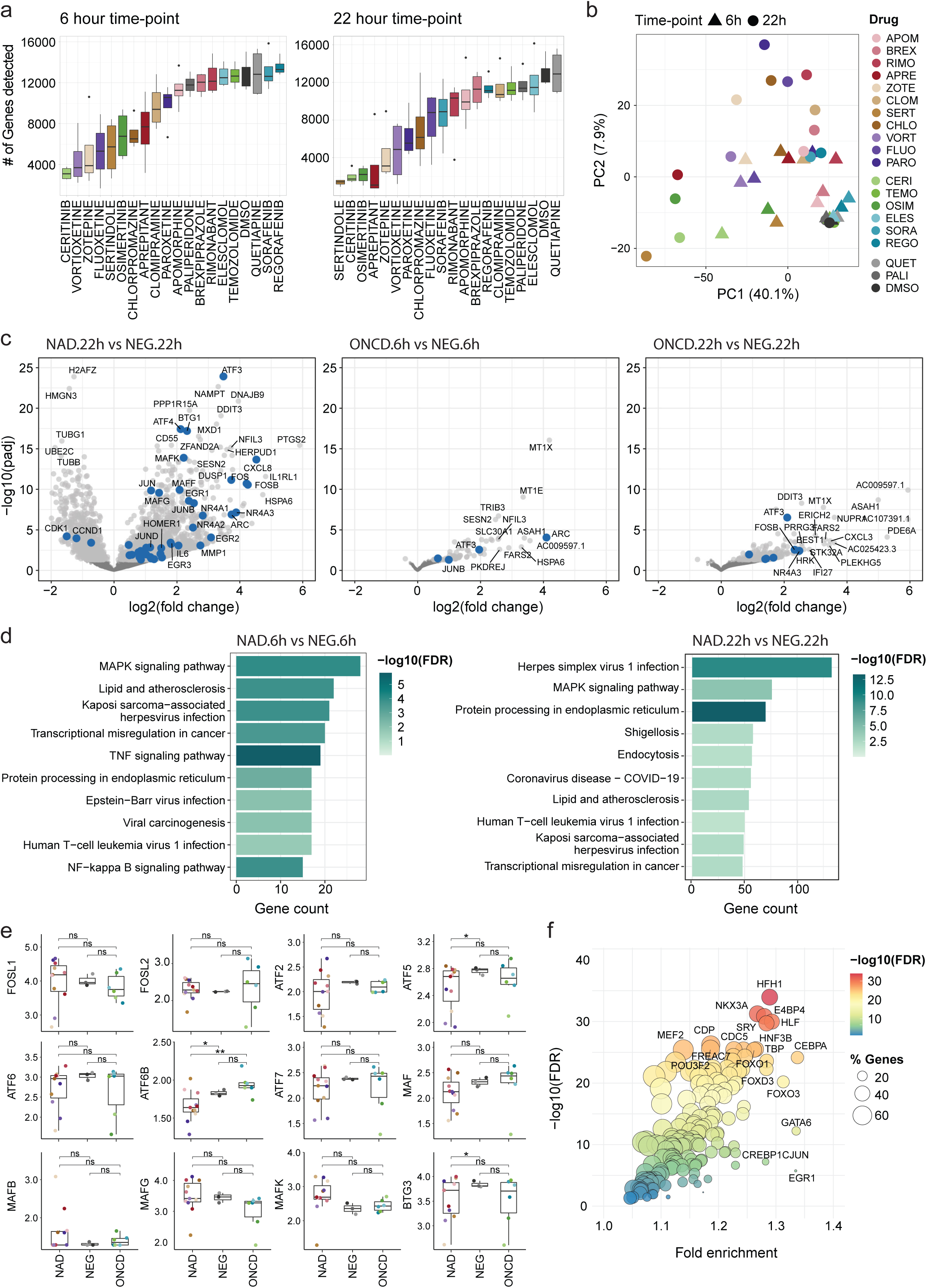
DRUG-Seq reveals a consistent transcriptional response to neuroactive drugs with anti-glioblastoma efficacy. **a**, Number of genes detected by DRUG-Seq (y-axis) per drug condition (columns) and by time-point n=20 drugs, n=2 time-points, n=4 replicates per drug/time-point. **b**, Principal component analysis (PCA) of averaged RNA-Seq counts per drug (color) and time-point (shape). **c**, Comparisons of drug induced transcriptional profiles by DRUG-Seq shown as (log2(fold change) versus –log10(adjusted *P*-value)) for NADs vs NEGs (22h, left), ONCDs vs CTRLs (6h, middle), and ONCDs vs CTRLs (22h, right). Genes above a –log10(0.05 adjusted P-value) threshold (light grey) and non-significant genes (dark grey) are shown. Highlighted genes (blue) include AP-1 transcription factor (TF) network genes (PID AP1 PATHWAY^81^) and key COSTAR signature genes. **d**, Top enriched KEGG terms for differentially expressed genes based on DESeq2 comparisons of NADs vs NEGs (6h, left) and NADs vs NEGs (22h, right). Bars represent the number of differentially expressed genes present in the annotation, and colors indicate –log10(false discovery rate). **e**, Expression of AP-1 transcription factor family and BTG genes additional to *Fig. 4d*. Visualization and statistical tests as in *Fig. 4d*. *P*-values: not significant (ns) P > 0.05, *P < 0.05, **P < 0.01. **f**, Transcription factor binding site enrichment analysis of genes that were upregulated in NAD treated cells in *Extended Data* Fig. 7c (22h, left). Circles correspond to transcription factor annotations, circle sizes scale with the fraction of genes present in the annotation, and colors indicate –log10(false discovery rate). **a,e** Boxplots as in Fig. 1c.

**Extended Data Fig. 8:**
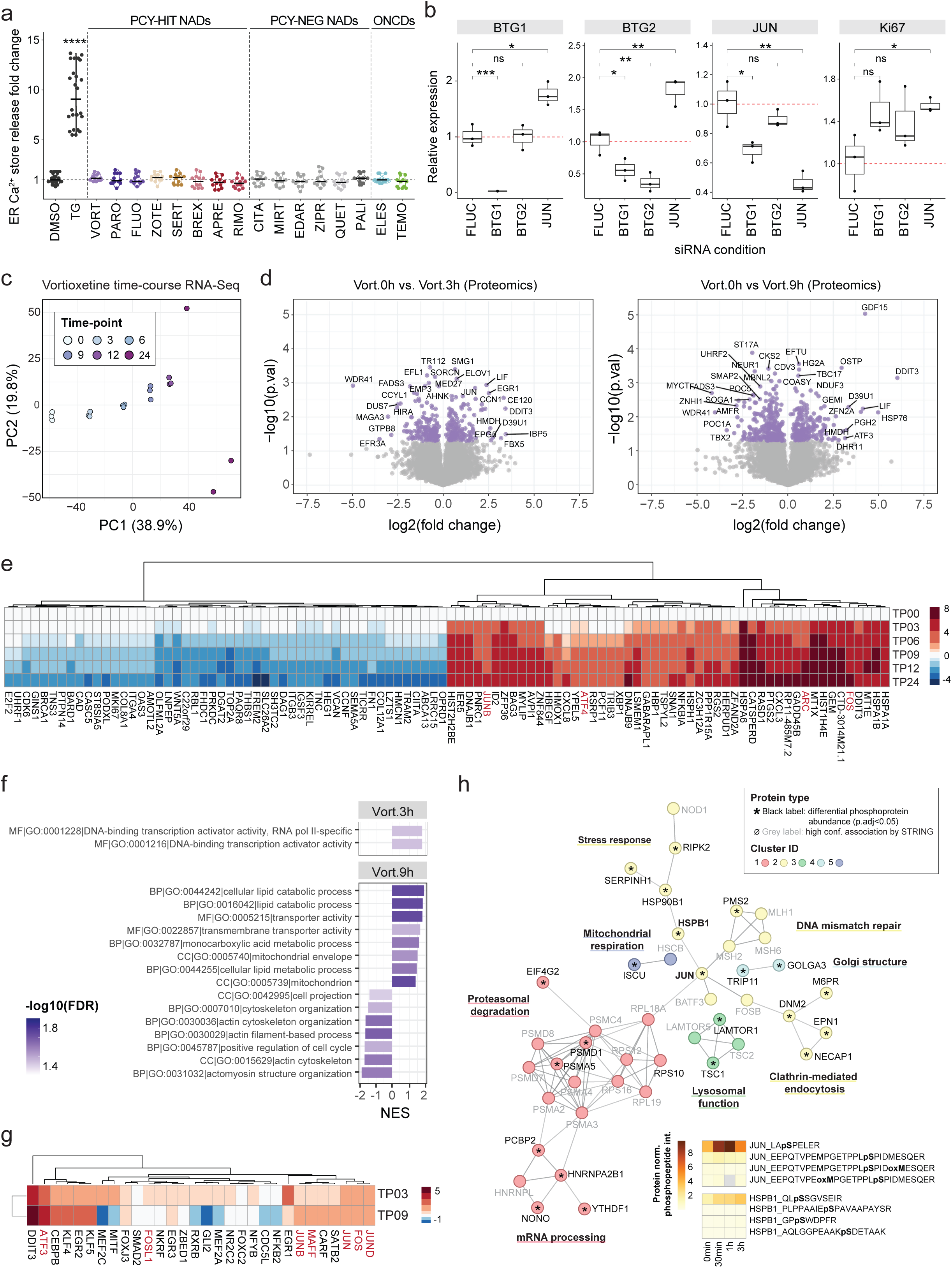
Measuring ER calcium store release, siRNA-mediated silencing of COSTAR signature genes, and Vortioxetine-induced transcriptomic, proteomic and phosphoproteomic response. **a**, ER calcium store release upon drug treatment relative to DMSO vehicle control (fold change, 190-430 seconds interval) measured by FLIPR assays in LN-229 cells (n=4 assay plates; n=18 conditions; n=12 wells/drug; DMSO and Thapsigargin (TG) positive control, n=24 wells each). Different drug categories including PCY-hit NADs, n=8 drugs; PCY-neg NADs, n=6 drugs; PCY-hit ONCDs, n=2 drugs; and TG. Two-sided t-test against DMSO vehicle control. *P*-values adjusted for multiple comparisons by Holm correction. *P*-values: TG, 2.86e-16. ****P < 0.0001. Line indicates the median value. **b**, Relative gene expression of BTG1, BTG2, JUN and Ki67 (panels) upon siRNA knockdown (columns) normalized to the FLUC negative control siRNA (n=3 biological replicates; black dots). Boxplots as in *Fig. 1c*. **c**, Principal component analysis (PCA) of replicate-averaged RNA-Seq counts following Vortioxetine treatment (20µM) in LN-229 cells (n=3 replicates/time-point) colored by time-point. **d**, Time-point comparisons (left, 3 hours vs 0 hours; right, 9 hours vs 0 hours) of proteomics measurements following Vortioxetine treatment (Vort, 20µM; n=3 replicates/condition) in LN-229 cells shown as volcano plots of log2(fold change) versus –log10(*P*-value). Proteins above a –log10(0.05 P-value) threshold are colored in purple. **e**, Heatmap of log2(fold change) in gene expression per time-point (rows; relative to 0h) for the top 100 genes (columns) contributing to PC1 in *Extended Data* Fig. 8c. TP; time-point. AP-1 transcription factors and AP-1 effector genes in red. **f**, Gene Ontology (GO) gene set enrichment analysis of signed –log10(P-values) of comparisons from *Extended Data* Fig. 8d. Bars represent the normalized enrichment score (NES) and colors indicate –log10(false discovery rate). **g**, Log2(fold change) in protein expression per time-point (rows; relative to 0h) for the proteins (columns) contributing to enriched GO term “GO:0001216 DNA-binding transcription activator activity” in *Extended Data* Fig. 8f. AP-1 transcription factors are labeled in red. **h**, Connected protein-protein interaction network of differentially abundant phosphoproteins upon Vortioxetine treatment (20µM; n=3 replicates/condition) in LN-229 cells at any time-point. 22 out of 67 connected and significantly enriched phosphoproteins are shown (asterisks; black labels) with high confidence STRING protein interactions (grey labels). Cluster IDs (node colors) are based on the MCL algorithm with annotated biological pathways. Heatmap depicts protein abundance-normalized phosphopeptide (rows) intensities of JUN and HSPB1 across time-points (columns). Both genes are also significantly upregulated at the transcript level across all time-points.

**Extended Data Fig. 9:**
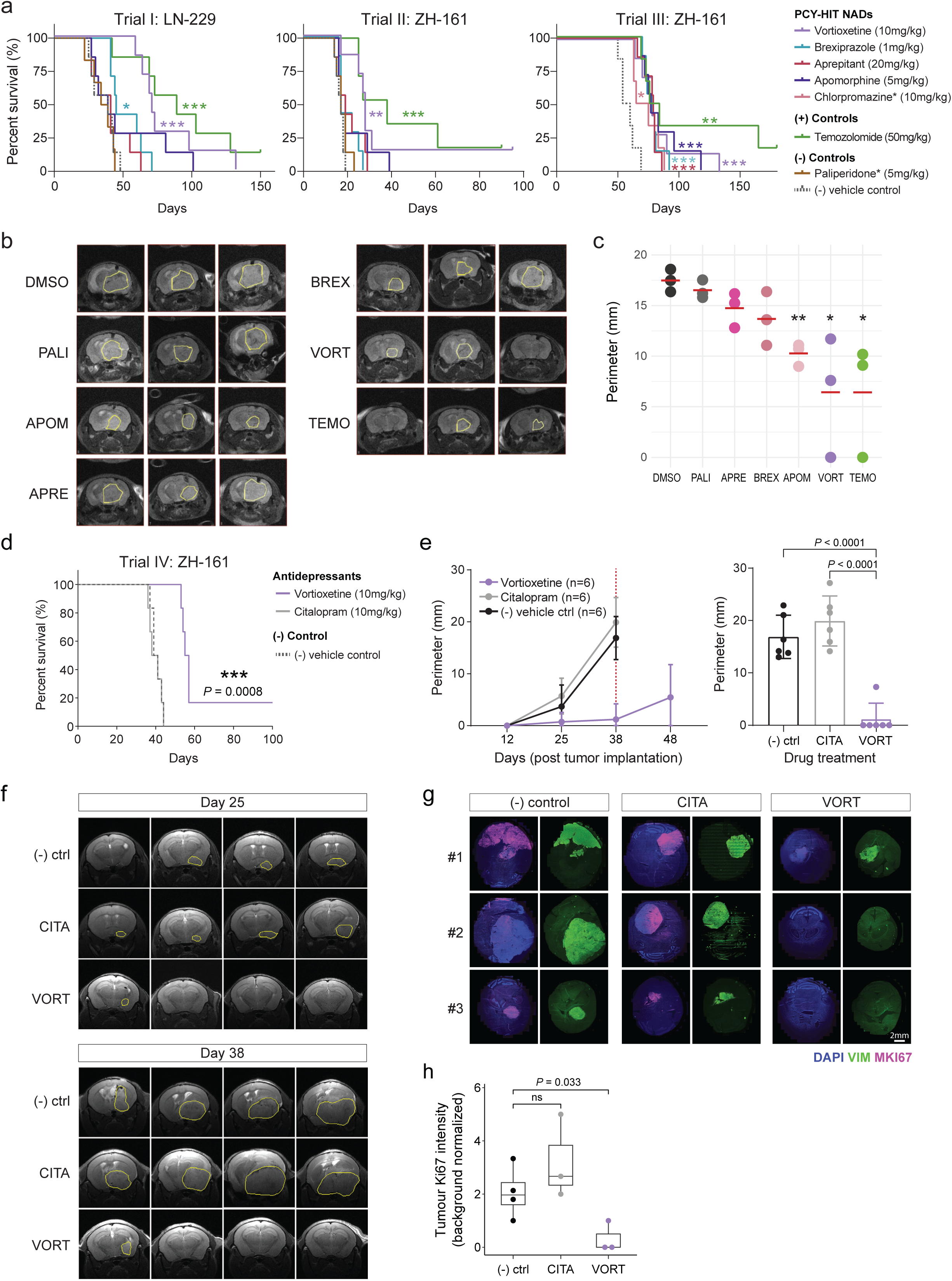
Top neuroactive drugs confer a significant survival benefit in orthotopic *in vivo* mouse models of glioblastoma. **a**, Complete survival analysis across three independent *in vivo* trials: Trial I: LN-229, Trial II: ZH-161, and Trial III: ZH-161, each with n=6-7 tumour-bearing mice per drug treatment group and n=7 drug treatments per trial. Mice were treated with their respective drugs for each trial intraperitoneally (*i.p.*) between days 5-21 after tumour implantation. PCY-HIT NADs: Vortioxetine (VORT; 10mg/kg; Trial I, *P*=0.0001; Trial II, *P*=0.0016; Trial III, *P*=0.0006); Brexpiprazole (BREX; 1mg/kg; Trial I, *P*=0.0249; Trial II, ns; Trial III, *P*=0.0002); Aprepitant(APRE; 20mg/kg; Trial I, ns; Trial II, ns; Trial III, *P*=0.0006); Apomorphine (APOM; 5mg/kg; Trial I, ns; Trial II, ns; Trial III, *P*=0.0005); Chlorpromazine(CHLO; 10mg/kg; Trial III, *P*=0.011). Positive control (+): Temozolomide (TMZ; 50mg/kg; Trial I, *P*=0.0009; Trial II, *P*=0.0002; Trial III, *P*=0.0011). PCY-NEG NAD: Paliperidone (PALI; 5mg/kg; Trial I, ns; Trial II, ns), and a negative vehicle control. Drug names with asterisk (*) denote drugs used in a subset of the three *in vivo* trials. Survival plotted as Kaplan–Meier curves and *P*-values calculated using log-rank (Mantel-Cox) test. Censored mice denoted as tick marks. **b,** Representative MRI images of three ZH-161 transplanted mice (columns) after 15 days per drug treatment (n=7 drugs). Tumour perimeters are indicated in yellow. **c,** Quantification of tumour perimeters corresponding to *Extended Data* Fig. 9b. Dots represent the perimeter in mm (y-axis) for individual mice per drug (columns), red lines indicate mean value. Two-sided t-test. *P*-values: Apomorphine (APOM; *P*=0.0014); Vortioxetine (VORT; *P*=0.034); Temozolomide (TMZ; *P*=0.0284). *P*-values: not significant (ns) P > 0.05, *P < 0.05, **P < 0.01. **d,** Survival analysis of *in vivo* Trial IV: ZH-161-iRFP720 comparing the efficacy of the antidepressant Vortioxetine (10mg/kg; *P*=0.0008) with a PCY-NEG antidepressant Citralopram (10mg/kg) and a negative vehicle control (n=6 mice/treatment group). Drug treatment schedule and statistical analysis was performed as in *Extended Data* Fig. 9a. **e,** Quantification of tumour perimeters of drug-treated mice in **d,** at multiple time-points (left panel; n=6 mice/treatment group) post tumour implantation by MRI. Right panel illustrates individual data points (mice) at day 38 post-implantation. **f,** Representative MRI images of four ZH-161-iRFP720 transplanted mice (columns) after 25 days (top) and 38 days (bottom) of drug treatment (n=3 drugs). Tumour perimeters are indicated in yellow. **g,** Representative immunohistochemistry images of horizontal sections of mouse brains (n=3 mice/treatment group) stained with human-specific Ki67 and Vimentin (VIM). **h,** Quantification of Ki67 tumour intensity normalized to background per treatment group (n=3-4 mice analyzed per treatment).

**Extended Data Fig. 10:**
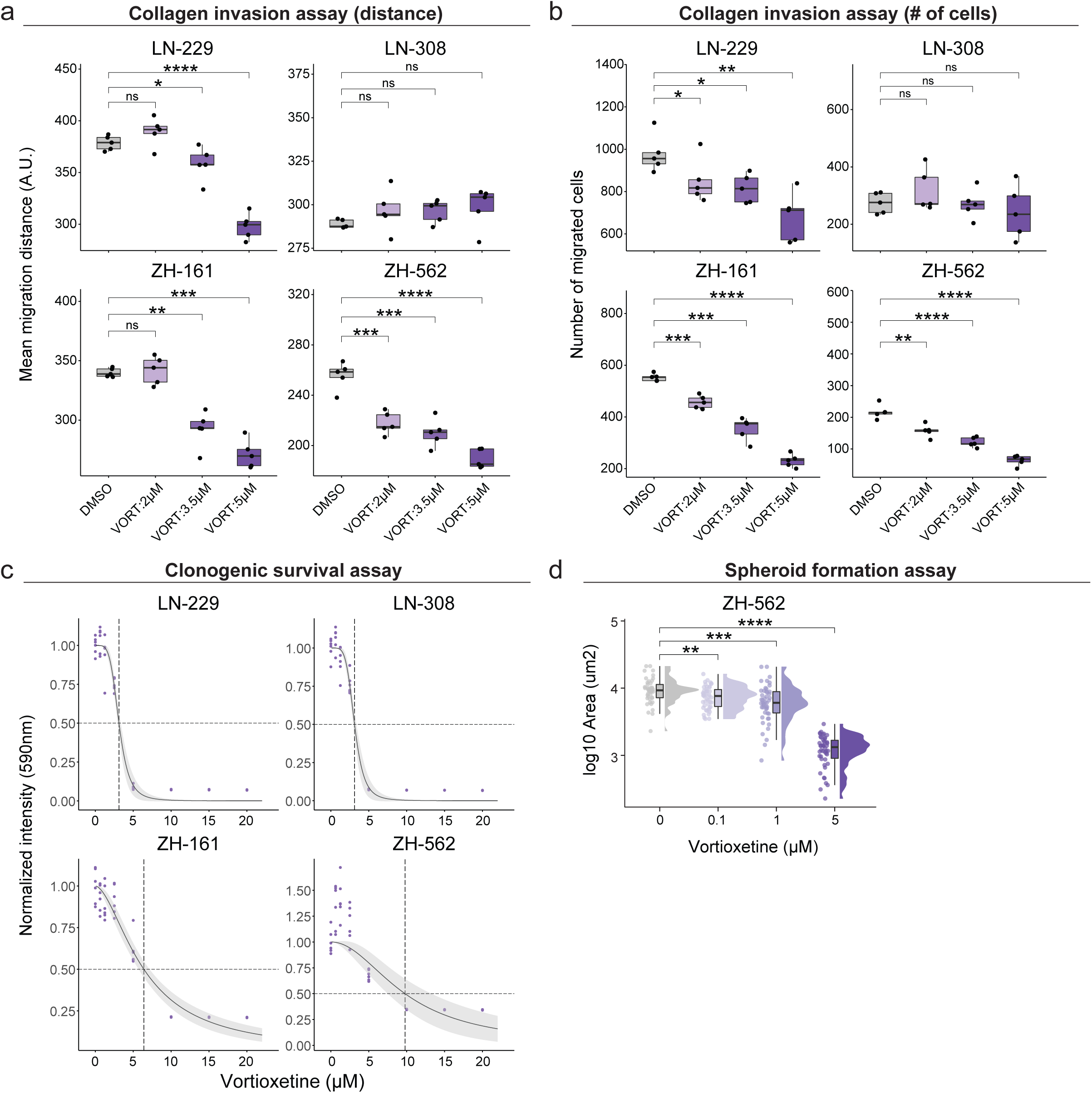
The multi-faceted anti-glioblastoma effects of Vortioxetine. **a**, Mean cell migration distance per condition (n=5 replicate wells) and **b,** number of migrated cells measured in a collagen-based spheroid invasion assay after 36 hours of Vortioxetine treatment (2, 3.5, 5µM) across four glioblastoma cell lines; LN-229 (n=560-1125 cells/well), LN-308 (n=137-426 cells/well), ZH-161 (n=200-574 cells/well), ZH-562 (n=38-253 cells/well). **c,** Clonogenic survival measured by a resazurin-based cell viability assay after 11-13 days of Vortioxetine treatment (7 concentrations; 0.625-20µM, n=6 replicate wells/concentration, n=50-500 cells/well) across four glioblastoma cell lines; LN-229 LN-308, ZH-161, ZH-562. Dose-response fitted with a two-parameter log-logistic distribution with 95% confidence intervals (light grey) and ED50 (dashed lines). **d,** Spheroid formation analyzed by the 2D area of the ZH-562 line measured after 12 days of Vortioxetine treatment (0.1-5µM). Approximately 5 cells/well initially seeded. DMSO; 0µM, n=45 replicate wells; 0.1µM, n=46, *P*=0.005; 1µM, n=47, *P*=0.00027; 5µM, n=46, *P*<0.0001. Data is shown per concentration as a boxplot, individual data points, and histogram. Boxplots as in *Fig. 1c*.

## Methods

### Patient sample processing

Surgically removed tumours were collected at the University Hospital of Zurich (Universitätsspital Zürich, USZ) with approval by the Institutional Review Board, ethical approval number KEK-StV-Nr.19/08, BASEC number 2008-02002. Metadata of the prospective and retrospective glioblastoma patient cohorts including clinical parameters, experiment inclusion, and genetics summary can be found in *Supplementary Table 1*. The prospective cohort consists of patients where fresh tissue was processed directly within 4 hours after surgery (n=27 patients for drug screening, plus an additional n=9 patients for validation experiments). For progression-free-survival (PFS) analysis of the prospective cohort, only patients with newly diagnosed glioblastoma that received concomitant Temozolomide (TMZ) chemotherapy were included. The retrospective cohort (n=18 patients) consists of patients for which snap-frozen bio-banked tissue was available. All retrospectively studied patients included received maintenance TMZ with overall-survival (OS) documented as a clinical endpoint. Retrospective samples were selected to cover a broad spectrum of progression-free survival outcomes, and were further selected based on quality control measures including cell viability, cell number, and the amount of debris present in the sample.

#### Patient sample dissociation for ex vivo drug screening

Tissue samples were first washed with PBS and cut into small pieces using single-use sterile scalpels. Subsequent dissociation was performed in reduced serum media (DMEM media; #41966 with 2% FBS; #10270106, 1% Pen-strep; #15140122, and 25mM HEPES; #15630056, all products from Gibco) supplemented with Collagenase IV (1mg/ml) and DNaseI (0.1mg/ml) using the gentle MACS Octo Dissociator (Miltenyi Biotec, 130-096-427) for maximally 40 minutes. Homogenates were filtered through a 70µm Corning cell strainer (Sigma-Aldrich, #CLS431751) and washed once with PBS containing 2mM EDTA. Myelin and debris removal was performed by a gradient centrifugation of the cell suspension in a 7:3 mix of PBS:Percoll (Sigma-Aldrich, #P4937), with an additional PBS wash. In case the cell pellet visibly contained a significant portion of red blood cells (RBCs), RBC lysis was performed with 1X RBC lysis buffer (Biolegend, #420301) at room temperature for 10 minutes prior to the PBS wash. Subsequently, cells were resuspended in reduced serum media, filtered once more through a 70µm Corning cell strainer, and counted using the Countess II Automated Cell Counter (Invitrogen). In case sufficient cell numbers remained after cell seeding for *ex vivo* drug testing (see also ‘Pharmacoscopy’ methods), cells were cryopreserved in 10% DMSO-containing cryopreservation media, and/or cultured in DMEM media supplemented with 10% fetal bovine serum, 1% Pen-strep, and 25mM HEPES to obtain low-passage patient-derived cell lines (PDCs) maintained as adherent cell cultures.

#### Patient in situ tissue characterization by H&E staining and immunohistochemistry (IHC)

Patient tissue samples obtained directly from surgery were fixed using 4% formalin and embedded in paraffin. The paraffin embedded tissue was sliced in 5 µm sections. Deparaffinized tissue sections were subsequently stained with hematoxylin (Artechemis AG, Switzerland) and eosin (Sigma-Aldrich, USA) using the Tissue-Tek Prisma automated staining system. Imaging of H&E stained patient tissue sections (*Supplementary Data Fig. S3*) was performed by bright-field imaging at 40x using the Pannoramic 250 Slide Scanner (3DHISTECH). Based on nuclei and cell morphologies observed in the H&E images, non-necrotic tumour regions were annotated and confirmed by a board-certified neuropathologist (D.K., E.R.).

For immunohistochemistry (IHC) of patient tissue sections, deparaffinization and rehydration was performed using an automated continuous linear stainer (Medite, COT20) containing xylene and serial dilutions of EtOH (100%, 95%, 70%). Tissue sections were immediately subjected to heat-mediated antigen retrieval at 95°C for 10 minutes in 1X pH6.0 sodium citrate buffer (Sigma-Aldrich, #21545) using Micromed T/T Mega Multifunctional Microwave Rapid Histoprocessor (Milestone). Tissue sections were subsequently fixed with 4% PFA (Sigma-Aldrich, #F8775) in PBS, blocked in 5% FBS and 0.1% Triton containing PBS, and stained overnight at 4°C in blocking solution with the following primary antibodies and dilutions: anti-Nestin (1:150, Biolegend, #656802, clone 10C2), anti-EGFR (1:300, Abcam, #ab98133), anti-S100 beta antibody (1:300, Abcam, #ab215989, clone EP1576Y). Secondary antibodies used include the following: donkey anti-sheep IgG (H+L) cross-adsorbed secondary antibody, Alexa Fluor™ 488 (Thermo Scientific, #A11015), goat anti-mouse IgG (H+L) highly cross-adsorbed secondary antibody, Alexa Fluor™ Plus 555 (Thermo Scientific, #A32727), goat anti-rabbit IgG (H+L) highly cross-adsorbed secondary antibody, Alexa Fluor Plus 647 (Thermo Scientific, #A32733).

Confocal imaging of tissue sections was performed at 100x magnification with the Nikon Spinning Disk SoRa microscope where z-stack images compiling 31 series of 0.2 μm z-steps were taken. Maximum intensity projection of z-stacks were obtained by ImageJ and single cells were segmented based on their nuclei (DAPI channel) and cytoplasm (binarized Nestin channel) using CellProfiler 2.2.0. Downstream image analysis was performed with MATLAB R2019a-R2020a based on ‘Nuclei’ and associated ‘Cell’ features from CellProfiler. Nestin+ cells were filtered based on the intensity histogram of the Nestin channel. Manual tracing of single-cell morphologies (Extended Data Fig. 2j) was performed in Adobe Illustrator from binarized Nestin channel images by creating vectorized objects using the ‘Image Trace’ tool and visually verified traces selected for display purposes.

### Cell culture

The adherent human glioblastoma cell lines LN-229 (ATCC, #CRL-2611) and LN-308, and patient-derived cell cultures (P022.C, P024.C, P030.C) were cultured in Dulbecco’s modified Eagle medium (DMEM, #41966, Gibco) supplemented with 10% fetal bovine serum (FBS, #10270106, Gibco). Adherent human glioblastoma cell lines and patient-derived cell cultures were passaged using Trypsin-EDTA (0.25%, Gibco, #25200056). For DRUG-seq, RNA-Seq, siRNA knockdown, and proteomics measurements using LN-229 cells, low-passage cells below passage 15 were used. The spheroid human glioblastoma-initiating cell lines ZH-161 and ZH-562 was generated from freshly isolated tumour tissue and cultured in Neurobasal medium (NB, #21103049, Gibco) supplemented with B27 (Gibco, #17504044), 20 ng/mL b-FGF (Peprotech, #AF-100-18B), 20 ng/mL EGF (Peprotech, #AF-100-15), 2 mM L-glutamine (Gibco, #25030081). ZH-161 and ZH-562 cells were passaged using Accutase (Stemcell Technologies, #07920). Glioblastoma cell lines were authenticated at the Leibniz Institute DSMZ (Braunschweig, Germany) and regularly tested negative for mycoplasma. All cell cultures were maintained at 37°C, 5% CO_2_ in a humidified incubator.

### Pharmacoscopy (drug testing, immunocytochemistry, confocal microscopy, image analysis)

The method of ‘*pharmacscopy*’ refers to high-content image-based drug testing including the following steps of cell seeding, drug testing, immunocytochemistry, confocal microscopy, image analysis, and pharmacoscopy score calculation for each tested drug.

#### Cell seeding and drug testing

To perform pharmacoscopy of glioblastoma patient samples, freshly dissociated cells from resected tissue (see also methods relating to ‘*Patient sample processing*’) were seeded into clear-bottom, tissue-culture treated, CellCarrier-384 Ultra Microplates (Perkin Elmer, #6057300) with 0.5-1.5×10^4 cells/well, typically within 4 hours of surgery. For cultured glioblastoma cell lines and patient derived cell cultures, trypsinized or accutase-treated cells were seeded at 0.5-2.5×10^3 cells/well in 384 well plates. Prior to cell seeding, drugs were re-suspended as 5mM stock solutions and dispensed into 384 well plates using an Echo 550 liquid handler (Labcyte) at their respective concentrations in a randomized plate layout to control for plate effects. Detailed information regarding drugs used in this study can be found in *Supplementary Table 3*. Different drug libraries included glioblastoma drugs (GSDs, n=3 drugs), oncology drugs (ONCDs, n=65 drugs), and neuroactive drugs (NADs, n=67 drugs). GSDs were tested at the following concentrations: Temozolomide (TMZ; 1st-line glioblastoma chemotherapy; 50, 100, 250, 500µM) and Lomustine/Carmustine (CCNU and BCNU respectively; 2nd-line glioblastoma chemotherapies; 10, 50, 100, 250µM). All ONCDs were tested at 1 and 10μM concentrations. All NADs were tested at 20μM and for select NADs (*Extended Data Fig. 4a,b* and *Supplementary Data Fig. S4*) a concentration range of 0.1-100μM was tested. Drug plates included the following number of replicate wells per drug/concentration: GSD plate, drug, n=3 wells, DMSO, n=16 wells; NAD plate, drug, n=4 wells, DMSO, n=16-24 wells; ONCD plate, drug, n=4 wells, DMSO, n=16 wells. Cells were incubated for 48 hours with drugs in reduced serum media at 37°C, 5% CO_2_ before proceeding to cell fixation.

#### Immunocytochemistry

Cells were fixed with 4% PFA (Sigma-Aldrich, #F8775) in PBS and blocked in 5% FBS and 0.1% Triton containing PBS. For characterization of cellular and morphological composition across glioblastoma patient samples, cells were stained overnight at 4°C in blocking solution with the following antibodies and dilutions: Alexa Fluor® 488 anti-S100 beta (1:1000, Abcam, #ab196442, clone EP1576Y), PE anti-Nestin (1:150, Biolegend, #656806, clone 10C2), Alexa Fluor® 488 anti-CD3 (1:300, Biolegend, #300415, clone UCHT1), Alexa Fluor® 647 anti-CD45 (1:300, Biolegend, #368538, clone 2D1) and DAPI (1:1000, Biolegend, #422801, stock solution 10mg/ml). Due to the manufacturer discontinuation of the Alexa Fluor® 488 anti-S100 beta antibody, from patient sample P030 and onwards in the prospective cohort, samples were either stained with a self-conjugated Alexa Fluor® 488 anti-S100 beta antibody, where Alexa Fluor™ 488 NHS Ester (Thermo Scientific, #A20000) was conjugated to the anti-S100 beta antibody (Abcam, #ab215989, clone EP1576Y) or the following antibody panel where the 488 and 555 channel markers were swapped: Alexa Fluor® 488 anti-Nestin (1:150, Biolegend, #656812, clone 10C2), Alexa Fluor® 555 anti-S100 beta (1:1000, Abcam, #ab274881, clone EP1576Y), PE anti-CD3 (1:300, Biolegend, #300441, clone UCHT1), Alexa Fluor® 647 anti-CD45 (1:300, Biolegend, #368538, clone 2D1).

Other conjugated antibodies used in this study include the following: Alexa Fluor® 647 anti-Tubulin Beta 3 (1:1000, Biolegend, #657406, clone AA10), Alexa Fluor® 488 anti-Vimentin (1:500, Biolegend, #677809, clone O91D3), Alexa Fluor® 555 anti-Cleaved Caspase-3 (1:500, Cell Signaling Technology, #9604S), Alexa Fluor® 546 anti-HOMER (1:300, Santa Cruz Biotechnology, #sc-17842 AF546, clone D-3), PE anti-CFOS (1:300, Cell Signaling Technology, #14609S, clone 9F6), FITC anti-ATF4 (1:300, Abcam, #ab225332), Alexa Fluor® 488 anti-JUND (1:300, Santa Cruz Biotechnology, #sc-271938 AF488, clone D-9), Alexa Fluor® 594 anti-CD45 (1:300, Biolegend, #368520, clone 2D1).

Other unconjugated antibodies used in this study include the following: anti-Connexin43 (1:500, Cell Signaling Technology, #83649T), anti-EGFR (1:300, Abcam, #ab98133), anti-CHI3L1 (1:300, Cell Signaling Technology, #47066S, clone E2L1M), anti-Nestin (1:150, Biolegend, #656802, clone 10C2), anti-S100 beta antibody (1:300, Abcam, #ab215989, clone EP1576Y), anti-Ki67 (1:300, Cell Signaling Technology, #9129S, clone D3B5). For unconjugated primary antibodies, the following secondary antibodies were used: donkey anti-sheep IgG (H+L) cross-adsorbed secondary antibody, Alexa Fluor™ 488 (Thermo Scientific, #A11015), goat anti-mouse IgG (H+L) highly cross-adsorbed secondary antibody, Alexa Fluor™ Plus 555 (Thermo Scientific, #A32727), goat anti-rabbit IgG (H+L) highly cross-adsorbed secondary antibody, Alexa Fluor Plus 647 (Thermo Scientific, #A32733).

#### Confocal imaging and image analysis

Imaging of the 384 well plates was performed with an Opera Phenix automated spinning-disk confocal microscope (Perkin Elmer, HH14000000) at 20x magnification for all drug screening assays with the exception of 3D glioblastoma cell lines (ZH-161, ZH-562) where drug screening was performed at 10x magnification. Select images were imaged at 40x for visualization. Single cells were segmented based on their nuclei (DAPI channel) using CellProfiler 2.2.0. Downstream image analysis was performed with MATLAB R2019a-R2020a. Fractions of marker positive cells for each drug condition were derived for each patient sample based on the histograms of the local background corrected intensity measurements across the whole drug plate. In patient samples, marker positive populations were defined as following: glioblastoma cells ([Nestin+ or S100B+] and CD45-) and four Nestin+CD45-GSC morphotypes: M1 polygonal with extensions (M1:PE), M2 elongated with extensions (M2:EE), M3 round big cells (M3:RB), and M4 round small cells (M4:RS). Other cell types include immune cells (CD45+ and S100B-Nestin-) and marker-negative cells (S100B-Nestin-CD45-). Marker positive fractions were averaged across each well/drug. Deep learning analysis of Nestin+ glioblastoma stem cell (GSC) morphologies was performed at baseline (DMSO) and upon neuroactive drug (NAD) treatment for each patient, where the methodology is further detailed in the methods section ‘*Deep learning of glioblastoma stem cell morphologies*’.

#### Pharmacoscopy score calculation

The pharmacoscopy score (PCY score) quantifies the drug-induced relative reduction of any marker-or morphology-defined cell population by measuring the change of a defined target population upon drug treatment compared to DMSO (-) vehicle control. In patient samples, the PCY score is calculated based on the fraction of ([Nestin+ or S100B+] and CD45-cells) out of all cells, or the fraction of [Nestin+CD45-] GSC morphotypes (M1-M4) out of all cells. In patient-derived cell lines, the score is based on [Nestin+] cells out of all cells. By all cells, we refer to any detected cell with a DAPI+ nucleus.

**PCY score = 1** − **{ [TP_DRUG_]** ÷ **[TP_DMSO_] }**

where: **TP_DRUG_** = fraction of the target population in a given DRUG condition of all cells

**TP_DMSO_**= fraction of the target population in the DMSO control condition of all cells

PCY scores are averaged across technical replicates for each drug or control condition. For example, a positive PCY score of 0.5 can be interpreted as a drug-induced 50% reduction of the target population relative to the DMSO vehicle control. Additionally, in cases where the drug is cytotoxic and non-selectively kills every cell population equally or the drug does not exert an effect, the PCY score will be 0, and in cases where the target population proliferates upon drug-treatment, or exhibits higher toxicity to other cell populations other than the target population, the PCY score will be negative. In summary, a positive PCY score of 1 represents the strongest possible “on-target” response, a PCY score of 0 indicates no effect/equal cytotoxicity, and a negative PCY score indicates higher toxicity to other cell populations other than the defined target population. In cases where a target population is not defined as such for the glioblastoma cell lines, drug response and cell viability is measured as total cell number reduction in LN-229 and LN-308 lines and a reduction of 2D-projected total spheroid area in ZH-161 and ZH-562 lines.

#### Demonstration of pharmacoscopy score robustness to apoptotic cells

To experimentally validate the robustness of the PCY score to apoptotic cells, we performed complete *ex vivo* neuroactive drug (NAD, n=67 drugs) screens in two validation patient samples (P048 and P049) by explicitly staining the entire drug plate for cleaved CASP3 by immunofluorescence. Presented in *Extended Data* Fig. 4c-e, the drug response results show excellent reproducibility, both when comparing the original PCY scores with the PCY scores obtained after excluding CASP3+ cells by immunofluorescence, as well as when comparing the PCY scores after excluding CASP3+ cells defined either by IF or by the CNN apoptotic classifier (*see also Methods* ‘Image-based deep learning - Deep learning of apoptotic cell morphologies*’*). Finally, we re-calculated the PCY scores by excluding the CNN-classified apoptotic cells measured across all 27 patient samples and 67 neuroactive drugs and compared it to the original PCY scores reported in the manuscript (*Fig. 2f*). The drug response correlation with or without the inclusion of apoptotic cells was 0.988, demonstrating that the PCY score is highly robust to the presence of apoptotic cells (*Extended Data* Fig. 4e), and can be expected to be equally robust to other forms of cell death.

### Targeted Next Generation Sequencing (NGS, Oncomine Comprehensive Assay v3)

Tissue blocks from patient-matched glioblastomas were used to determine genetic alterations including mutations, copy number variations and gene fusions. Formalin-fixed paraffin-embedded (FFPE) tissue blocks were collected from the Tissue Biobank at the University Hospital Zurich (USZ). Tumour area was marked on the H&E slide and relative tumour cell content was estimated by a trained pathologist. 1-3 cores cylinders (0.6 mm diameter) from the tumour area of the FFPE blocks were used for DNA and RNA isolation. DNA was isolated with the Maxwell 16 FFPE Tissue LEV DNA Purification Kit (Promega, #AS1130). The double-strand DNA concentration (dsDNA) was determined using the fluorescence-based Qubit dsDNA HS Assay Kit. RNA was extracted with the Maxwell 16 FFPE Tissue LEV RNA Purification Kit (Promega, #AS1260). To avoid genomic DNA contamination, samples were pretreated with DNase1 for 15 min at room temperature (RT). Library preparation with 20 ng DNA or RNA input was conducted using the Oncomine Comprehensive Assay v3. Adaptor/barcode ligation, purification and equilibration was automated with Tecan Liquid Handler (EVO-100). NGS libraries were templated using Ion Chef and sequenced on a S5 (Thermo Fisher Scientific), and data were analyzed using Ion Reporter Software 5.14 with Applied Filter Chain: Oncomine Variants (5.14) settings and Annotation Set: Oncomine Comprehensive Assay v3 Annotations v1.4.

For NGS data analysis, the Ion Reporter Software within Torrent Suite Software was used, enabling detection of small nucleic variants (SNVs), copy number variations (CNVs), gene fusions and indels from 161 unique cancer driver genes. Detected sequence variants were evaluated for their pathogenicity based on previous literature and the ‘ClinVar’ database ^82^. Gene alterations described as benign or likely benign were not included in our results. Non-pathogenic mutations harboring a Minor Allele Frequency higher than 0.01 were not selected. The Default Fusion View parameter was selected. For CNV confidence range, the default filter was used to detect gains and losses using the confidence interval values of 5% confidence interval for Minimum Ploidy Gain over the expected value and 95% confidence interval for Minimum Ploidy Loss under the expected value. CNV low confidence range was defined for gain by copy number from 4 to 6 (lowest value observed for CNV confidence interval 5%:2.9) and loss from 0.5 to 1 (highest value observed for CNV confidence interval 95%:2.43). High confidence range was defined by gain up to 6 copy number (lowest value observed for CNV confidence interval 5%:4.54) and loss below 0.5 copy number (highest value observed for CNV confidence interval 95%:1.37). 5% and 95% interval confidence of all selected loss and gain are available in *Supplementary Table 2*. The minimum number of tiles required was eight. Results are reported as detected copy number.

### Single-cell RNA-Seq and re-analysis of other published datasets

#### Generation of cohort-matched scRNA-Seq datasets of glioblastoma patient samples

Cryopreserved single-cell suspensions of glioblastoma patient samples that were part of the prospective cohort were thawed in reduced serum media (DMEM containing 2% FBS) and used for single-cell RNA-Seq experiments. Viability markers SYTOX Blue (1 μM, Thermo Fisher, #S11348) and DRAQ5 (1 μM, Biolegend, #424101) were added to the cell suspension at least 15 minutes before sorting. FACS gates were set based on CD45 (Alexa Fluor® 594 anti-CD45, 1:20, Biolegend, #368520, clone 2D1), SYTOX Blue and DRAQ5 intensities to isolate live CD45+ and CD45-populations as shown in *Supplementary Fig. S1* using the BD FACSAriaTM Fusion Cell Sorter. Cells were sorted into DNA LoBind® Eppendorf tubes (VWR, #525-0130), then CD45-cells were mixed with CD45+ cells at 2:1 to 10:1 ratios depending on cell availability to enrich for glioblastoma cells. Single-cell transcriptomes from four patient samples (P007, P011, P012, P013; n=7684 cells) visualized in *Fig. 1b*, *Extended Data* Fig. 1b*, and Supplementary Fig. S1* are referred to as ‘Lee *et al.*; this study’. For patient sample P024 that was used to measure the effect of Vortioxetine drug treatment, FAC-sorted cells were incubated for 3 hours with or without 20µM Vortioxetine before proceeding to library preparation. Single-cell RNA-Seq library preparation was performed using the Chromium Next GEM Single Cell 3’ v3.0 and v3.1 kits (10x Genomics). Libraries were sequenced on the Novaseq 6000 (Illumina). Read alignment to the GRCh38 human reference genome, generation of feature-barcode matrices, and aggregation of multiple samples were performed using the Cell Ranger analysis pipeline (10x genomics, versions 3.0.1 and 6.1.1). Four patient samples (P007, P011, P012, P013) were processed in November 2019 with the earlier version of 10x Genomics library prep kits and Cell Ranger analysis pipeline while the later sample (P024) was processed in September 2021. Quality control for this in-house dataset was performed by only analyzing high-quality cells with fewer than 10% of mitochondrial transcripts and genes that had at least a count of 2 in at least 3 cells. For the Lee *et al.* dataset, an expression threshold of log2(count+1) > 3 was applied to consider a gene expressed in a given cell. Only patient samples with more than 50 positive cells for a given gene were considered in *Fig. 1c* and *Extended Data* Fig. 1g.

UMAP clusters in *Fig. 1b* are based on Leiden community detection and cell types are assigned by marker expression. Top marker genes per scRNA-Seq cluster in *Fig. 1b* that are expressed in more than 10 percent of cells per cluster are shown in *Supplementary Fig. S1b,c*. Glioblastoma (GBM) clusters are numbered in descending order based on cluster-averaged expression of the Gene Ontology term “stem cell differentiation” (GO:0048863).

#### Re-analysis of other published scRNA-Seq datasets

To analyze additional glioblastoma patient cohorts by single-cell RNA-Seq, we utilized two published datasets: (Neftel *et al.* 2019) and (Yu *et al.* 2020). For Neftel *et al.*, we removed cells with less than 2^9 detected genes and/or more than 15% of mitochondrial transcripts. For Yu *et al.* the data was already prefiltered, but patient samples (7-9, 14-15) that did not correspond to glioblastoma (grade IV astrocytomas) were not included. For both datasets only genes that had at a count of 2 in at least 2 cells were included in the analysis. For the Neftel *et al.* and Yu *et al.* datasets, expression thresholds of log2(count+1) over 5 and 3, respectively, were applied to consider a gene expressed in a given cell. Only patient samples with more than 50 positive cells for a given gene were considered in *Fig. 1c* and *Extended Data* Fig. 1g.

#### Cell-type specific enrichment analysis (CSEA) of gene modules enriched in Nestin/S100B/CD45-negative cells (‘All Neg’)

To determine putative cell types represented in Nestin-S100B-CD45-cells (‘All Neg’) by scRNA-Seq we analyzed the log2(fold change) of ‘All Neg’ enriched genes compared to ‘([Nestin+ or S100B+] and CD45-)’ glioblastoma cells. First, we created an aggregated average ‘metacell’ for each patient and subpopulation (either ‘All Neg’ or glioblastoma cells) by summing the counts across each patient-subpopulation and dividing this by the number of cells in the corresponding patient-subpopulation. This generated an aggregated average ‘All Neg’ metacell and glioblastoma metacell for each patient. Next, considering only genes where the aggregate-averaged expression is above 1 in at least one metacell type, we calculated the log2(fold change) of [‘All Neg’ metacell]/[glioblastoma metacell] per gene and per patient. Manhattan distance-based clustering of the top-10 log2(fold change) of ‘All Neg’ enriched genes per patient are visualized in *Extended Data* Fig. 1k across the three scRNA-Seq patient cohorts. Finally, dendrogram tree cutting of ‘All Neg’ enriched genes yielded gene modules (Modules 1-3) that were analyzed by WebCSEA (cell-type specific enrichment analysis; across 111 scRNA-Seq panels of human tissues and 1,355 tissue cell types; ^83^) to determine most likely cell types represented by the respective gene modules. The top-7 most likely cell types representing each ‘All Neg’ gene module ranked by the lowest combined p-values are shown in *Extended Data* Fig. 1l.

#### Neural and patient specificity score analysis

Neural specificity scores and patient specificity scores for each gene were defined as follows: using the in-house dataset, we identified putative cell types by unsupervised clustering using Monocle ^84^ and annotated the clusters based on known marker genes as being either immune or neural cells. We then obtained a list of differentially expressed genes between immune and neural cells using DESingle ^85^, using a logFC cutoff of 0.5. This yielded a list of 11571 neural-specific and 1157 immune specific genes. Using these lists as cell-type specific gene sets, we calculated an immune- and a neural score for each cell using singscore, and classified every cell in the additional datasets as either neural or immune based on a linear combination of both scores. To assess how specifically a gene is expressed in neural cells, we defined a ‘*neural specificity score’* as follows: [*neural specificity = fraction of neural cells expressing gene – fraction of immune cells expressing gene*] where we define expression of a gene in a cell as having any non-zero count. Thus, a positive score indicates that a gene is more often found in neural cells than in immune cells, and vice versa for negative scores. This score ranges from -1 (gene is expressed in all immune cells and no neural cells) to =1 (gene is expressed in all neural cells and no immune cells). Note that for low expressed genes, this score will be close to 0, reflecting the fact that we cannot make clear statements about cell type specificity for genes with expression values close to the detection limit of scRNA-Seq. To assess how much gene expression for a single gene varies across patients, we defined a ‘*patient specificity score’* as follows: First, for every gene *gi* and every patient *pj* we calculated a cell type composition independent fraction of cells expressing gene *gi* as [*Fraction_expressing_ij = fraction_expressing_immune_ij + fraction_expressing_neural_ij*]. We then defined patient specificity as the median absolute deviation (MAD) of fraction_expressing across all patients, thus defining [*Patient_specificity_i = mad(Fraction_expressing_i,:)*].

### Deep learning-based image analysis

#### Deep learning of glioblastoma stem cell (GSC) morphotypes

To generate a training dataset, Nestin+CD45-cells identified by immunofluorescence (IF) across the whole prospective glioblastoma patient cohort (n=27 patients) were cropped as 5-channel 150×150 pixel images. These single-cell image crops were then manually-curated and labeled as four morphological classes (M1-M4 GSC morphotypes) based on their shape, size, and presence of tumour extensions. A convolutional neural network (CNN) with a modified AlexNet architecture ^86^ with the number of output classes set to 4 was then trained on this manually-curated training data with 12,757 images per class and 51,028 images in total. CNN training included usage of the Adam optimizer, with a mini-batch size of 150 and a maximum number of 30 epochs. The initial learning rate was set to 0.001 with a piecewise learning rate schedule and a drop factor of 0.01 every 6 epochs.

Network performance on a manually-curated test image dataset consisting of 10,204 Nestin+ single-cell crops is shown as a confusion matrix in *Extended Data* Fig. 2a. All Nestin+ single-cell images were subsequently classified by this pre-trained CNN to determine GSC morphotype abundances across patients and drug conditions. For visualization of the CNN-based GSC morphotypes, UMAP plots were generated based on the CNN feature space that consists of ten dimensional activations taken from the 2nd-last fully connected layer of the network. The CNN feature space of 84,180 cells (maximally 1000 cells per class and patient, n=27 patients) was projected on the UMAP using the following parameters: distance metric, seuclidean; number of neighbors, 10; minimal distance, 0.06. Different morphological and marker-based features from the original cell segmentation determined by CellProfiler 2.2.0 such as cell area, eccentricity, and roundness, and mean marker intensity were selected for visualization.

#### Deep learning of multicellular IHC images

For deep learning of multicellular IHC images of DAPI- and Nestin-stained patient tissue sections a pre-trained version of the AlexNet CNN available in MATLAB R2023a was used. 50 full size IHC images (2304 x 2304 pixels; n=5 images/patient across 10 patients) at 100x magnification were automatically resized using an augmented image datastore and analyzed by this AlexNet for unsupervised clustering of image features. Dimensionality reduction on the activations from the last fully connected layer ‘fc8’ was performed by principal component analysis (PCA) as shown in *Fig. 1j*.

#### Deep learning of apoptotic cell morphologies

To generate a training dataset, across 6 glioblastoma patient samples, cleaved CASP3+/-cells by IF were cropped as 5-channel 50×50 pixel images. 6,072 single-cell image crops were then manually-curated and labeled as two classes (CASP3+ or CASP3-) based on their cleaved CASP3 staining. A convolutional neural network (CNN) with a modified AlexNet architecture ^86^ with the image input size set as 50×50×2 (2-channel classifier; including only the brightfield (BF) and DAPI channels) and number of output classes set to 2 (CASP3+/CASP3-) was then trained on this manually-curated image dataset (n=6,072 single-cell images; split by a 8:2 ratio into training and test data, respectively). CNN training included usage of the Adam optimizer, with a mini-batch size of 64 and a maximum number of 30 epochs. The initial learning rate was set to 0.01 with a piecewise learning rate schedule and a drop factor of 0.1 every 10 epochs.

Network performance on a manually-curated test image dataset consisting of 1,214 single-cell crops is shown as a confusion matrix in *Extended Data* Fig. 1i. All DAPI+ nuclei detected in patient samples were retrospectively classified by this apoptotic classifier CNN based on the BF and DAPI channels to quantify apoptotic fractions across the prospective patient cohort, marker-based subpopulations, GSC morphotypes, and drug conditions. Cells were classified as apoptotic (CASP3+) based on a CNN confidence threshold of 87%, close to the True Positive Rate (TPR) of the classifier.

### siRNA knockdown and quantitative real-time PCR

All siRNAs used in the study were part of the MISSION® esiRNA (Sigma-Aldrich, Euphoria Biotech) library (*Supplementary Table 5*) and ordered as custom gene arrays (esiOPEN, esiFLEX). FLUC esiRNA (EHUFLUC) targeting firefly Luciferase was used as a negative control, and KIF11 esiRNA (EHU019931) was used as a positive control for transfection and viability. For all siRNA experiments, low-passage LN-229 cells were used. siRNAs were transfected at 10ng/well in 384 well plates and 40ng/well in 96 well plates using Lipofectamine RNAiMAX (Invitrogen, #13778075). Imaging and drug incubation experiments were conducted in 384 well plates, while Incucyte live cell imaging and cell lysis preparation for RNA extraction and quantitative real-time PCR was performed in 96 well plates. For 384 well plates, both the siRNAs and Lipofectamine transfection reagent were dispensed using a Labcyte Echo liquid handler in a randomized plate layout to control for plate effects when possible. For data presented in *Extended Data* Fig. 5c,d*, and Extended Data* Fig. 8b, cells were incubated at 37°C, 5% CO_2_ for 48 hours following siRNA transfection before fixing, immunohistochemistry, and RNA extraction. For data presented in *Fig. 4j*, following 48 hours of siRNA transfection, cells were incubated for an additional 24 hours with either DMSO vehicle control or Vortioxetine (10µM) before fixing and subsequent analysis.

siRNA knockdown efficiency and relative abundance for the following target genes; BTG1, BTG2, JUN, and MKI67 was measured by TaqMan™ Array plates (Applied Biosystems, Standard, 96-well Plate; Format 16 with candidate endogenous controls) using the TaqMan™ Fast Advanced Master Mix (Thermo Scientific, #A44360) on a QuantStudio™ 3 Real-Time PCR System (Applied Biosystems, #A28567). Total RNA from LN-229 cells was extracted using the Direct-zol RNA MicroPrep Kit (Zymo Research, #R2062), RNA concentration was measured using the Qubit 4 Fluorometer (Thermo Scientific), and cDNA synthesized with the iScript™ cDNA Synthesis Kit (Bio-Rad, #1708890). For each TaqMan biological replicate assay (n=3 replicates) 25ng of cDNA per sample was used. To calculate the relative abundance of each target gene, the geometric mean Ct value of four endogenous control genes (18s rRNA, GAPDH, HPRT, GUSB) was subtracted from each [sample-target gene] Ct value to derive the deltaCt (dCt) value. Then, the mean deltaCt value from FLUC negative control samples was subtracted from each [sample-target gene] deltaCt value to derive the delta-deltaCt (ddCt) value. Finally, relative abundance (fold-difference) of each [sample-target gene] was calculated as the 2^(-ddCt).

### COSTAR: Convergence of secondary drug-targets analyzed by regularized regression

COSTAR is an interpretable molecular machine learning approach that utilizes logistic LASSO regression in a cross-validation setting to learn a multi-linear model that identifies the minimal set of drug-target connections that maximally discriminates PCY-hit drugs from PCY-negative drugs.

Drug-target connections were retrieved from the Drug Target Commons (DTC) ^50^. DTC is a crowd-sourced platform that integrates drug-target bioactivities curated from both literature and public databases such as PubChem and ChEMBL. Drug-target annotations (DTC bioactivities) listed as of August 2020 were included, with the target organism limited to *Homo sapiens*. Among PCY-tested drugs in our NAD and ONCD libraries, 127 out of 132 drugs had DTC ‘bioactivity’ annotations. PTGs with biochemical associations to a given drug correspond to bioactivities with the inhibitory constant ‘KI’ as the ‘End Point Standard Type’. Extended PTGs (ePTGs) include all annotated drug bioactivities. Secondary target genes (STGs) down-stream of ePTGs were retrieved by high-confidence protein-protein interactions annotated in the STRING database (interaction score≥0.6). The final drug-target connectivity map that was used for COSTAR consisted of 127 PCY-tested drugs, 975 extended primary targets, 10,573 secondary targets, and 114,517 network edges. The 127 drugs were labeled either as PCY-hits (n=30, equally split across NADs and ONCDs) or PCY-negative drugs (n=97) based on the ranked mean PCY score across patients.

A 20-fold cross-validated LASSO generalized linear model was trained in Matlab with the drug-target connectivity map as the predictor variable and PCY-hit status (hit vs. neg) as the binomially-distributed response variable to identify the optimal regularization coefficient (lambda) across a geometric sequence of 60 possible values. Final model coefficients were fitted using the lamba value corresponding to the minimum deviance in the cross-validation analysis shown in *Extended Data* Fig. 6a. COSTAR performance was first evaluated on the training dataset, represented as a confusion matrix in *Fig. 3h*. Using this trained linear model, COSTAR was next utilized as an *in silico* drug screening tool to predict the PCY-hit probability (COSTAR score) based on the connectivity of 1,120,823 compounds annotated in DTC (*Supplementary Table 6*). For interpretability, COSTAR subscores, defined as the individual connectivity to target genes multiplied by their respective coefficients (betas) in the linear model, can be investigated in *Extended Data* Fig. 6b,c. COSTAR predictions from this *in silico* screen were further experimentally validated in glioblastoma patient samples on a set of new drugs predicted as either COSTAR-hits or COSTAR-negs (n=48 drugs total; n=23 COSTAR-hits; n=25 COSTAR-negs).

### DRUG-Seq

High-throughput multiplexed RNA sequencing was performed with the Digital RNA with pertUrbation of Genes (DRUG-Seq) method as described in ^79^ with a few modifications. Modifications to the published method are the following: 1) extraction of RNA prior to cDNA reverse transcription with the Zymo Direct-zol-96 RNA isolation kit (Zymo, #R0256) 2) change of reverse transcription primers for compatibility with standard Illumina sequencing primers 3) cDNA clean-up prior to library amplification performed with the DNA Clean & Concentrator-5 kit (Zymo, #D4013) 4) tagmentation was performed with 2ng input and sequencing library generated using the Nextera XT library prep kit (Illumina, #FC-131-1024). In short, 1×10^4 LN-229 cells were plated in CellCarrier-96 Ultra Microplates (PerkinElmer, #6055302) and incubated overnight in reduced serum media at 37°C, 5% CO_2_ prior to drug treatment. A total of 20 drugs (*Supplementary Table 3*) were profiled across two different time-points (6 hours and 22 hours; n=4 replicates per drug and time-point). These 20 drugs were selected to include PCY-hit NADs spanning diverse drug classes (n=11), PCY-hit ONCDs (n=7), PCY-negative NADs (n=2), and a DMSO control. Cells in drug-treated 96-wells were lysed with TRIzol™ Reagent (ThermoFisher, #15596018) and then subsequent cDNA and library prep was performed as described above. 100bp (80:20) paired-end reads were generated using Illumina’s NextSeq 2000 platform.

### Calcium assays on the FLIPR platform

For calcium assays, 24 hours prior to the experiment, LN-229 cells were seeded at a density of 70,000 cells/well on poly-D-Lysine-coated ViewPlateTM-96 F TC 96-well black polystyrene clear bottom microplates (PerkinElmer, #6005182) in 100µl full medium. Calcium 6 dye stock solution was prepared by dissolving a vial from Calcium 6 assay kit (Molecular Devices, #5024048) in 10 ml sterile-filtered nominal Ca^2+^ free (NCF), modified Krebs buffer containing 117mM NaCl, 4.8mM KCl, 1mM MgCl2, 5mM D-glucose, 10mM HEPES (pH 7.4) and 500µl aliquots were stored at -20°C. Before each experiment, the dye stock was freshly diluted 1:10 in NCF Krebs buffer and after removing the medium from the cells, 50µl of the diluted dye was applied per well. In order to allow the cells to absorb the dye into their cytosol, they were incubated at 37°C for 2 hours in the dark. For assay setup outlined in *Fig. 4e*, cells were treated with their respective PCY-drug after a period of equilibration in 2mM calcium-containing buffer. The fluorescence Ca^2+^ measurements were carried out using FLIPR Tetra® (Molecular Devices) where cells were excited using a 470–495nm LED module and the emitted fluorescence signal was filtered with a 515–575nm emission filter according to the manufacturer’s guidelines. For fold change calculations presented in *Fig. 4f*, normalized calcium levels for each drug were calculated by averaging calcium levels after drug treatment (400-600 seconds interval) divided by the basal level of calcium prior to drug administration (200-300 seconds interval).

In the ER Ca^2+^ store release assay, the stable baselines were established for 50 seconds before 50µl of 2µM (2X) Thapsigargin (Sigma-Aldrich, #T9033) or 40µM (2X) drug solutions freshly prepared in NCF Krebs buffer were robotically dispensed to the cells to determine whether the drugs impact the ER Ca^2+^ stores. Next, the cells were incubated and fluorescence was monitored in the presence of Thapsigargin or drugs for another 5 min. In the extracellular Ca^2+^ uptake assay, after initial recording of the baseline, 50µl of 4mM CaCl2 (2X) prepared in NCF Krebs buffer was dispensed onto the cells to re-establish a physiological 2mM calcium concentration and the fluorescence was monitored for 5 min. Next, 60µM (3X) drug solutions freshly prepared in Krebs buffer containing 2mM CaCl_2_ were robotically dispensed to the cells and the fluorescence was recorded for an additional 4 min. The raw data was extracted with the ScreenWorks software version 3.2.0.14. The values represent average fluorescence level of the Calcium 6 dye measured over arbitrary selected and fixed time frames.

### Time-course RNA-Seq library preparation and sequencing

LN-229 cells were seeded at 2×10^5 cells/well into in 6-well Nunc™ Cell-Culture Treated Multidishes (ThermoFisher, #140675) and incubated overnight in reduced serum media at 37°C, 5% CO_2_ prior to drug treatment. The following day, Vortioxetine (Avachem Scientific, #3380) was manually added to each well at a final concentration of 20µM. At the start of the experiment, LN-229 cells that were not treated with Vortioxetine were collected as the 0 hour time-point. After 3, 6, 9, 12, and 24 hours following Vortioxetine treatment, drug-containing media was removed and cells were collected in TRIzol™ Reagent (ThermoFisher, #15596018). Total RNA was isolated using Direct-zol RNA MicroPrep Kit (Zymo Research, #R2062) and RNA quality and quantity was determined with the Agilent 4200 TapeStation. Sample RIN scores ranged from 5.9-10 (mean: 9.33). RNA input was normalized to 300-400 ng and RNA libraries were prepared using the Illumina Truseq stranded mRNA library prep. 100bp single-end reads were generated using Illumina’s Novaseq 6000 platform with an average sequencing depth of approximately 50 million reads per replicate. Reads were mapped and aligned to the reference human genome assembly (GRCh38.p13) using STAR/2.7.8a and counts were extracted using featureCounts. Subsequent read normalization (variance stabilizing transformation, vsd-normalized counts) and RNA-Seq analysis including differential expression (DE) analysis was performed with the R package ‘DESeq2’ ^87^.

### Time-course proteomics and phosphoproteomics

Cell preparation and Vortioxetine treatment was performed as in ‘Time-course RNA-Seq library preparation’ except cell numbers were scaled to be seeded in T-150 culture flasks and 3 time-points were measured (0, 3, 9 hours). Peptides for mass spectrometry measurements were prepared using the PreOmics iST kit (PreOmics) on the PreON (HSE AG). The robot was programmed to process 8 samples in parallel. During the first step of processing, cell pellets were resuspended in 50µl of lysis buffer and denatured for 10 minutes at 95°C. This step was followed by 3 hours of digestion with trypsin and Lys-C. Peptides were dried in a speed-vac (Thermo Fisher Scientific) for 1 hour before being resuspended in LC-Load buffer at a concentration of 1 μg/μl with iRT peptides (Biognosys).

Samples were analyzed on an Orbitrap Lumos mass spectrometer (Thermo Fisher Scientific) equipped with an Easy-nLC 1200 (Thermo Fisher Scientific). Peptides were separated on an in-house packed 30 cm RP-HPLC column (Michrom BioResources, 75 μm i.d. x 30 cm; Magic C18 AQ 1.9 μm, 200 Å). Mobile phase A consisted of HPLC-grade water with 0.1% formic acid, and mobile phase B consisted of HPLC-grade ACN (80%) with HPLC-grade water and 0.1% (v/v) formic acid. Peptides were eluted at a flow rate of 250 nl/min using a non-linear gradient from 4% to 47% mobile phase B in 228 min. For data-independent acquisition (DIA), DIA-overlapping windows were used and a mass range of m/z 396-1005 was covered. The DIA isolation window size was set to 8 and 4 m/z, respectively, and a total of 75 or 152 DIA scan windows were recorded at a resolution of 30,000 with an AGC target value set to 1200%. HCD fragmentation was set to 30% normalized collision. Full MS were recorded at a resolution of 60,000 with an AGC target set to standard and the maximum injection time set to auto. DIA data were analyzed using Spectronaut v14 (Biognosys). MS1 values were used for the quantification process, peptide quantity was set to mean. Data were filtered using Qvalue sparse with a precursor and a protein Qvalue cut-off of 0.01 FDR. Interference correction and local cross-run normalization was performed. For PRM measurements, peptides were separated by reversed-phase chromatography on a 50 cm ES803 C18 column (Thermo Fisher Scientific) that was connected to a Easy-nLC 1200 (Thermo Fisher Scientific). Peptides were eluted at a constant flow rate of 200 nl/min with a 117 min non-linear gradient from 4–52% buffer B (80% ACN, 0.1% FA) and 25-50%B. Mass spectra were acquired in PRM mode on an Q Exactive HF-X Hybrid Quadrupole-Orbitrap MS system (Thermo Fisher Scientific). The MS1 mass range was 340–1,400 m/z at a resolution of 120,000. Spectra were acquired at 60,000 resolution (automatic gain control target value 2.0*10e5); Normalized HCD collision energy was set to 28%, maximum injection time to 118 ms. Monitored peptides were analyzed in Skyline v20 and results were uploaded to PanoramaWeb. Targeted MS experiments can be accessed via Panorama (https://panoramaweb.org/GlioB.url). DIA and phosphopeptide enrichment datasets are available from MASSIVE under ftp://massive.ucsd.edu/MSV000090357/.

For phosphopeptide enrichment, protein lysate from LN-229 cells was prepared using a deoxycholate-based buffer. 500 μg of Vortioxetine-treated cells (time course of 0 mins, 30mins, 1h, 3h in triplicates) were used as starting material. A tryptic digest was performed for 16h. Samples were then purified on Macrospin C18 columns (Harvard Apparatus). Phosphopeptides were specifically enriched using IMAC cartridges on the Bravo AssayMAP liquid handling platform (Agilent). In short, samples were dissolved in 160 μl of loading buffer (80% ACN, 0.1% TFA). Then, the AssayMAP phosphoenrichment protocol was performed with slight modifications in terms of washing volume and speed. After purification, dried peptides were resuspended in LC buffer and subjected to DDA-MS on a QExactive H-FX mass spectrometer (Thermo Fisher Scientific) equipped with an Easy-nLC 1200 (Thermo Fisher Scientific). Peptides were separated on an ES903 column (Thermo Fisher Scientific, 75 μm i.d. x 50 cm; particle size 2 μm). Mobile phase A consisted of HPLC-grade water with 0.1% formic acid, and mobile phase B consisted of HPLC-grade ACN (80%) with HPLC-grade water and 0.1% (v/v) formic acid. Peptides were eluted at a flow rate of 250 nl/min using a non-linear gradient from 3% to 56% mobile phase B in 115 min. MS1 spectra were acquired at a resolution of 60,000 with an AGC target value of 3e6 and a maximum injection time of 56 ms. The scan range was between m/z 350-1650. A data-dependent top 12 method was used with a precursor isolation window of 1.3 m/z. MS/MS scans were acquired with normalized collision energy of 27 at a resolution of 15,000. AGC target was 1e5 with a maximum injection time of 22 ms. Dynamic exclusion was set to 30s. Data analysis was performed using FragPipe (v19.1) with the LFQ-phospho workflow ^88^. Min site localization probability was set to 0.75 in Ionquant^89^. Statistical analysis was performed on the phosphoprotein-filtered combined protein output in FragPipe-Analyst. Benjamini-Hochberg adjusted p-value cutoff was set to 0.05, log-fold change cutoff was 1. No imputation was selected.

### Incucyte live cell imaging

To measure cell proliferation in real-time, 2.5×10^3 LN-229 cells/well were plated in CellCarrier-96 Ultra Microplates (PerkinElmer, #6055302) 24 hours prior to the experiment, and transfected with BTG1, BTG2, and FLUC (-) MISSION® esiRNAs (Sigma-Aldrich, Euphoria Biotech, 40ng/well) using Lipofectamine RNAiMAX (Invitrogen, #13778075). Further details regarding siRNAs can be found in *Supplementary Table 5* and *Methods* related to ‘siRNA knockdown and quantitative real-time PCR**’.** Real-time confluence of cell cultures (n=4 replicate wells/condition) was monitored by imaging every 2 hours for 7 days at 10x magnification with the ‘phase’ channel using the Incucyte live-cell analysis system S3 (Sartorius). Automatic image segmentation and analysis of the phase contrast images was performed by the Incucyte base analysis software (version 2020B).

### Clonogenic survival assay

Adherent cells (LN-229: 50 cells; LN-308: 300 cells) were seeded in six replicates in 100 µL per well in 96-well plates and incubated overnight. On the following day, medium was replaced by fresh medium containing indicated final concentrations of Vortioxetine or DMSO. Glioblastoma-initiating cells (500 cells) were seeded in 75 µL medium and incubated overnight. Treatment was initiated by addition of 75 µL medium containing 2x concentrated Vortioxetine or DMSO to reach indicated final concentrations. DMSO concentration was kept at 0.5% for all treatments and controls. Following treatment addition, cells were cultured for 11 (LN-229) to 13 days (other cell lines) and clonogenic survival was estimated from a resazurin-based assay ^90^ using a Tecan M200 PRO plate reader (λEx = 560 nm / λEm = 590 nm).

### Collagen-based spheroid invasion assay

Spheroid invasion assay was performed as described (Kumar et al. 2015). Briefly, 2000 cells were seeded in six replicates into cell-repellent 96 U-bottom well plates (Greiner, #650979) and incubated for 48 hours to allow spheroid formation. Subsequently, 70 µl medium were removed, spheroids were overlaid with 70 µl 2.5% Collagen IV (Advanced Biomatrix, #5005-B) in 1xDMEM containing sodium bicarbonate (Sigma-Aldrich #S8761) and collagen was solidified in the incubator for 2 hours. Collagen-embedded spheroids were then overlaid with 100 µl chemoattractant (NIH-3T3-conditioned medium) containing 2x concentrated Vortioxetine/DMSO (0.5% final DMSO concentration across conditions) and incubated for 36 hours. Spheroids were stained with Hoechst and images were acquired on a MuviCyte imaging system (Perkin Elmer, #HH40000000) using a 4x objective. Images were contrast-enhanced and converted to binary using ImageJ/Fiji and quantified with the automated Spheroid Dissemination/Invasion counter software (aSDIcs), which quantifies the migration distance from the center of the spheroid for each detected cell nucleus ^91^.

#### *In vivo* drug testing

All animal experiments were done under the guidelines of the Swiss federal law on animal protection and were approved by the cantonal veterinary office (ZH98/2018). CD1 female nu/nu mice (Janvier, Le Genest-Saint-Isle, France) of 6 to 12 weeks of age were used in all experiments and 100’000 LN-229-derived- or 150’000 ZH-161-derived cells were implanted ^92^. Tumour-bearing mice were treated from day 5 – day 21 after tumour implantation with intraperitoneally (*i.p.*) administered Vortioxetine daily 10mg/kg, Citalopram daily 10mg/kg, Paliperidone daily 5mg/kg, Apomorphine daily 5mg/kg, Aprepitant daily 20mg/kg, Brexpiprazole daily 1mg/kg, Chlorpromazine three time per week 10mg/kg, Temozolomide 50mg/kg for five consecutive days, CCNU 20mg/kg at day 7 and 14 after tumour implantation, or daily DMSO control. Magnetic resonance imaging (MRI) was performed with a 4.7T imager (Bruker Biospin, Ettlingen, Germany) when the first mouse became symptomatic for *in vivo* Trials I-III or a 7T imager (Bruker BioSpin) at days 12, 25, 38 and 48 after tumor implantation for in vivo Trial IV. Coronal T2-weighted images were acquired using ParaVision 360(Bruker BioSpin). Tumor regions were identified manually by two independent raters and maximum perimeter was outlined and quantified using MIPAV (11.0.7).

Mouse brains were embedded in Shandon Cryochrome™ (Thermo Scientific) and were cut horizontally by 8μm steps until reaching the tumour. Sections were stained for 1 second with 0.4% methylene blue and rinsed with deionized water (2×10 dips) to confirm tumours (when present) under the microscope. Tissue sections were stored in the dark, in dry boxes overnight before being stored at -80°C. Tissue sections were subsequently fixed with 4% PFA (Sigma-Aldrich, #F8775) in PBS, blocked in 5% FBS and 0.1% Triton containing PBS, and stained overnight at 4°C in blocking solution with DAPI and the following antibodies and dilutions: Alexa Fluor® 488 anti-Vimentin (1:500, Biolegend, #677809, clone O91D3), anti-Ki67 (1:300, Cell Signaling Technology, #9129S, clone D3B5), goat anti-rabbit IgG (H+L) highly cross-adsorbed secondary antibody, Alexa Fluor Plus 647 (Thermo Scientific, #A32733). Imaging was performed by 20x fluorescence imaging using the Pannoramic 250 Slide Scanner (3DHISTECH).

## References

1. Ostrom, Q. T., Cioffi, G., Waite, K., Kruchko, C. & Barnholtz-Sloan, J. S. CBTRUS Statistical Report: Primary Brain and Other Central Nervous System Tumors Diagnosed in the United States in 2014-2018. Neuro. Oncol. 23, iii1–iii105 (2021).

2. Osswald, M. et al. Brain tumour cells interconnect to a functional and resistant network. Nature 528, 93–98 (2015).

3. Venkatesh, H. S. et al. Neuronal Activity Promotes Glioma Growth through Neuroligin-3 Secretion. Cell 161, 803–816 (2015).

4. Weil, S. et al. Tumor microtubes convey resistance to surgical lesions and chemotherapy in gliomas. Neuro. Oncol. 19, 1316–1326 (2017).

5. Venkataramani, V. et al. Glutamatergic synaptic input to glioma cells drives brain tumour progression. Nature 573, 532–538 (2019).

6. Venkatesh, H. S. et al. Electrical and synaptic integration of glioma into neural circuits. Nature 573, 539–545 (2019).

7. Alcantara Llaguno, S., et al. Cell-of-origin susceptibility to glioblastoma formation declines with neural lineage restriction. Nat. Neurosci. 22, 545–555 (2019).

8. Venkataramani, V. et al. Glioblastoma hijacks neuronal mechanisms for brain invasion. Cell 185, 2899–2917.e31 (2022).

9. Krishna, S. et al. Glioblastoma remodelling of human neural circuits decreases survival. Nature 617, 599–607 (2023).

10. Cancer Genome Atlas Research Network. Comprehensive genomic characterization defines human glioblastoma genes and core pathways. Nature 455, 1061–1068 (2008).

11. Brennan, C. W. et al. The somatic genomic landscape of glioblastoma. Cell 155, 462–477 (2013).

12. Suvà, M. L. et al. Reconstructing and reprogramming the tumor-propagating potential of glioblastoma stem-like cells. Cell 157, 580–594 (2014).

13. Neftel, C. et al. An Integrative Model of Cellular States, Plasticity, and Genetics for Glioblastoma. Cell 178, 835–849.e21 (2019).

14. Varn, F. S. et al. Glioma progression is shaped by genetic evolution and microenvironment interactions. Cell 185, 2184–2199.e16 (2022).

15. Stupp, R. et al. Radiotherapy plus concomitant and adjuvant temozolomide for glioblastoma. N. Engl. J. Med. 352, 987–996 (2005).

16. Hegi, M. E. et al. MGMT gene silencing and benefit from temozolomide in glioblastoma. N. Engl. J. Med. 352, 997–1003 (2005).

17. Weller, M. et al. EANO guidelines on the diagnosis and treatment of diffuse gliomas of adulthood. Nat. Rev. Clin. Oncol. 18, 170–186 (2021).

18. Singh, S. K. et al. Identification of human brain tumour initiating cells. Nature 432, 396–401 (2004).

19. Bao, S. et al. Glioma stem cells promote radioresistance by preferential activation of the DNA damage response. Nature 444, 756–760 (2006).

20. Chen, J. et al. A restricted cell population propagates glioblastoma growth after chemotherapy. Nature 488, 522–526 (2012).

21. Harder, B. G. et al. Developments in blood-brain barrier penetrance and drug repurposing for improved treatment of glioblastoma. Front. Oncol. 8, 462 (2018).

22. Le Rhun, E. et al. Molecular targeted therapy of glioblastoma. Cancer Treat. Rev. 80, 101896 (2019).

23. Gómez-Oliva, R. et al. Evolution of Experimental Models in the Study of Glioblastoma: Toward Finding Efficient Treatments. Front. Oncol. 10, 614295 (2020).

24. Patel, A. P. et al. Single-cell RNA-seq highlights intratumoral heterogeneity in primary glioblastoma. Science 344, 1396–1401 (2014).

25. John Lin, C.-C., et al. Identification of diverse astrocyte populations and their malignant analogs. Nat. Neurosci. 20, 396–405 (2017).

26. Lan, X. et al. Fate mapping of human glioblastoma reveals an invariant stem cell hierarchy. Nature 549, 227–232 (2017).

27. Lee, J. H. et al. Human glioblastoma arises from subventricular zone cells with low-level driver mutations. Nature 560, 243–247 (2018).

28. Couturier, C. P. et al. Single-cell RNA-seq reveals that glioblastoma recapitulates a normal neurodevelopmental hierarchy. Nat. Commun. 11, 3406 (2020).

29. Yu, K. et al. Surveying brain tumor heterogeneity by single-cell RNA-sequencing of multi-sector biopsies. Natl Sci Rev 7, 1306–1318 (2020).

30. Dolma, S. et al. Inhibition of Dopamine Receptor D4 Impedes Autophagic Flux, Proliferation, and Survival of Glioblastoma Stem Cells. Cancer Cell 29, 859–873 (2016).

31. Caragher, S. P. et al. Activation of Dopamine Receptor 2 Prompts Transcriptomic and Metabolic Plasticity in Glioblastoma. J. Neurosci. 39, 1982–1993 (2019).

32. Bi, J. et al. Targeting glioblastoma signaling and metabolism with a re-purposed brain-penetrant drug. Cell Rep. 37, 109957 (2021).

33. Snijder, B. et al. Image-based ex-vivo drug screening for patients with aggressive haematological malignancies: interim results from a single-arm, open-label, pilot study. Lancet Haematol 4, e595–e606 (2017).

34. Kornauth, C. et al. Functional Precision Medicine Provides Clinical Benefit in Advanced Aggressive Hematologic Cancers and Identifies Exceptional Responders. Cancer Discov. 12, 372– 387 (2022).

35. Heinemann, T. et al. Deep morphology learning enhances ex vivo drug profiling-based precision medicine. Blood Cancer Discov (2022) doi:10.1158/2643-3230.BCD-21-0219.

36. Sheng, M. & Greenberg, M. E. The regulation and function of c-fos and other immediate early genes in the nervous system. Neuron 4, 477–485 (1990).

37. Rouault, J. P. et al. Identification of BTG2, an antiproliferative p53-dependent component of the DNA damage cellular response pathway. Nat. Genet. 14, 482–486 (1996).

38. Yap, E.-L. & Greenberg, M. E. Activity-Regulated Transcription: Bridging the Gap between Neural Activity and Behavior. Neuron 100, 330–348 (2018).

39. Yuniati, L., Scheijen, B., van der Meer, L. T. & van Leeuwen, F. N. Tumor suppressors BTG1 and BTG2: Beyond growth control. J. Cell. Physiol. 234, 5379–5389 (2019).

40. Malta, T. M. et al. Machine Learning Identifies Stemness Features Associated with Oncogenic Dedifferentiation. Cell 173, 338–354.e15 (2018).

41. Jacob, F. et al. A Patient-Derived Glioblastoma Organoid Model and Biobank Recapitulates Inter- and Intra-tumoral Heterogeneity. Cell 180, 188–204.e22 (2020).

42. Dahlstrand, J., Collins, V. P. & Lendahl, U. Expression of the Class VI Intermediate Filament Nestin in Human Central Nervous System Tumors1. Cancer Res. 52, 5334–5341 (1992).

43. Yuan, X. et al. Isolation of cancer stem cells from adult glioblastoma multiforme. Oncogene 23, 9392–9400 (2004).

44. Zhang, M. et al. Nestin and CD133: valuable stem cell-specific markers for determining clinical outcome of glioma patients. J. Exp. Clin. Cancer Res. 27, 85 (2008).

45. Schmid, J. A. et al. Efficacy and feasibility of Pharmacoscopy-guided treatment for acute myeloid leukemia patients that exhausted all registered therapeutic options. medRxiv (2023) doi:10.1101/2023.03.28.23287745.

46. Kropivsek, K. et al. Ex vivo drug response heterogeneity reveals personalized therapeutic strategies for patients with multiple myeloma. Nat Cancer 4, 734–753 (2023).

47. Wick, W. et al. Phase II Study of Radiotherapy and Temsirolimus versus Radiochemotherapy with Temozolomide in Patients with Newly Diagnosed Glioblastoma without MGMT Promoter Hypermethylation (EORTC 26082). Clin. Cancer Res. 22, 4797–4806 (2016).

48. Chinnaiyan, P. et al. A randomized phase II study of everolimus in combination with chemoradiation in newly diagnosed glioblastoma: results of NRG Oncology RTOG 0913. Neuro. Oncol. 20, 666–673 (2018).

49. Lombardi, G. et al. Regorafenib compared with lomustine in patients with relapsed glioblastoma (REGOMA): a multicentre, open-label, randomised, controlled, phase 2 trial. Lancet Oncol. 20, 110–119 (2019).

50. Tang, J. et al. Drug Target Commons: A Community Effort to Build a Consensus Knowledge Base for Drug-Target Interactions. Cell Chem Biol 25, 224–229.e2 (2018).

51. Hai, T. & Curran, T. Cross-family dimerization of transcription factors Fos/Jun and ATF/CREB alters DNA binding specificity. Proc. Natl. Acad. Sci. U. S. A. 88, 3720–3724 (1991).

52. Wang, L. et al. A single-cell atlas of glioblastoma evolution under therapy reveals cell-intrinsic and cell-extrinsic therapeutic targets. Nat Cancer 3, 1534–1552 (2022).

53. Karin, M., Liu, Z. g. & Zandi, E. AP-1 function and regulation. Curr. Opin. Cell Biol. 9, 240–246 (1997).

54. Sanyal, S., Sandstrom, D. J., Hoeffer, C. A. & Ramaswami, M. AP-1 functions upstream of CREB to control synaptic plasticity in Drosophila. Nature 416, 870–874 (2002).

55. Carlsson, P. & Mahlapuu, M. Forkhead Transcription Factors: Key Players in Development and Metabolism. Dev. Biol. 250, 1–23 (2002).

56. Ho, K. K., Myatt, S. S. & Lam, E. W.-F. Many forks in the path: cycling with FoxO. Oncogene 27, 2300–2311 (2008).

57. Benayoun, B. A., Caburet, S. & Veitia, R. A. Forkhead transcription factors: key players in health and disease. Trends Genet. 27, 224–232 (2011).

58. Sheng, M., Thompson, M. A. & Greenberg, M. E. CREB: a Ca(2+)-regulated transcription factor phosphorylated by calmodulin-dependent kinases. Science 252, 1427–1430 (1991).

59. Whitmarsh, A. J. & Davis, R. J. Transcription factor AP-1 regulation by mitogen-activated protein kinase signal transduction pathways. J. Mol. Med. 74, 589–607 (1996).

60. Park, C. Y. et al. Tissue-aware data integration approach for the inference of pathway interactions in metazoan organisms. Bioinformatics 31, 1093–1101 (2015).

61. Grünblatt, E., Mandel, S., Berkuzki, T. & Youdim, M. B. Apomorphine protects against MPTP-induced neurotoxicity in mice. Mov. Disord. 14, 612–618 (1999).

62. Guilloux, J.-P. et al. Antidepressant and anxiolytic potential of the multimodal antidepressant vortioxetine (Lu AA21004) assessed by behavioural and neurogenesis outcomes in mice. Neuropharmacology 73, 147–159 (2013).

63. Maeda, K. et al. Brexpiprazole I: in vitro and in vivo characterization of a novel serotonin-dopamine activity modulator. J. Pharmacol. Exp. Ther. 350, 589–604 (2014).

64. Berger, M. et al. Hepatoblastoma cells express truncated neurokinin-1 receptor and can be growth inhibited by aprepitant in vitro and in vivo. J. Hepatol. 60, 985–994 (2014).

65. Oliva, C. R., Zhang, W., Langford, C., Suto, M. J. & Griguer, C. E. Repositioning chlorpromazine for treating chemoresistant glioma through the inhibition of cytochrome c oxidase bearing the COX4-1 regulatory subunit. Oncotarget 8, 37568–37583 (2017).

66. Torrisi, S. A. et al. Fluoxetine and Vortioxetine Reverse Depressive-Like Phenotype and Memory Deficits Induced by Aβ1-42 Oligomers in Mice: A Key Role of Transforming Growth Factor-β1. Front. Pharmacol. 10, 693 (2019).

67. Lee, J.-K., Nam, D.-H. & Lee, J. Repurposing antipsychotics as glioblastoma therapeutics: Potentials and challenges. Oncol. Lett. 11, 1281–1286 (2016).

68. Tan, S. K. et al. Drug Repositioning in Glioblastoma: A Pathway Perspective. Front. Pharmacol. 9, 218 (2018).

69. Caragher, S. P., Hall, R. R., Ahsan, R. & Ahmed, A. U. Monoamines in glioblastoma: complex biology with therapeutic potential. Neuro. Oncol. 20, 1014–1025 (2018).

70. Lee, J.-K. et al. Spatiotemporal genomic architecture informs precision oncology in glioblastoma. Nat. Genet. 49, 594–599 (2017).

71. Lee, J.-K. et al. Pharmacogenomic landscape of patient-derived tumor cells informs precision oncology therapy. Nat. Genet. 50, 1399–1411 (2018).

72. MacLeod, G. et al. Genome-Wide CRISPR-Cas9 Screens Expose Genetic Vulnerabilities and Mechanisms of Temozolomide Sensitivity in Glioblastoma Stem Cells. Cell Rep. 27, 971–986.e9 (2019).

73. Stockslager, M. A. et al. Functional drug susceptibility testing using single-cell mass predicts treatment outcome in patient-derived cancer neurosphere models. Cell Rep. 37, 109788 (2021).

74. Shekarian, T. et al. Immunotherapy of glioblastoma explants induces interferon-γ responses and spatial immune cell rearrangements in tumor center, but not periphery. Sci Adv 8, eabn9440 (2022).

75. Vladimer, G. I. et al. Global survey of the immunomodulatory potential of common drugs. Nat. Chem. Biol. 13, 681–690 (2017).

76. Shilts, J. et al. A physical wiring diagram for the human immune system. Nature (2022) doi:10.1038/s41586-022-05028-x.

77. Meister, H. et al. Multifunctional mRNA-Based CAR T Cells Display Promising Antitumor Activity Against Glioblastoma. Clin. Cancer Res. 28, 4747–4756 (2022).

78. Eferl, R. & Wagner, E. F. AP-1: a double-edged sword in tumorigenesis. Nat. Rev. Cancer 3, 859– 868 (2003).

79. Ye, C. et al. DRUG-seq for miniaturized high-throughput transcriptome profiling in drug discovery. Nat. Commun. 9, 4307 (2018).

80. Li, J. et al. DRUG-seq Provides Unbiased Biological Activity Readouts for Neuroscience Drug Discovery. ACS Chem. Biol. 17, 1401–1414 (2022).

81. Schaefer, C. F. et al. PID: the Pathway Interaction Database. Nucleic Acids Res. 37, D674–9 (2009).

82. Landrum, M. J. et al. ClinVar: improving access to variant interpretations and supporting evidence. Nucleic Acids Res. 46, D1062–D1067 (2018).

83. Dai, Y. et al. WebCSEA: web-based cell-type-specific enrichment analysis of genes. Nucleic Acids Res. 50, W782–W790 (2022).

84. Trapnell, C. et al. The dynamics and regulators of cell fate decisions are revealed by pseudotemporal ordering of single cells. Nat. Biotechnol. 32, 381–386 (2014).

85. Miao, Z., Deng, K., Wang, X. & Zhang, X. DEsingle for detecting three types of differential expression in single-cell RNA-seq data. Bioinformatics 34, 3223–3224 (2018).

86. Krizhevsky, A., Sutskever, I. & Hinton, G. E. ImageNet classification with deep convolutional neural networks. in Proceedings of the 25th International Conference on Neural Information Processing Systems - Volume 1 1097–1105 (Curran Associates Inc., 2012).

87. Love, M. I., Huber, W. & Anders, S. Moderated estimation of fold change and dispersion for RNA-seq data with DESeq2. Genome Biol. 15, 550 (2014).

88. Kong, A. T., Leprevost, F. V., Avtonomov, D. M., Mellacheruvu, D. & Nesvizhskii, A. I. MSFragger: ultrafast and comprehensive peptide identification in mass spectrometry–based proteomics. Nat. Methods 14, 513–520 (2017).

89. Yu, F., Haynes, S. E. & Nesvizhskii, A. I. IonQuant Enables Accurate and Sensitive Label-Free Quantification With FDR-Controlled Match-Between-Runs. Mol. Cell. Proteomics 20, 100077 (2021).

90. Riss, T. L. et al. Cell Viability Assays. in Assay Guidance Manual (eds. Markossian, S. et al.) (Eli Lilly & Company and the National Center for AdvancingTranslational Sciences, 2013).

91. Kumar, K. S. et al. Computer-assisted quantification of motile and invasive capabilities of cancer cells. Sci. Rep. 5, 15338 (2015).

92. Weiss, T. et al. NKG2D-Dependent Antitumor Effects of Chemotherapy and Radiotherapy against Glioblastoma. Clin. Cancer Res. 24, 882–895 (2018).

